# Prenatal liver is resistant to low glucose-induced inhibition of mTORC1

**DOI:** 10.1101/2025.07.21.665892

**Authors:** Chen-Jie Zhang, Chuan-Jin Yu, Yi Cheng, Jie-Xue Pan, Long-Yun Ye, Yun-Hui Tang, Xue-Yun Qin, Zhong-Liang Lin, Chen-Song Zhang, Guo-Lian Ding, Sheng-Cai Lin, He-Feng Huang

## Abstract

Prenatal development may adopt unique patterns of metabolic responses to different nutrient supplies. AMPK and mTOR are two central metabolic regulators, with AMPK acting to slow down and mTOR to promote anabolism. Typically, low glucose inhibits TRPVs, leading to AMPK activation and mTOR complex I (mTORC1) inhibition. Here, we observed surprisingly that mTORC1 in the foetal liver remains active in low glucose, although AMPK is effectively activated in an AMP-dependent manner. Mechanistically, we found that TRPV4, a key liver TRPV, is acetylated at K608, disabling its inactivation by fructose-1,6-bisphosphate (FBP)-unoccupied aldolase. Mutation of K608 to arginine renders TRPV4 able to respond to low glucose in foetal hepatocytes. Consistently, embryonic liver-specific expression of acetylation-defective TRPV4-K608R mutant led to mTORC1 inhibition, and caused abnormal intrauterine development or even foetal death. Our findings reveal that the TRPV-mediated resistance of mTORC1 inhibition in low glucose acts as a safeguard for normal foetal development.

## INTRODUCTION

The interplay between mTORC1 and AMPK is crucial in maintaining metabolic homeostasis within an organism in response to fluctuating glucose availability^1^. In ample glucose, mTORC1 is active, and promotes anabolism^2–4^. Conversely, during periods of glucose and/or energy deficiency, AMPK is activated, which promotes catabolism and inhibits anabolism^5–7^. It has been shown that low glucose can autonomously dissociate mTORC1 from the lysosomal surface, thereby inhibiting mTORC1^8^. Meanwhile, the activated AMPK can further inhibit mTORC1^9^.

The low glucose-induced inhibition of mTORC1 takes place on the surface of the lysosome, through the glucose-sensing pathway, in which the vacuolar H+-ATPases (v-ATPase), a proton pump, plays a pivotal role^10,11^. It was shown that the v-ATPase-associated glycolytic enzyme aldolase, the enzyme that cleaves the glycolytic intermediate fructose-1,6-bisphosphate (FBP) to dihydroxyacetone phosphate (DHAP) and glyceraldehyde-3-phosphate (G3P), is the sensor of glucose^12,13^. When glucose levels are low, an increased proportion of the v-ATPase-associated aldolase becomes unoccupied with FBP/DHAP, which in turn blocks the endoplasmic reticulum (ER)-localized transient receptor potential V (TRPV) calcium channels, converting the low cellular glucose signal to a low calcium signal at the ER-lysosome contact^10,13,14^. In low local Ca^2+^, TRPVs then interact also with v-ATPase and likely reconfigure the aldolase-v-ATPase complex, resulting in inhibition of the v-ATPase^10^. As a result, the intrinsically disordered scaffold protein AXIN, in complex with LKB1, translocates to the lysosomal surface by utilising v-ATPase and its associated Ragulator (comprised of 5 LAMTOR subunits, LAMTOR1-5) as docking sites^11,15^. On these interactions, AXIN promotes the conformational changes of Ragulator, disabling its ability to release GTP from RAGC [one component of the small GTPase RAGs (RAGA to RAGD)]^11,16^, and hence triggering the switch of RAG heterodimers from an “on” state (RAGA^GTP^-RAGC^GDP^) to an “off” state (RAGA^GDP^-RAGC^GTP^)^17^. Consequently, mTORC1 is dissociated from the Ragulator-RAGs complex^18^, and hence also detached from its allosteric activator, the lysosomal pool of Rheb GTPase^19,20^. It is noteworthy that the dissociation of mTORC1 from the lysosome is autonomous, as it can still be inhibited in the absence of AMPK^8,21^, albeit at a slightly slower rate^11,22,23^. Meanwhile, upon translocation onto the lysosomal surface, the AXIN-carried LKB1, an upstream kinase, phosphorylates and activates the lysosomal pool of AMPK^11^. In addition to the abovementioned inhibition of mTORC1 as the result of dissociation from the lysosomal surface, the activated AMPK can phosphorylate Raptor^24^, an essential component of mTORC1^25^, or phosphorylate TSC2 to inhibit Rheb^9^, providing additional inhibitory mechanisms for mTORC1 in response to low glucose. In addition to glucose, amino acids and growth factors also play critical roles in maintaining the activity of mTORC1. For example, high leucine and arginine can activate RAGs via the v-ATPase-Ragulator complex^17,26–30^. In addition, leucine and arginine can activate mTORC1 through the GATOR1/2 complex that turns on RAGs^31–33^. Apart from RAGs, it is also shown that amino acids could inhibit TSC2 or elevate intracellular Ca^2+^ that activates Rheb^34–36^, or induce the ubiquitination of mTOR itself that allosterically activates mTORC1^37^. In parallel, growth factors activate Rheb through the PI3K-AKT pathway that inhibits _TSC238-40._

It is known that glucose scarcity occurs during foetal development^41^. However, despite the low glucose or maternal hypoglycaemia, the foetus grows rapidly, at a normal growth rate, which demands active anabolism^41–43^. For example, in rats, under maternal postprandial conditions, foetal blood glucose levels are only around 3 mM^42^. When the mother undergoes a 24-h fasting, foetal blood glucose levels can decrease to 2 mM and below^42^. On the other hand, mTORC1 activity has also been shown to be required for normal foetal development^44,45^. Therefore, how mTORC1 activity is maintained under the hypoglycaemia conditions during foetal development needs to be explored.

## RESULTS

### mTORC1 is active in foetal hepatocytes in low glucose

We subjected BNL-Cl2 cells, a cell line derived from mouse foetal liver, to treatment with different glucose concentrations. In low glucose (< 5 mM), we found that BNL-Cl2 cells did not show inhibition of mTORC1 activity despite significant activation of AMPK (Figure 1A), as evidenced by the phosphorylation of S6K and the phosphorylation of AMPKα, respectively. As a control, AML12 cells (derived from adult mouse liver), HEK293T cells and MEFs all exhibited rapid inhibition of mTORC1 when cultured in low glucose (Figure 1A). We also tested the effects of low glucose over different time durations, showing that prolonged glucose starvation still did not inhibit mTORC1 in the BNL-CL2 cells (Figure 1C). Moreover, in primary mouse hepatocytes derived from the foetal liver (PFH), we found that activity of mTORC1 also remained active in low glucose (Figure 1B, D). These results indicate that low glucose cannot inhibit mTORC1 in cells derived from the foetal liver.

**Figure 1.**
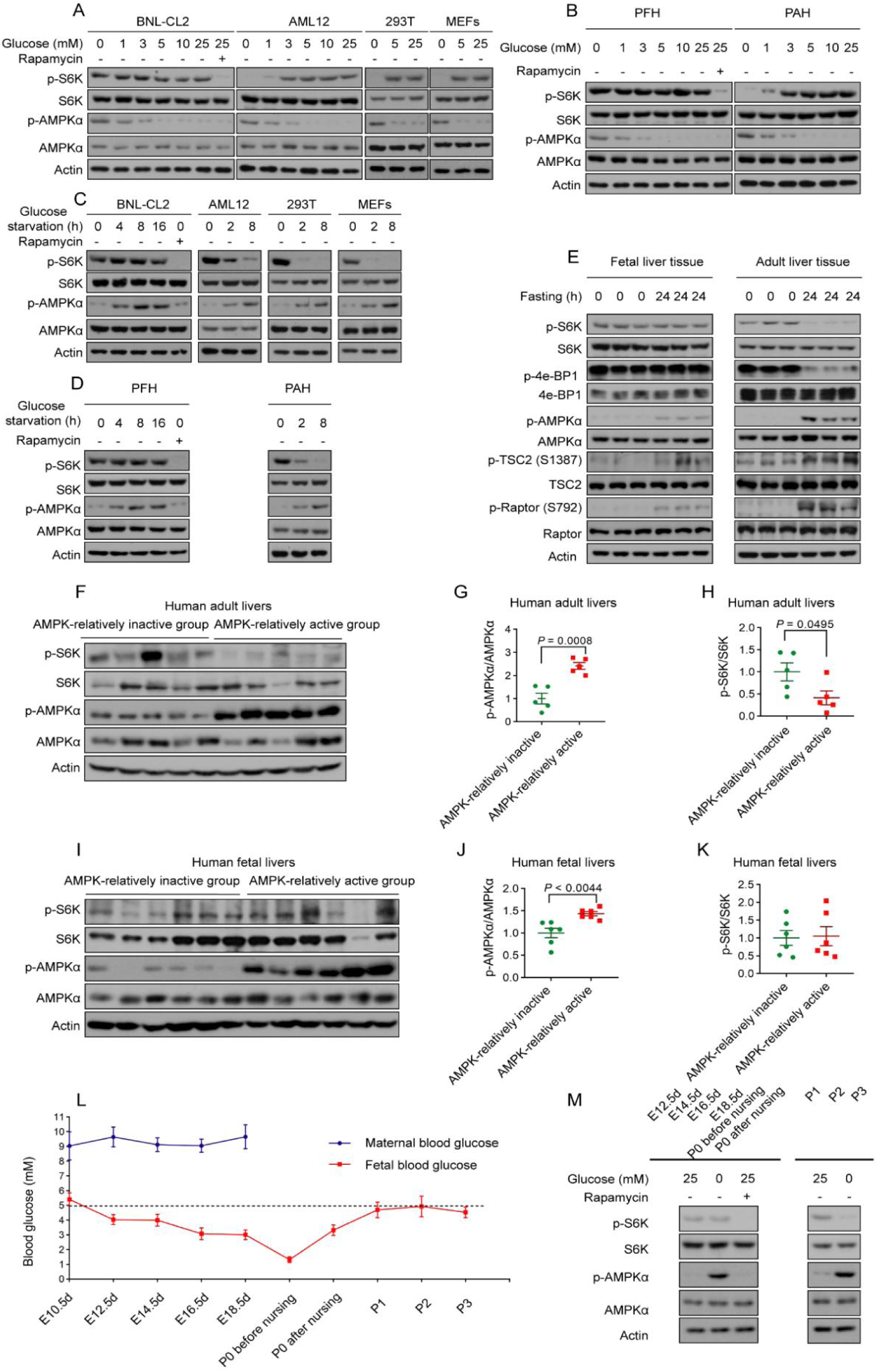
mTORC1 remains active in foetal hepatocytes in low glucose. A-D. mTORC1 remains active in foetal hepatocytes when cultured in low glucose. The foetal hepatocyte cell line BNL-CL2 (A, C), primary foetal hepatocytes (PFH; B, D), the adult hepatocyte cell line AML12 (A, C), and primary adult hepatocytes (PAH; A, C), along with HEK293T cells and MEFs as controls (A, C), were incubated in different concentrations of glucose for 4 h (A-D, BNL-CL2, PFH), 2 h (A-C, others), or other desired time periods (C, D), followed by determination of phosphorylation of S6K (p-S6K). As a control, rapamycin (100 nM, treatment for 1 h) was used to inhibit S6K phosphorylation. See also phosphorylation of AMPKα in these cells. E. mTORC1 is active in foetal livers after starvation. Liver tissues collected from pregnant mice embryos (E18.5d; foetal liver tissue) and adult mice (8-week-old; adult liver tissue), after fasting for 24 h, were subjected to determination of p-S6K by immunoblotting. F-K. The relationship between AMPK activation and mTORC1 activation does not show a negative correlation in foetal human livers. Liver tissues collected from embryos during first-trimester pregnancy termination surgeries (F-H), or from adult patients (I-K), all of whom had fasted before the surgery, were subjected to determination of the levels of p-S6K and p-AMPKα. Based on the levels of p-AMPKα, the samples were categorized into two groups: AMPK-relatively inactive and AMPK-relatively active. The ratios of p-S6K/S6K and p-AMPKα/AMPKαin each group, as well as the correlation between them, are shown in graphs G, H, J, and K as mean ± SEM, with n = 5, and P values by unpaired two-tailed Student’s *t*-test. L. During pregnancy and parturition, the blood glucose levels of foetal mice are much lower than those of female mice. Blood glucose levels in pregnant (n = 5) and foetal (n = 10) mice at various developmental stages were measured, and the data are shown as mean ± SEM. M. Mouse foetal liver mTORC1 cannot be inhibited by low glucose until the day of birth. Primary foetal hepatocytes isolated from mice at various developmental stages were incubated in low glucose for 4 h, followed by determination of the levels of p-S6K and p-AMPKα. Experiments in this figure were performed three times.

At the organismal level, we fasted pregnant mice (foetuses of E18.5), along with and unpregnant female adult mice, for 24 h. This resulted in a significant decrease of plasma glucose (from 3.5 to 2.1 mM in the foetus; from 9.3 to 4.4 mM in the adults, Figure S8H), and activation of hepatic AMPK in both foetus and adults (Figure 1E; note that the level of AMPK activation in foetal liver was much higher than that in adult liver, as shown in Figure S1A). We also observed active mTORC1 in the foetal liver, and inhibition of mTORC1 in the adult liver (Figure 1E). In human liver samples taken from adult patients, we found strongly negative correlation between AMPK activation and mTORC1 activation (Figure 1F-H). In comparison, we did not observe such a correlation in human foetal livers, where higher AMPK activation did not correspond to lower mTORC1 activation (Figure 1I-K).

Overall, these data indicate that low glucose is unable to inhibit mTORC1 in foetal hepatocytes or in the foetal liver, even when AMPK is activated allosterically activated by increased levels of AMP, as detailed below. However, the resistance of mTORC1 to inhibition by low glucose occurs only in the foetal liver tissue, but not in the other tissues. For example, in the foetal heart, lung, and muscle tissues, mTORC1 was significantly inhibited after 24 h of maternal fasting (Figure S1B-D). Interestingly, this resistance to inhibition occurred only during the perinatal state and disappeared as soon as at postnatal day 1 (Figure 1L, M).

### Inability to inhibit the v-ATPase-Ragulator-RAG complex underlies maintenance of mTORC1 in foetal hepatocytes

We next investigated the mechanisms through which the low glucose-induced inhibition of mTORC1 is blocked in the foetal liver. We examined the phosphorylation of TSC2 and Raptor, substrates of AMPK, which mediate the inhibition of mTORC1 by AMPK. As shown in Figure 1e, we found intact phosphorylation of TSC2 (at S1387)^9^ and Raptor (S792)^24^ in the fasted foetal liver tissue. We also investigated whether a lack of glucose could dissociate mTORC1 from the lysosomal membrane as consequence of detaching from RAG and Ragulator^16,17^. We immunoprecipitated RagA, and observed that mTORC1 remained associated with RAG in foetal hepatocytes in low glucose (Figure S1E). In contrast, mTORC1 was not associated with RAG in adult hepatocytes (Figure S1E). Immunofluorescent staining further showed that mTORC1 remained co-localised with the lysosomal marker LAMP2 in foetal hepatocytes in low glucose, but not in adult hepatocytes (Figure 2A-D and Figure S1F-I). We also determined the activity of v-ATPase, assessed by the fluorescent intensity of Lysosensor dye that positively correlates with the lysosomal acidity^46^. We observed that glucose starvation did not affect the lysosomal acidity in foetal hepatocytes (Figure S1J-L), indicating that v-ATPase remains active. In contrast, low glucose effectively inhibited v-ATPase in adult hepatocytes (Figure S1M-O). As a control, forced inhibition of v-ATPase in foetal hepatocytes by concanamycin A (conA) effectively triggered the lysosomal translocation of AXIN (Figure S2A-B), disrupted the lysosomal localisation of mTORC1 (Figure 2A-D) and led to its inhibition (Figure 2E). The results above indicate that the mTORC1 activity in the foetal liver is maintained due to the unaffected v-ATPase-Ragulator-RAG axis in low glucose.

**Figure 2.**
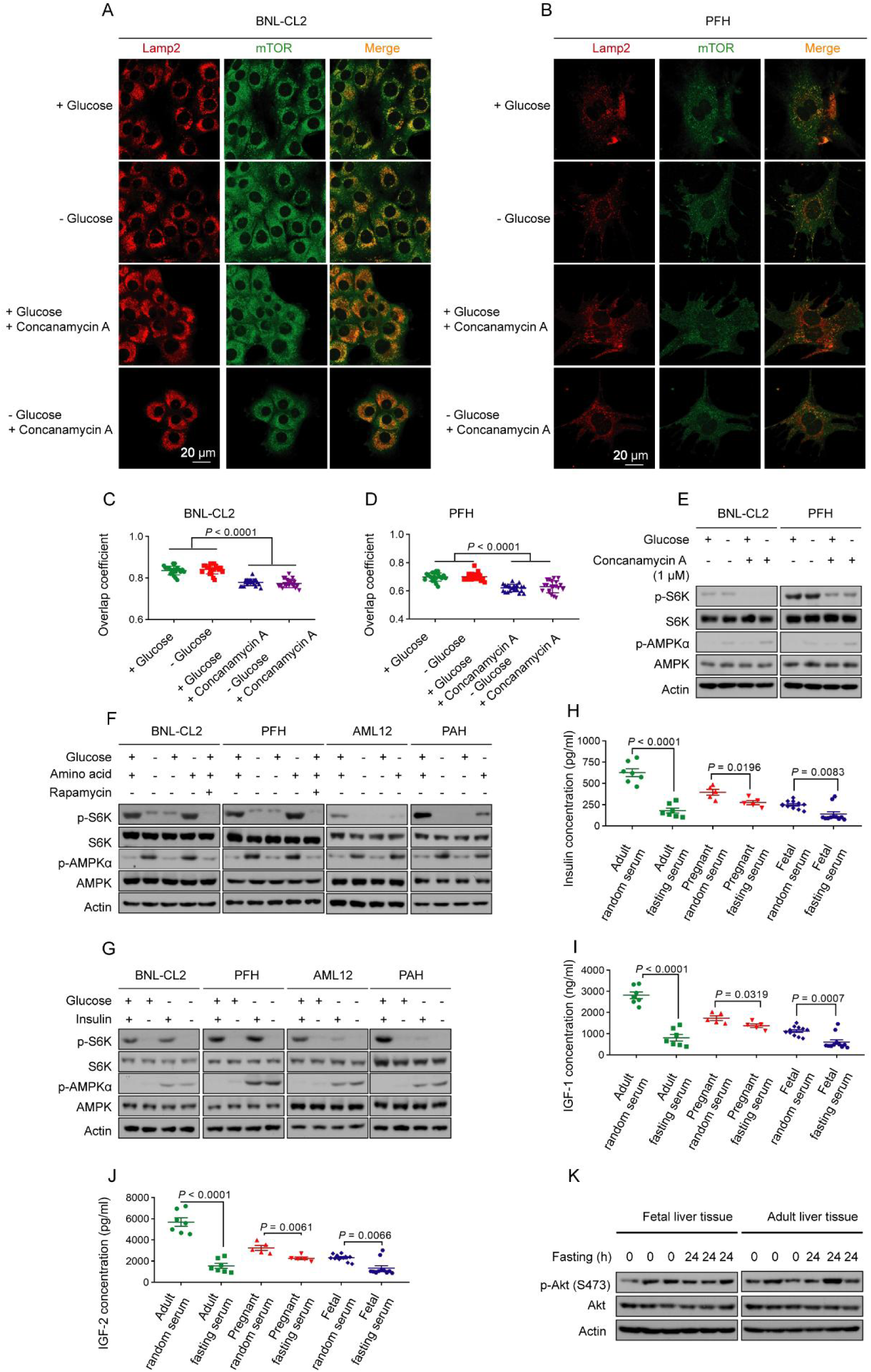
Constitutively active v-ATPase-Ragulator-RAG axis maintains the mTORC1 activity in foetal hepatocytes in low glucose. A-D. mTORC1 remains localized on lysosomes in low glucose in foetal hepatocytes. The BNL-CL2 foetal hepatocyte cell line (A, C) and primary foetal hepatocytes (PFH; B, D) were either glucose-starved or treated with 1 μM concanamycin A, the v-ATPase inhibitor, as a control for 6 h. Endogenous mTOR (labeled in green with rabbit anti-mTOR antibody, CST 2983T) and the lysosome marker LAMP2 (labeled in red with rat anti-LAMP2 antibody, Abcam AB13524) were then stained. Representative images are shown in A and B, and the Mander’s overlap coefficients in C and D (presented as mean ± SEM, n = 15 (C) or 8 (D) for each group, with P values calculated using unpaired two-tailed Student’s *t*-test). E. Forced inhibition of v-ATPase effectively inhibits mTORC1 in foetal hepatocytes. The BNL-CL2 cells and primary foetal hepatocytes were treated with 1 μM concanamycin A for 6 h, followed by determination of p-S6K by immunoblotting. F, G. mTORC1 in foetal hepatocytes is sensitive to deprivation of amino acids or serum. Both BNL-CL2 cells and primary foetal hepatocytes, as well as adult hepatocytes as control, were deprived of serum overnight and then re-stimulated with 5 μg/ml insulin for 30 minutes, deprived of amino acids for 2 h, or starved of glucose for 2 h. Cells were then lysed, followed by determination of p-S6K by immunoblotting. H-K. Maternal fasting does not inhibit AKT signaling in foetal livers. The pregnant mice were fasted for 24 h, followed by determination of the p-AKT levels in the liver of their foetuses (E18.5d; foetal liver tissue) by immunoblotting (K), and the insulin (H), IGF-1 (I), and IGF-2 (J) concentrations in the serum of the foetuses by ELISA. See also insulin, IGF-1, and IGF-2 levels in the serum of pregnant mice, as well as p-AKT levels in the liver of adult mice (8-week-old), as controls. Data in H-J are shown as mean ± SEM; n = 7 (adult mice of H-J), 5 (pregnant mice of H-J), or 11 (foetal mice of H-J) mice for each condition, and P values by unpaired two-tailed Student’s *t*-test, except for some foetal mice (H, J) where unpaired two-tailed Mann-Whitney test is used. Experiments in this figure were performed three times, except H-J one time.

We also examined the effects of amino acids and growth factors on mTORC1 in foetal liver cells, and found that mTORC1 is readily inhibited in these cells upon the withdrawal of amino acids or growth factors (serum), as evidenced by decreased p-S6K (Figure 2F, G) and lysosomal localisation of mTORC1 (Figure S2C-F). Therefore, mTORC1 in foetal hepatocytes is resistant specifically to glucose withdrawal, but not to the removal of amino acids or growth factors. Consistent with the constitutive activity of mTORC1, maternal fasting did not decrease the levels of amino acids, particularly leucine, arginine, and glutamine, each of which could independently keep mTORC1 in an active state in the presence of both growth factors and glucose^47,48^ (Figure S2G-J; see also the contents of other amino acids in Figure S3A-Q). Similarly, maternal fasting did not inhibit the PI3K-AKT pathway in the foetal liver (Figure 2K), although it did lower the serum levels of insulin, IGF-1, and IGF-2 (Figure 2H-J).

### Low glucose-triggered lysosomal AMPK pathway is blocked in the foetal liver

We next investigated why the v-ATPase-Ragulator-RAG axis in the foetal liver is not inhibited by low glucose. In adult liver or hepatocytes, low glucose renders the v-ATPase-associated aldolase unoccupied with FBP, triggering inhibition of v-ATPase and consequentially the inhibition of the Ragulator-RAG complex^10,13^. We thus determined the levels of FBP and DHAP in foetal hepatocytes in low glucose, and observed a strong decrease in FBP and DHAP levels in both foetal liver cell lines and primary foetal hepatocytes in low glucose (Figure S4A-L; see also the levels of other glycolytic intermediates in Figure S5 and S6). Such a decrease could also be observed in the foetal liver after maternal fasting (Figure S4M-O), similarly to the adult mouse livers (Figure S4P-R; the levels of other glycolytic intermediates are shown in Figure S7). Therefore, the maintenance of foetal liver mTORC1 activity in low glucose is not due to sustained intracellular glucose, FBP, or DHAP.

We also investigated whether aldolase and TRPV are affected by low glucose. We found that in foetal hepatocytes, all three isozymes of aldolase (ALDOA to ALDOC) could bind to TRPV (Figure 3A; represented by TRPV4, as it had the highest mRNA levels in both BNL-CL2 (according to transcriptomics) and foetal liver tissues (according to qPCR) compared to other TRPVs (Figure S8A, B), and it could be detected in the immunoblotting in foetal hepatocytes and livers (Figure S8C). In addition, low glucose/FBP enhanced the aldolase-TRPV4 interaction (Figure 3A), as in HEK293T cells^10^. Surprisingly, the TRPV activity, evidenced by the fluorescence GCaMP6s-TRPV4 indicator^10^, remained unaffected in low glucose levels in foetal hepatocytes (Figure 3B, D), whereas the TRPV activity decreased in adult liver or 293T cells under the same low glucose condition (Figure 3C, E, F). When TRPV was pharmacologically inhibited by the antagonist BCTC, as evidenced by the decreased GCaMP6s-TRPV4 fluorescence (Figure 3B, D), mTORC1 activity was decreased significantly in foetal hepatocytes (Figure 3G), and mTORC1 was dissociated from the lysosomes, regardless of glucose availability (Figure 3H, I). We also found that BCTC could inhibit the activity of v-ATPase, likely as a consequence of the inhibition of TRPV (Figure S8D, E), and effectively triggered the lysosomal translocation of AXIN in foetal hepatocytes (Figure S8F, G). In addition, the TRPV agonist GSK101 did not change the already high mTORC1 activity in foetal hepatocytes; in contrast, GSK101 sufficiently overrode the low-glucose-induced inhibition of TRPV to restore mTORC1 activity in adult hepatocytes (Figure 3J, K). These results suggest that TRPVs are not inhibited by low glucose in foetal liver cells, such that the low glucose signal cannot be transmitted to inhibit v-ATPase or to trigger the lysosomal translocation of AXIN, events required for mTORC1 dissociation from the lysosome^11,13^.

**Figure 3.**
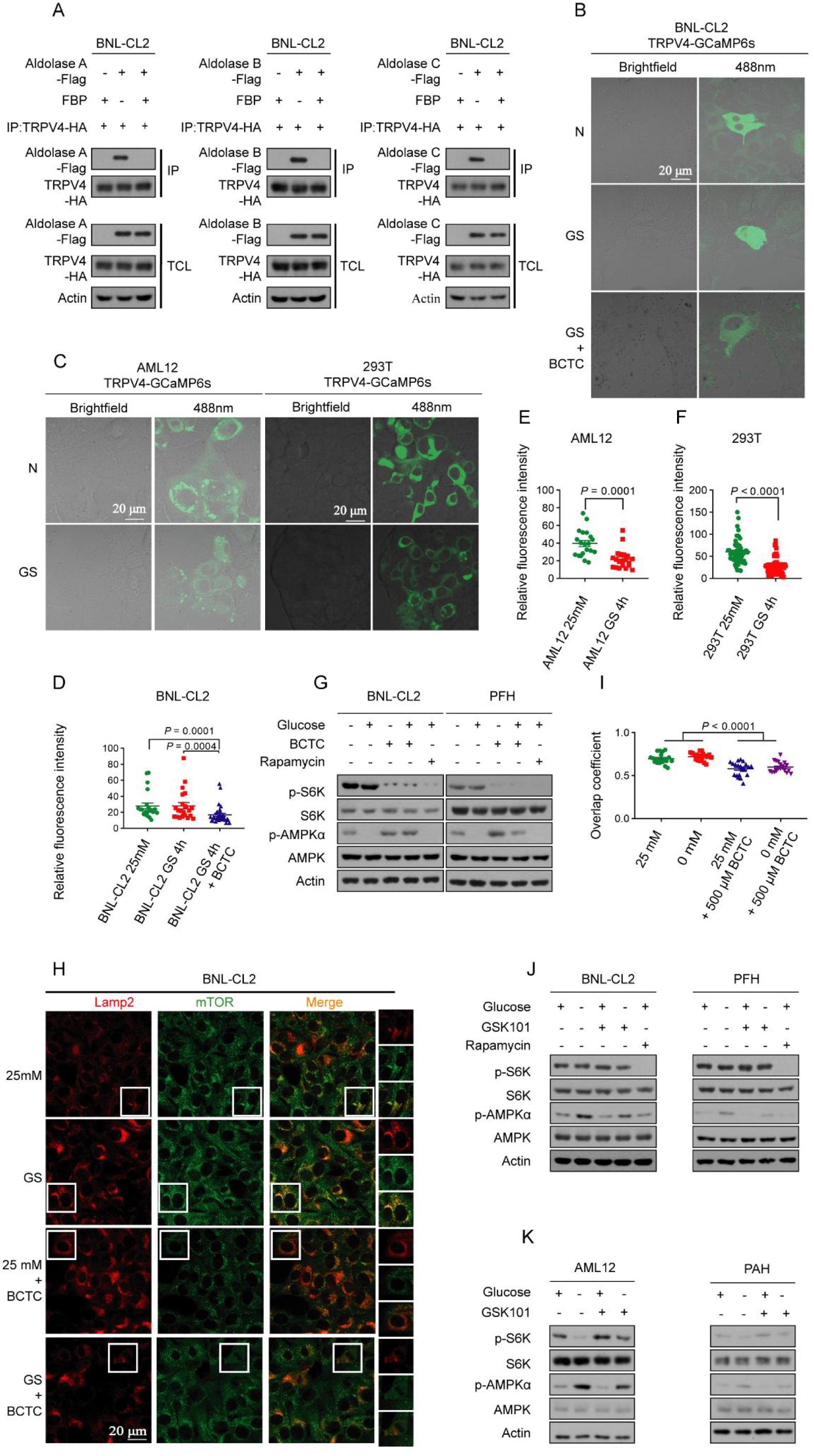
Low glucose fails to trigger the lysosomal AMPK pathway in the foetal liver. A. Aldolase is capable of binding to TRPV channels in foetal hepatocytes. The BNL-CL2 (foetal hepatocytes) cells stably expressing HA-tagged TRPV4 were glucose-starved for 4 h. Cells were then lysed, and the lysates were incubated with 200 μM FBP for 12 h. The interaction between aldolase and TRPV4 in the lysates was then determined by immunoprecipitation of HA-tag, followed by immunoblotting. B-F. The TRPV channels in foetal hepatocytes remain active even in low glucose conditions. The BNL-CL2 (B, D; foetal hepatocytes), AML12 (left panel of C, and E; adult hepatocytes), and HEK293T cells (right panel of C, and F), all stably expressing TRPV4-GCaMP6s indicator, were glucose-starved for 4 h The intensities of the indicator were then determined. As a control, the TRPV inhibitor BCTC at 500 μM was applied to the cells for 2 h. Statistical analysis data are shown as mean ± SEM; BNL-CL2 25mM n=22 (D), BNL-CL2 GS 4 h n=22 (D), BNL-CL2 GS 4 h + BCTC n=32 (D); AML12 25mM n=19 (E), AML12 GS 4 h n=19 (E); 293T 25mM n=53 (F), 293TAML12 GS 4 h n=62 (F). P values by unpaired two-tailed Mann-Whitney test (D) and unpaired two-tailed Student’s *t*-test (E, F). G-I. Forced inhibition of TRPV suppresses mTORC1 in foetal hepatocytes. The BNL-CL2 (G, H; foetal hepatocytes) cells or primary foetal hepatocytes (G) were treated with 500 μM BCTC for 2 h or 100 nM rapamycin for 1 h, or glucose-starved for 4 h, followed by determination of p-S6K by immunoblotting (G), or the lysosomal localization of mTOR by immunofluorescent staining (H; and the Manders’ overlap coefficients are shown in I as mean ± SEM, 25mM n=14, 0mM n=15, 25mM + BCTC n=15, 0mM + BCTC n=13; P values by unpaired two-tailed Student’s *t*-test, except for 0mM vs 25mM + BCTC where unpaired two-tailed Mann-Whitney test is used). J, K. Activation of TRPV does not further activate mTORC1 in foetal hepatocytes. The BNL-CL2 (J, left panel) cells or primary foetal hepatocytes (J, right panel), along with adult hepatocytes as a control (K), were treated with glucose-starved or 1 mM GSK101, the TRPV agonist, for 4 h, followed by determination of p-S6K by immunoblotting. Experiments in this figure were performed three times, except A one time.

We also investigated why AMPK can still be activated in low glucose in foetal hepatocytes while the lysosomal pathway is apparently not operating. To that end, we determined the levels of ATP, ADP, and AMP. We found that glucose starvation significantly increased the ratios of AMP:ATP and ADP:ATP in foetal hepatocytes (Figure S8H-J). This indicates that the foetal AMPK is allosterically activated through the canonical, AMP-dependent mechanism^49^.

### Acetylation of TRPV during intrauterine development confers insensitivity to low glucose

We next investigated how TRPV is resistant to inhibition by FBP-unoccupied aldolase under low glucose conditions. We found that the protein level of TRPV in foetal livers was lower than that in adult livers (Figure S9A-B). However, as BCTC could inhibit foetal liver TRPV (Figure 3B, D, G-I; Figure S8D-G), we concluded that the relatively low protein levels of TRPV in foetal livers, compared to those in adult livers, cannot explain the resistance of foetal TRPV to inhibition by low glucose. We therefore explored whether alteration of the post-translational modifications of TRPV may contribute to the constitutive activation of this cation channel. Some 41 acetylated residues, all lysine, were hit by protein mass spectrometry. We then individually mutated the 41 lysine residues to arginine (K to R mutation) to mimic the deacetylated state, and found that the TRPV4-K608R mutant, when expressed in foetal hepatocytes, rendered mTORC1 sensitive to low glucose levels (Figure 4A). Importantly, the levels of acetylated TRPV4-K608 were significantly higher in foetal liver/hepatocytes than in adult liver/hepatocytes, as confirmed by quantitative mass spectrometry analysis (Figure S9C-M). The K608 residue is also highly conserved among various species and among different members of the TRPV family (Figure S10A). In addition, we found that the v-ATPase inhibition (Figure S10B-D), AXIN translocation to the lysosome (Figure S10E-H), and the mTORC1 dissociation from the lysosome (Figure 4B-E), could all be observed in foetal hepatocytes expressing TRPV4-K608R, as seen in adult cells, although the mutation of K608 did not affect the basal activity of TRPV4 (Figure S10I, J) or change its affinity with the FBP-unoccupied aldolase (Figure S10K). We therefore concluded that the acetylation of K608 renders TRPV4 insensitive to the low glucose-/FBP-unoccupied aldolase-induced inhibition.

**Figure 4.**
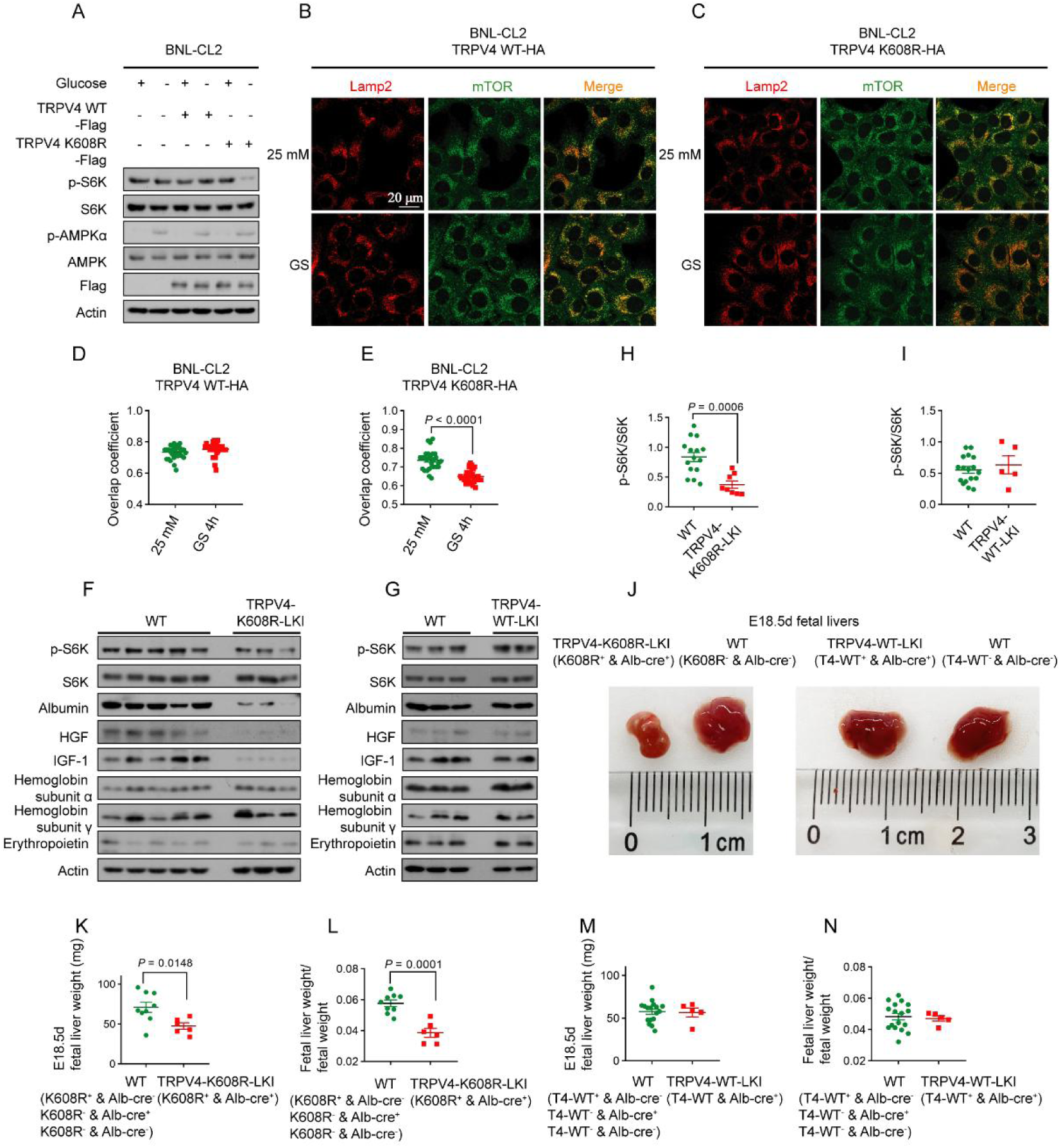
Acetylation of TRPV during intrauterine development confers insensitivity to low glucose. A-E. The acetylation-defective, TRPV4-K608R mutant dominantly inhibits mTORC1 in foetal hepatocytes in low glucose. The BNL-CL2 cells stably expressing TRPV4-K608R mutant or wildtype TRPV4 as a control, were glucose-starved for 4 h, followed by determination of the levels of p-S6K (A) and the lysosomal localization of mTOR (representative images shown in B and C, while Manders’ overlap coefficient, as mean ± SEM in D and E; n=30 cells for each condition (D); 25mM n=28 cells (E), GS 4 h n=34 cells (E) and P values by unpaired two-tailed Student’s *t*-test (D) and by unpaired two-tailed Mann-Whitney test (E). F-I. TRPV4-K608R dominantly inhibits mTORC1 in foetal livers. Liver tissues collected from E18.5d mice with liver-specific TRPV4-K608R (F, H) or wildtype TRPV4 (G, I) knocked-in were lysed, followed by determining p-S6K levels by immunoblotting. Statistical analysis data are shown as mean ± SEM, WT n=15 (H), TRPV4-K608R-LKI n=8 (H), WT n=17 (I), TRPV4-WT-LKI n=5 (I) and P values by unpaired two-tailed Student’s *t*-test. J-N. TRPV4-K608R causes liver abnormalities during the foetal development. The appearance of the livers (J), along with liver weight (K, M, mean ± SEM, WT n=9 (K), TRPV4-K608R-LKI n=6 (K), WT n=17 (M), TRPV4-WT-LKI n=5 (M) and P values by unpaired two-tailed Student’s *t*-test), and the ratios of liver weight/body weight (L, N, mean ± SEM, WT n=9 (L), TRPV4-K608R-LKI n=6 (L), WT n=17 (N), TRPV4-WT-LKI n=5 (N) and P values by unpaired two-tailed Student’s *t*-test) of liver-specific (induced by the Alb-CRE promoter) TRPV4-K608R (J-L) or wildtype TRPV4 (M, N) knocked-in mice at E18.5d were determined. Experiments in this figure were performed three times, except F-I one time.

To investigate the physiological significance of TRPV acetylation during foetal development in utero, we generated transgenic mice with liver-specific expression of K608R-TRPV4 (induced by *Alb*-*Cre*; validated in Figure S11A-C). We then examined the phenotypes of the foetus in these mice. As shown in Figure S11D, approximately 18% (2/11) of embryos with liver-specific expression of K608R-TRPV4 (TRPV4-K608R-LKI) died during intrauterine development. Among the survived embryos, we observed a significant decrease of mTORC1 activity in the liver (Figure 4F-I). In agreement with a lower anabolic activity resulting from the inhibition of mTORC1, the livers of TRPV4-K608R-LKI embryos had lowered liver weights, with lower levels of albumin, hepatocyte growth factor (HGF), IGF-1, compared to their wild-type littermates (Figure 4J-N, Figure S12A-F). However, there were no differences in haemoglobin and erythropoietin levels between the two groups (Figure S12G-L). In addition, we quantified the number of surviving offspring from these mice, and found that none of offspring expressing K608R-TRPV4 survived to the weaning stage (Details showed in Table. S1). The TRPV4-K608R-LKI embryos were found dead, usually within several hours postpartum. Consistently, we found that rapamycin administration in wild-type embryos, which inhibits mTORC1 in the livers, recapitulates the postnatal death observed in the TRPV4-K608R-LKI embryos (Figure S12M-R). Note that although we attempted to block the foetal mTORC1 signalling by knocking out *Raptor* in the liver as well, it turned out to be unsuccessful. The protein levels of Raptor remained mostly unaffected in the foetal liver (Figure S12S-U), which is in stark contrast to those in adult livers (significantly decreased in the liver of *Raptor*^F/F^ mice crossed with the *Alb-Cre* mice)^50^. This observation may also explain why liver-specific knockout of *Raptor* can survive normally and remain fertile, although smaller sizes of the adult livers were observed^50^. Together, we found that the K608R-TRPV4 acetylation, through maintaining the mTORC1 activity in low glucose in the foetal liver, plays a critical role in embryo development.

## DISCUSSION

During pregnancy, most of the glucose provided by the uterine artery is consumed by the uterus and placenta, and only about a fraction of one-third can reach the foetus^51,52^. Therefore, the glucose concentrations in the foetus are lower compared to that in the mother, displaying a transplacental gradient of glucose concentration during embryonic development^53^. In addition, it was reported that in pregnant women with insulinoma, who constantly experience higher rates of insulin secretion and subsequently suffer from hyperinsulinemia, there are no significant defects in the survival and development of the foetus^54,55^. This requires a mechanism that sustains mTORC1 activity, as it is essential for maintaining the anabolic activities in the foetal livers amid maternal glucose fluctuations.

During embryogenesis, the liver is formed from hepatoblasts, which are derived from definitive endoderm at around 3-4 weeks post-conception in humans, or at E8.5-E9.0 in mice^56^. In mice, the differentiation of hepatoblasts into hepatocytes and cholangiocytes starts at E13.5, while in humans, mature hepatobiliary cells can be observed at around 7 weeks post-conception^57^. Our work also identified mTORC1 as a critical mediator for the development of the liver, in addition to its general roles in cell proliferation in both extraembryonic and embryonic compartments^44^. As we observed lowered foetal glucose levels from E10.5 to P1 in pregnant mice without dietary restrictions (Figure 1L), and the TRPV4-K608R-LKI foetal liver, in which mTORC1 is inhibited by low glucose, displays developmental abnormalities, it is evident that constitutively active mTORC1 plays a crucial role during the developmental stage of the foetal liver, which is continually exposed to low glucose levels.

Our current study has revealed the mechanism that prevents the glucose-sensing pathway from operating in the foetal for the sustainability of anabolic activities. We have found that the acetylation of TRPV4 on K608 residue renders TRPV4 unaffected by the FBP-unoccupied aldolase, which makes TRPV as a checkpoint for the glucose sensing-triggered AMPK activation and mTORC1 inhibition. Models of glucose-sensing pathway in adult and foetal livers are shown in Figure S13. Notably, while the glucose sensing-mediated AMPK activation cannot take place in the foetal liver, AMPK is effectively activated by the canonical AMP-dependent mechanism. It must be pointed out that it remains enigmatic why the canonically activated AMPK fails to inactivate Raptor or enhance TSC2, which collectively inactivates mTORC1 in the foetal liver.

The glucose sensing pathway begins to operate in the liver right after birth, highlighting the importance of regulating mTORC1 activity for maintaining the balance between anabolism and catabolism during postnatal development. Previous studies have shown that sustained RagA activity through introducing RagA-Q66L mutant (RagA^GTP^) allows mTORC1 to maintain constitutive activity. While RagA^GTP/GTP^ mice can develop normally in the uterus, they are unable to survive for more than 24 h after birth^8^. Therefore, the transition from the constitutive activity in the foetus to dynamically regulated mTORC1 after birth is crucial for the normal development of the newborns.

## RESOURCE AVAILABILITY

### Lead contact

Supplementary Information is available for this paper. Correspondence and requests for materials should be addressed to He-Feng Huang.

### Data availability

All software and data used are freely available either online through various servers (see Key Resources Table). The MS proteomics data has been deposited to the ProteomeXchange Consortium (http://proteomecentral.proteomexchange.org) through the iProX partner repository with the dataset identifier IPX0009924000. The transcriptome raw data of mice foetal liver cell line BNL-CL2 can be accessed from NCBI SRA BioProject ID: PRJNA1171108 (SRA records will be accessible with the following link after the indicated release date: https://www.ncbi.nlm.nih.gov/sra/PRJNA1171108). The analysis was performed using standard protocols with previously described computational tools. No custom code was used in this study. Full immunoblots are provided as Supplementary Figure1 in Supplementary Information. Source data are provided with this paper.

### Code availability

The analysis was performed using standard protocols with previously described computational tools. No custom code was used in this study.

## ACKNOWLEDGEMENTS

This work is supported by the National Natural Science Foundation of China (82088102, 92357306), the National Key R&D Program of China (2022YFC2703500), CAMS Innovation Fund for Medical Sciences (2019-I2M-5-064), Collaborative Innovation Program of Shanghai Municipal Health Commission (2020CXJQ01), Key Discipline Construction Project (2023-2025) of Three-Year Initiative Plan for Strengthening Public Health System Construction in Shanghai (GWVI-11.1-35), Shanghai Clinical Research Center for Gynecological Diseases (22MC1940200), Shanghai Urogenital System Diseases Research Center (2022ZZ01012) and Shanghai Frontiers Science Research Center of Reproduction and Development. We thank Y. Wu, C. Xie, and Z. Xu (Xiamen University) for help with mass spectrometry analysis; J. Wu (Xiamen University) for help with transgenic mice construction; C. Zhang and M. Zhu (Xiamen University) for help with metabolomics analysis, and L. Wang (Xiamen Cardiovascular Hospital of Xiamen University) for the critical discussions.

## AUTHOR CONTRIBUTIONS

C.-J.Z. and C.-J.Y. are responsible for the experimental work. J.-X.P., L.-Y.Y. and Y.-H.T. provided clinical samples. Y.C., X.-Y.Q. and Z.-L.L. assisted in completing the experiment. C.-S.Z., S.-C.L., H.-F.H. and G.-L.D. are responsible for experimental guidance, and C.-J.Z., C.-J.Y., C.-S.Z., S.-C.L., H.-F.H. and G.-L.D. are responsible for article writing.

## COMPETING INTERESTS

The authors declare no competing interests.

## Supplementary Information

**Figure S1.**
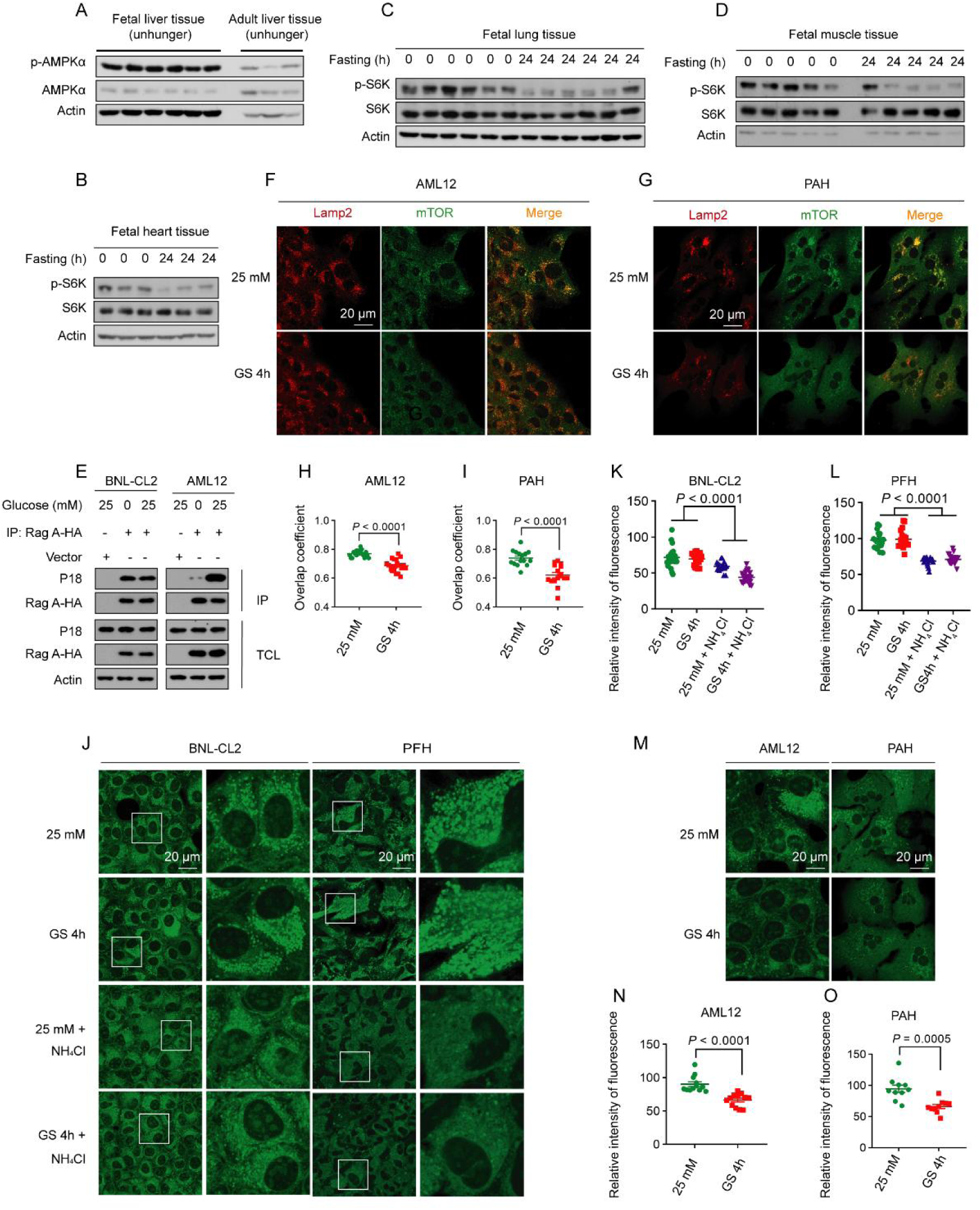
mTORC1 remains active in foetal hepatocytes in low glucose, but not in adult hepatocytes, related to Figure 1 and Figure 2. A. The basal AMPK activation in foetal liver is much higher than that in adult liver, n=6 for foetal liver tissue, n=3 for adult liver tissue. B-D. mTORC1 is significantly inhibited after starvation in the foetal heart, muscle, and liver. Tissues were collected from pregnant mice embryos (E18.5d) after being fasted for 0 h or 24 h, followed by determination of p-S6K by immunoblotting. E. mTORC1 remains associated with RAG in BNL-CL2 in low glucose. See also AML12 cells, as a control, in which mTORC1 is not associated with RAG. F-I. mTORC1 dissociates from the lysosomes in low glucose conditions in adult hepatocytes. The AML12 adult hepatocyte cell line (F, H) and primary adult hepatocytes (G, I) were glucose-starved for 4 h. Endogenous mTOR (labeled in green with rabbit anti-mTOR antibody, CST 2983T) and the lysosome marker LAMP2 (labeled in red with rat anti-LAMP2 antibody, Abcam AB13524) were then stained. Representative images are shown in F and G, and the Mander’s overlap coefficients in H and I (presented as mean ± SEM, n = 19 for each group of H or 15 for each group of I, with P values calculated using unpaired two-tailed Mann-Whitney test (H) and unpaired two-tailed Student’s *t*-test (I)). J-O. Glucose starvation does not affect the lysosomal acidity in foetal hepatocytes. The BNL-CL2 (left panel of J, and K), primary foetal hepatocytes (right panel of J, and L), AML12 (left panel of M, and N), and primary adult hepatocytes (right panel of M, and O) were either glucose-starved for 4 h or treated with 25 mM NH_4_Cl, the lysosome inhibitor, as a control for 2 h. In contrast, low glucose effectively inhibited v-ATPase in adult hepatocytes. The cells were incubated with 1 μM Lysosensor^TM^ Green DND-189 (Thermo L7535) for 30 min to detect the PH of lysosomes. Representative images are shown in J and M, and the Mander’s overlap coefficients in K, L, and N, O (presented as mean ± SEM, n = 25 for 25 mM group of K, n = 22 for 0 mM group of K, n = 21 for 25 mM + NH_4_Cl group of K, n = 24 for 0 mM + NH_4_Cl group of K; n = 10 for each group of L, n = 12 for each group of N, N = 10 for each group of O, with P values calculated using unpaired two-tailed Student’s *t*-test, except for 25 mM vs 25 mM + NH_4_Cl (K), 25 mM vs 0 mM + NH_4_Cl (K), 25 mM vs 25 mM + NH_4_Cl (L), 0 mM vs 25 mM + NH_4_Cl (L) where unpaired two-tailed Mann-Whitney test is used). Experiments in this figure were performed three times.

**Figure S2.**
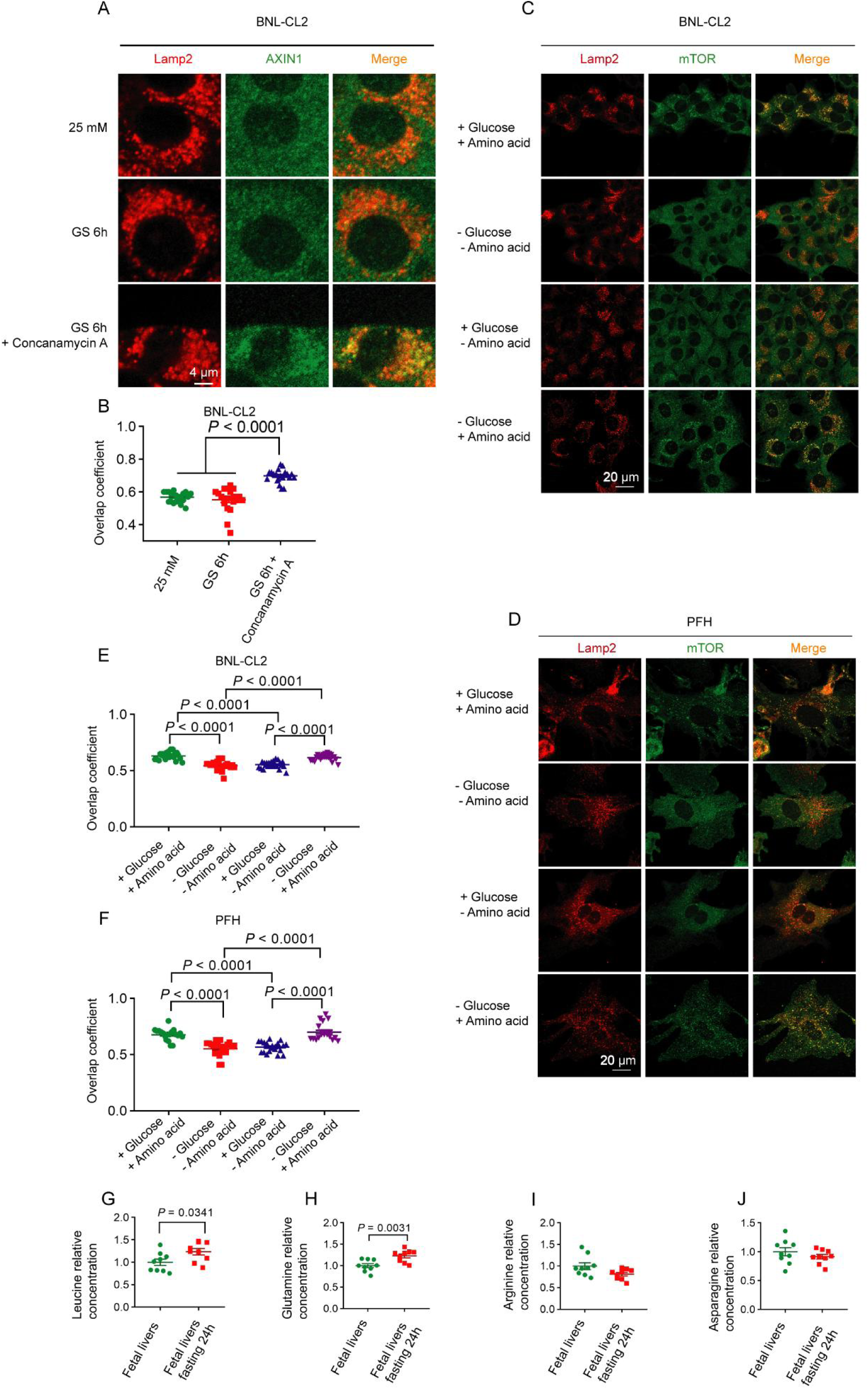
Constitutively active v-ATPase-Ragulator-RAG axis maintains mTORC1 activity in foetal hepatocytes in low glucose, related to Figure 2. A, B. Forced inhibition of v-ATPase in foetal hepatocytes by concanamycin A (conA) effectively triggers the lysosomal translocation of AXIN. BNL-CL2 were either glucose-starved or treated with 1μM Concanamycin A, the v-ATPase inhibitor, as a control for 6 h. Endogenous AXIN1 (labeled in green with goat anti-AXIN1 antibody, Santa Cruz Biotechnology SC-8567) and the lysosome marker LAMP2 (labeled in red with rat anti-LAMP2 antibody, Abcam AB13524) were then stained. Representative images are shown in A, and the Mander’s overlap coefficients in B (presented as mean ± SEM, n = 15 for each group, with P values calculated using unpaired two-tailed Student’s *t*-test, except for GS 6 h vs GS 6 h + Concanamycin A where unpaired two-tailed Mann-Whitney test is used). C-F. mTORC1 dissociates from the lysosomes in amino acid deficiency conditions in foetal hepatocytes. The BNL-CL2 (C, E) and primary foetal hepatocytes (D, F) were either glucose-starved or amino acid-starved for 2 h. Endogenous mTOR (labeled in green with rabbit anti-mTOR antibody, CST 2983T) and the lysosome marker LAMP2 (labeled in red with rat anti-LAMP2 antibody, Abcam AB13524) were then stained. Representative images are shown in C and D, and the Mander’s overlap coefficients are shown in E and F (presented as mean ± SEM, n = 13 for + Glucose + Amino acid group of E, n=15 for - Glucose - Amino acid group of E, n=13 for + Glucose - Amino acid group of E, n=13 for - Glucose + Amino acid group of E; n=10 for + Glucose + Amino acid group of F; n=11 for - Glucose - Amino acid group of F; n=10 for + Glucose - Amino acid group of F; n=10 for - Glucose + Amino acid group of F, with P values calculated using unpaired two-tailed Student’s *t*-test, except + Glucose + Amino acid vs + Glucose - Amino acid (E) where unpaired two-tailed Mann-Whitney test is used). G-J. Maternal fasting does not decrease the levels of amino acids, such as leucine, glutamine, arginine, and asparagine, all of which can regulate mTORC1 activity, in the foetal liver. Liver tissues were collected from E18.5d foetal mice after the pregnant females were fasted for 0 h or 24 h, followed by determination of amino acid levels through HPLC-MS analysis. The results show no significant change in the relative concentration of leucine (G), glutamine (H), arginine (I), and asparagine (J) in foetal liver tissues between the fasting and control groups (presented as mean ± SEM, n=9 for each group of G-J, with P values calculated using unpaired two-tailed Student’s *t*-test). Experiments in this figure were performed three times.

**Figure S3.**
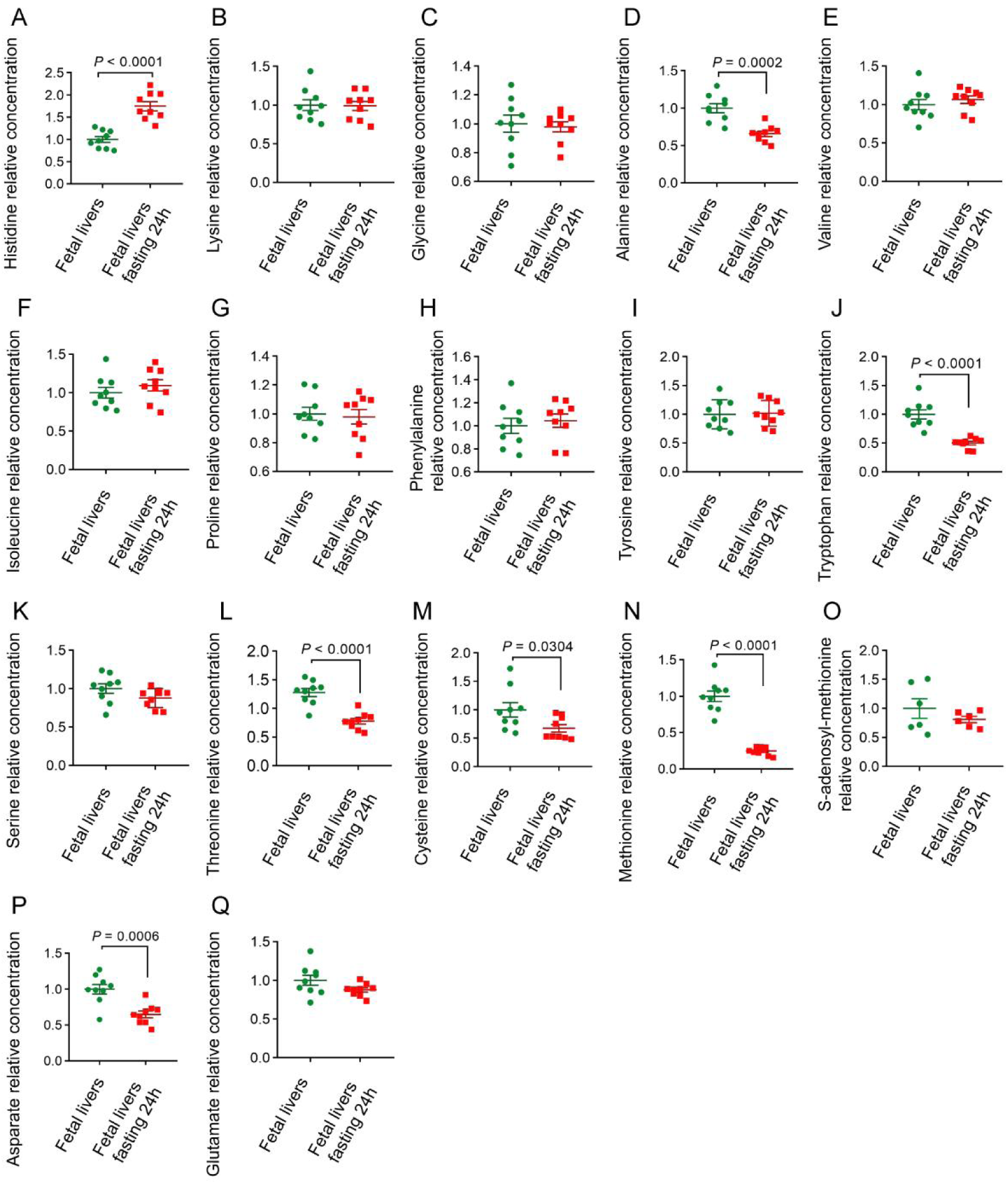
The overall level of amino acids in the liver of E18.5d foetal mice does not significantly change after the pregnant females being fasted, related to Figure 2. A-Q. Liver tissues were collected from E18.5d foetal mice after the pregnant females were fasted for 0 h or 24 h, followed by the detection of the relative concentration of histidine (A), lysine (B), glycine (C), alanine (D), valine (E), isoleucine (F), proline (G), phenylalanine (H), tyrosine (I), tryptophan (J), serine (K), threonine (L), cystine (M), methionine (N), s-adenosyl-methionine (O), asparate (P), glutamate (Q) in foetal liver tissues between the fasting and control groups using HPLC-MS (presented as mean ± SEM, n = 9 for each group of A-N, P, Q, except n = 6 for each group of O, with P values calculated using unpaired two-tailed Student’s *t*-test (A-I, K-M, P) and unpaired two-tailed Mann-Whitney test (J, N, O, Q).

**Figure S4.**
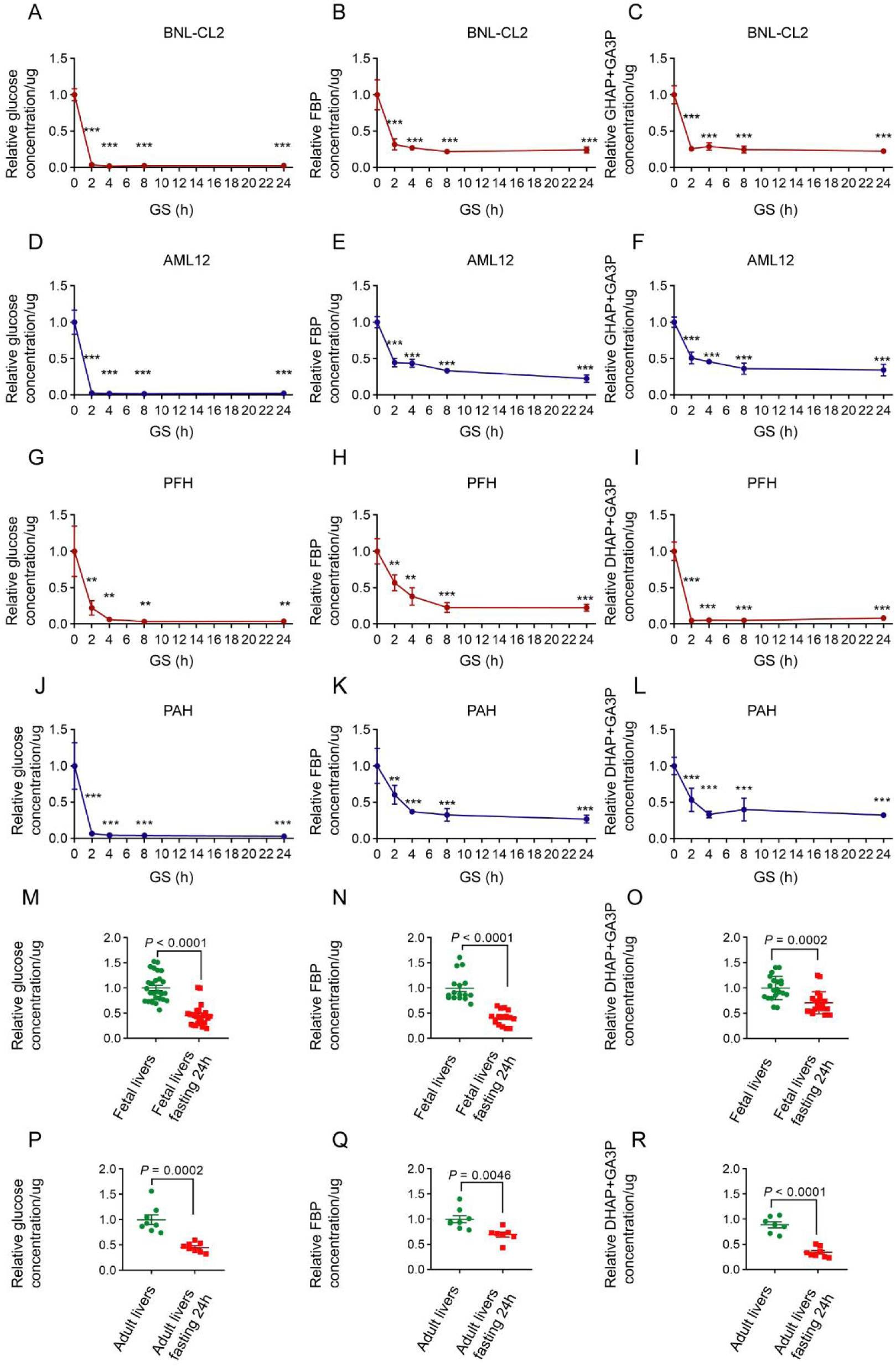
FBP and DHAP decrease in the foetal liver and adult livers in low glucose, related to Figure 3. A-L. Levels of FBP and DHAP are decreased in both foetal hepatocytes and adult hepatocytes in low glucose. BNL-CL2 (A-C), AML12 (D-F), PFH (G-I), and PAH (J-L) were glucose-starved for 0 h, 2 h, 4 h, 8 h, or 24 h, followed by determination of FBP and DHAP by HPLC-MS (presented as mean ± SEM, n = 5 for each condition of A-F, n = 4 for each condition of G-I, n = 6 for each condition of J-L, except n = 3 for PFH 0 h group of G-I, with P values calculated using unpaired two-tailed Student’s *t*-test; *: *P* <0.05; **: *P* <0.01; ***: *P* <0.001, all compared with 0 h group). M-R. Levels of FBP and DHAP are decreased in the liver tissue of foetal mice after maternal fasting (M-O), or in adult mouse livers (P-R). Liver tissues were collected from pregnant mice embryos (E18.5d; foetal liver tissue) or adult mice (8-week-old; adult liver tissue), after being fasted for 0 h or 24 h. FBP and DHAP were determined by HPLC-MS in these cells (presented as mean ± SEM, n = 27 for foetal livers group of M, n = 24 for foetal livers fasting 24 h group of M; n = 16 for each group of N, n = 20 for each group of o, n = 8 for each group of P; n = 8 for adult livers group of Q, n = 7 for adult livers fasting 24 h group of Q; n = 7 for adult livers group of R, n = 8 for adult livers fasting 24 h group of R; with P values calculated using unpaired two-tailed Student’s *t*-test (M, O, Q, R) and unpaired two-tailed Mann-Whitney test (N, P).

**Figure S5.**
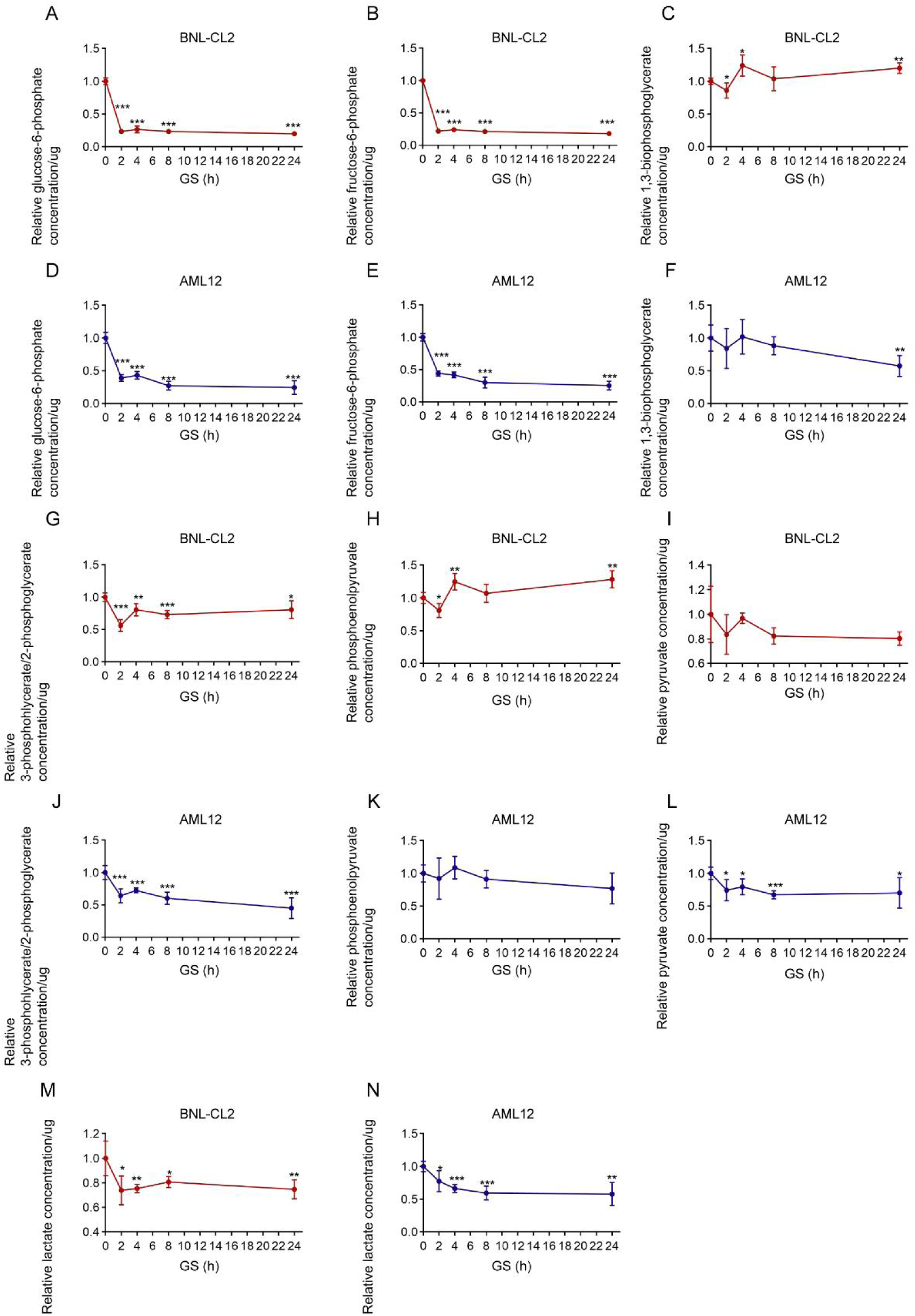
Changes of glycolytic intermediates, excepting FBP and DHAP, in hepatic cell lines after glucose starvation, related to Figure 3. A-N. The levels of glucose-6-phosphate, fructose-6-phosphate, 3-phosphoglycerate/2-phosphoglycerate, and lactate are decreased in both BNL-CL2 and AML12 cells in low glucose. BNL-CL2 (A-C, G-I, M) and AML12 (D-F, J-L, N) were glucose-starved for 0 h, 2 h, 4 h, 8 h, or 24 h. HPLC was used to determine glycolytic intermediates in these cells (presented as mean ± SEM, n = 5 for each condition of A-N, with P values calculated using unpaired two-tailed Student’s *t*-test; *: *P* <0.05; **: *P* <0.01; ***: *P* <0.001, all compared with 0 h group).

**Figure S6.**
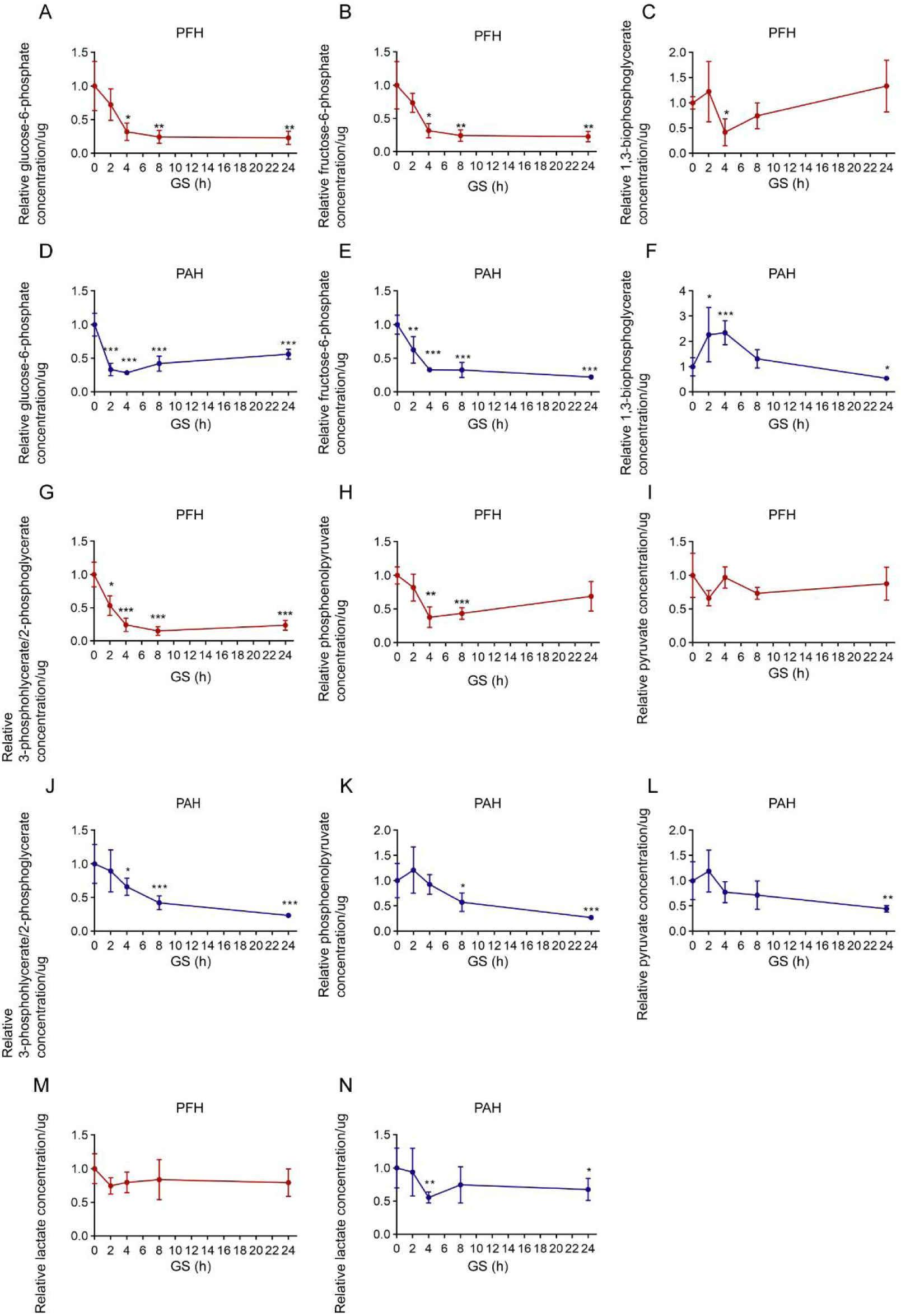
Changes of glycolytic intermediates, excepting FBP and DHAP, in primary hepatocytes after glucose starvation, related to Figure 3. A-N. The levels of glucose-6-phosphate, fructose-6-phosphate, 3-phosphoglycerate/2-phosphoglycerate, and lactate are decreased in both PFH and PAH in low glucose. PFH (A-C, G-I, M) and PAH (D-F, J-L, N) were glucose-starved for 0 h, 2 h, 4 h, 8 h, or 24 H. HPLC was used to determine glycolytic intermediates in these cells (presented as mean ± SEM, n = 4 for each condition of A-C, G-I, M, n = 6 for each condition of D-F, J-L, N, except n = 3 for PFH 0 h group of A-C, G-I, M with P values calculated using unpaired two-tailed Student’s *t*-test; *: *P* <0.05; **: *P* <0.01; ***: *P* <0.001, all compared with 0 h group).

**Figure S7.**
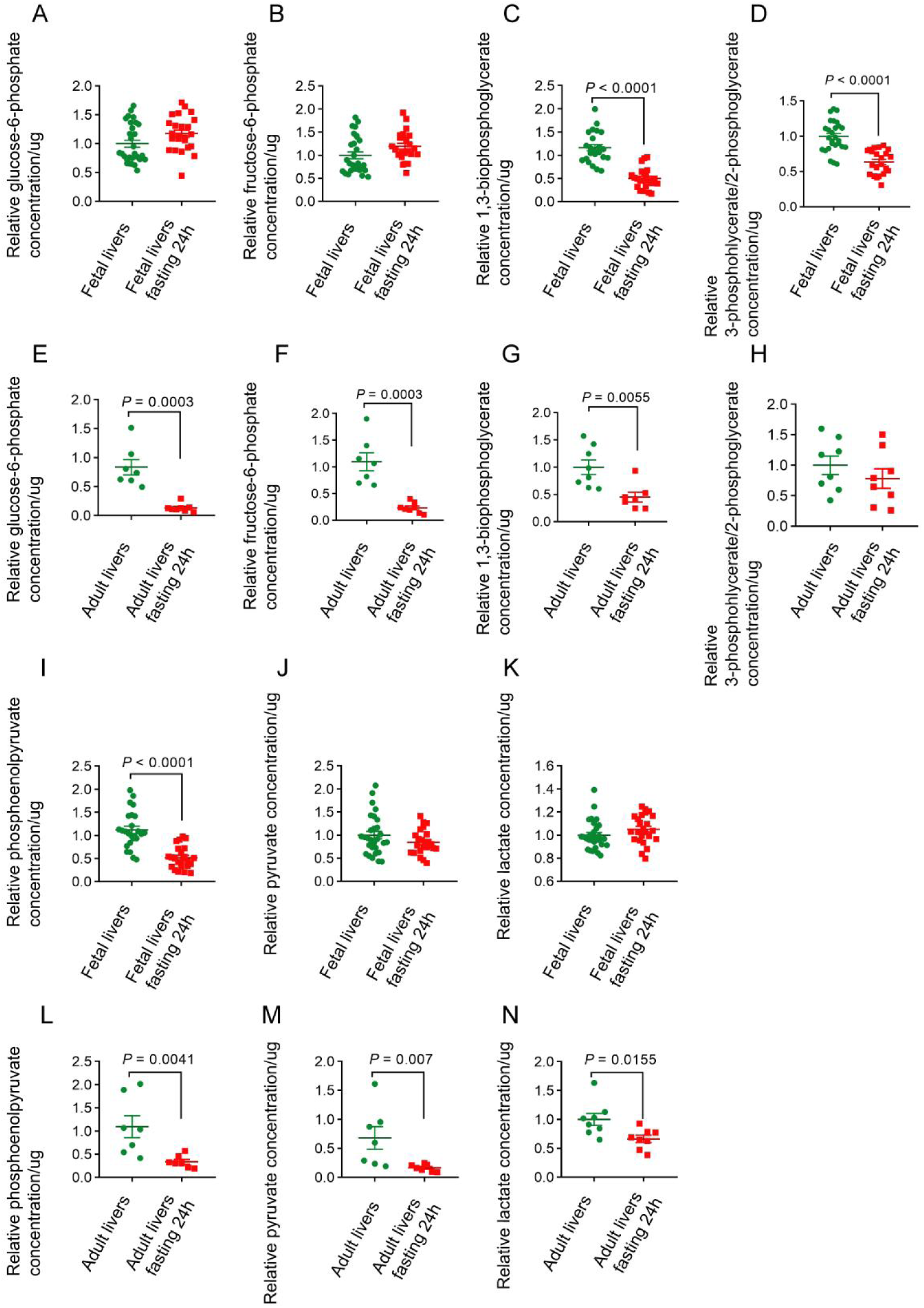
Changes of glycolytic metabolites, excepting FBP and DHAP, in the livers of E18.5d foetal mice after the pregnant females being fasted, related to Figure 3. A-N. Liver tissues collected from pregnant mouse embryos (E18.5d; foetal liver tissue) or adult mice (8-week-old; adult liver tissue) after fasting for 0 h or 24 h were subjected to HPLC to determine glycolytic intermediates (presented as mean ± SEM, n = 29 for foetal livers group of A, B, J, K, n = 24 for foetal livers fasting 24 h group of A, B, J, K; n = 23 for foetal livers group of C, n = 22 for foetal livers fasting 24 h group of C; n = 24 for foetal livers group of D, n = 19 for foetal livers fasting 24 h group of D; n = 25 for foetal livers group of I, n = 23 for foetal livers fasting 24 h group of I; n = 8 for each group of H, N; n=7 for each group of L, M; with P values calculated using unpaired two-tailed Student’s *t*-test (A-D, G, H, K, N) and unpaired two-tailed Mann-Whitney test (E, F, I, J, L, M).

**Figure S8.**
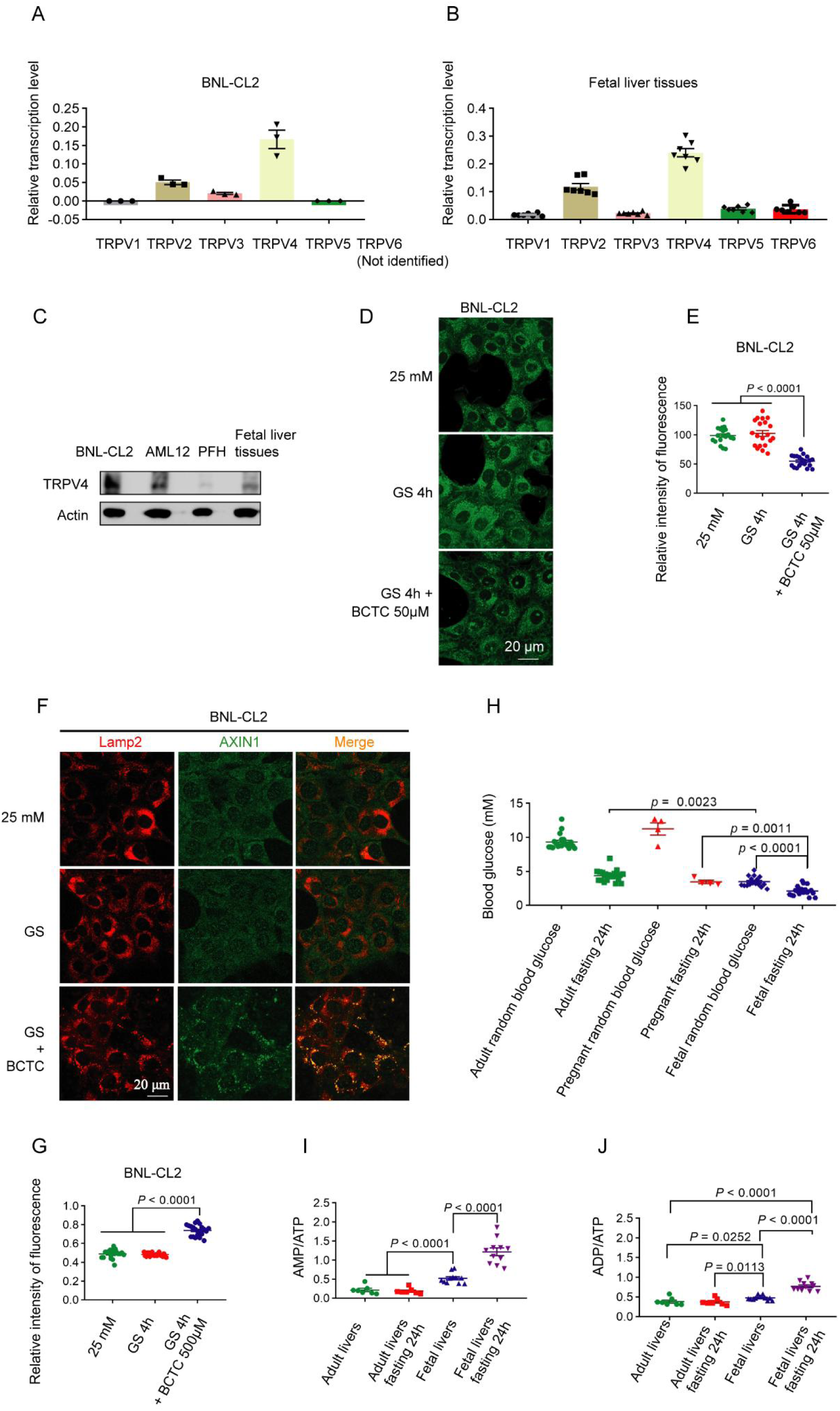
Foetal AMPK is allosterically activated through the canonical, AMP-dependent mechanism, related to Figure 3. A. TRPV4 has the highest mRNA levels in BNL-CL2 cells (by transcriptomics). Data presented as mean ± SEM, n = 3 for each group. B. TRPV4 has the highest mRNA levels in E18.5d foetal liver (by qPCR). Data presented as mean ± SEM, n = 7 for each group. C. TRPV4 can be detected in BNL-CL2, AML12, PFH, and foetal liver tissues. D, e. Forced inhibition of TRPVs in foetal hepatocytes can inhibit the activity of v-ATPase. The BNL-CL2 (D) was either glucose-starved or treated with 50 μM BCTC, the TRPV inhibitor, as a control for 4 h. The cells were incubated with 1μM Lysosensor^TM^ Green DND-189 (Thermo L7535) for 30 min to detect the PH of lysosomes. Representative images are shown in D, and the Mander’s overlap coefficients in E (presented as mean ± SEM, n = 18 for 25 mM group of E, n=19 for GS 4 h group of E, n = 16 for GS 4 h + 50 μM BCTC group of E, with P values calculated using unpaired two-tailed Mann-Whitney test, except 25 mM vs GS 4 h + 50 μM BCTC where unpaired two-tailed Student’s *t*-test is used). F, G. Forced inhibition of TRPVs in foetal hepatocytes by BCTC effectively triggers the lysosomal translocation of AXIN in foetal hepatocytes. BNL-CL2 were either glucose-starved for 4 h or treated with 500 μM BCTC, the TRPV inhibitor, for 2 h. Endogenous AXIN1 (labeled in green with goat anti-AXIN1 antibody, Santa Cruz Biotechnology SC-8567) and the lysosome marker LAMP2 (labeled in red with rat anti-LAMP2 antibody, Abcam AB13524) were then stained. Representative images are shown in F, and the Mander’s overlap coefficients in G (presented as mean ± SEM, n = 18 for 25 mM group, n = 18 for GS 4 h group, n = 21 for GS 4 h + BCTC 500 μM group, with P values calculated using unpaired two-tailed Mann-Whitney test). H. Fasted pregnant mice (foetuses of E18.5d) and 8-w female adult mice resulted in a significant decrease of plasma glucose after 24-h fasting (from 9.3 to 4.4 mM in the adults, from 11.3 to 3.5 mM in the pregnant mice, and from 3.5 to 2.1 mM in the foetus). Blood samples were collected from adult mice (8-week-old; adult blood glucose), pregnant mice (E18.5d; pregnant blood glucose), or embryos (E18.5d; foetal blood glucose), after being fasted for 0 h or 24 h. Blood glucose levels were detected by a portable human glucometer (Accu-Chek; Roche). Data are presented as mean ± SEM, n = 20 for each adult group of H, n = 4 for each pregnant group of H, n = 16 for foetal random blood glucose group of H, n = 21 for foetal fasting 24 h group of H, with P values calculated using unpaired two-tailed Student’s *t*-test. I, J. Glucose starvation significantly increases the ratios of AMP:ATP and ADP:ATP in foetal hepatocytes. Liver tissues were collected from pregnant mice embryos (E18.5d; foetal liver tissue) and adult mice (8-week-old; adult liver tissue), after being fasted for 0 h or 24 h. HPLC was used to detect energy levels in tissues (presented as mean ± SEM, n = 7 for each adult group of I, J, n = 11 for each foetal group of I, J with P values calculated using unpaired two-tailed Student’s *t*-test, except foetal livers vs foetal livers fasting 24 h (I, J) where unpaired two-tailed Mann-Whitney test is used). Experiments in this figure were performed three times, except A, B one time

**Figure S9.**
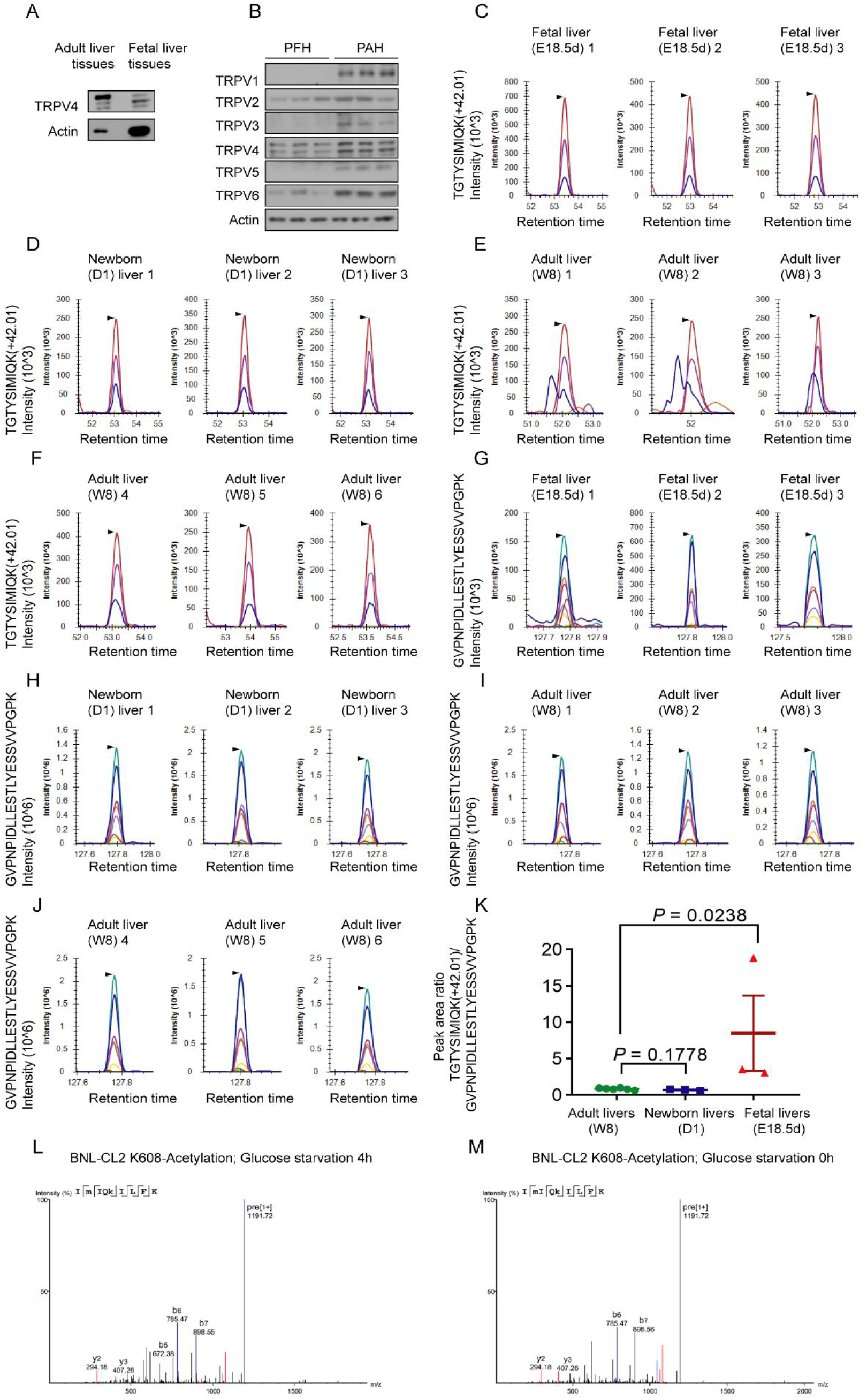
Levels of acetylated TRPV4-K608 are significantly higher in the foetal liver than in the adult liver, related to Figure 3 and Figure 4. A. The protein level of TRPV4 in foetal liver tissues is lower than those in adult liver tissue. B. The protein level of TRPV in PFH is lower than those in PAH. C-K. Levels of acetylated TRPV4-K608 are significantly higher in foetal liver than in adult liver, as determined by quantitative mass spectrometry analysis. Quantitative modification mass spectrometry detected a peptide segment containing K608-acetylation (TGTYSIMIQK (+42.01)) and a peptide segment (GVPNPIDLLESTLYESSVVPGPK). The peak area represents the relative abundance of each peptide segment in the sample. The peak area ratio of TGTYSIMIQK (+42.01)/GVPNPIDLLESTLYESSVVPGPK represented the relative level of acetylated trpv4 (at K608 site) in different samples (presented as mean ± SEM, n = 6 (adult livers of K; E, F, I, J), n = 3 (newborn livers of K; D, H), n=3 (foetal livers of K; C, G), with P values calculated using unpaired two-tailed Mann-Whitney test, except foetal livers vs newborn livers where unpaired two-tailed Student’s *t*-test is used. L, M. Qualitative mass spectrometry shows the presence of acetylation modification at the K608 site of TRPV4 in BNL-CL2. TRPV4 was enriched from BNL-CL2 cells stably expressing HA-tagged TRPV4 using immunoprecipitation. Experiments in this figure were performed three times.

**Figure S10.**
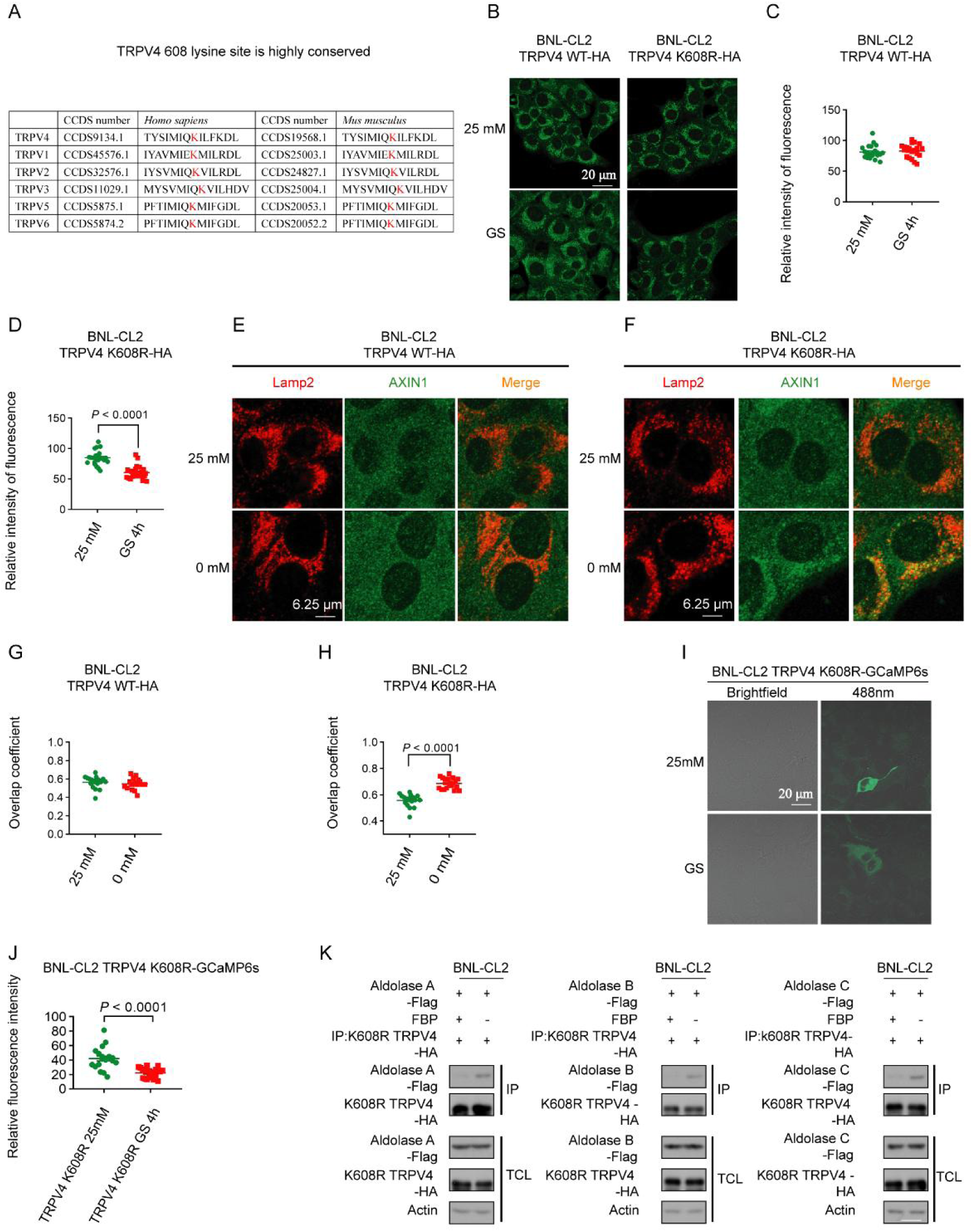
Deacetylation of TRPV during intrauterine development confers sensitivity to low glucose, related to Figure 4. A. The K608 residue is highly conserved among different species and different members of the TRPV family. B-D. v-ATPase inhibition can be observed in foetal hepatocytes expressing TRPV4-K608R in low glucose conditions, as seen in adult cells. The cells were incubated with 1 μM Lysosensor^TM^ Green DND-189 (Thermo L7535) for 30 min to detect the PH of lysosomes. The BNL-CL2 cells stably expressing HA-tagged TRPV4 WT (left panel of B, C) or HA-tagged TRPV4 K608R (right panel of B, D) were glucose-starved for 4 h. Representative images are shown in e, and the Mander’s overlap coefficients in C, D (presented as mean ± SEM, n = 21 for 25mM group of C, n=22 for GS 4 h group of C, n = 23 for 25mM group of D, n=24 for GS 4 h group of D, with P values calculated using unpaired two-tailed Student’s *t*-test). E-H. The lysosomal translocation of AXIN can be observed in foetal hepatocytes expressing TRPV4-K608R in low glucose conditions. The BNL-CL2 cells stably expressing HA-tagged TRPV4 WT (E, G) or HA-tagged TRPV4 K608R (F, H) were glucose-starved for 4 h. Representative images are shown in E, F and the Mander’s overlap coefficients in G, H (presented as mean ± SEM, n = 17 for 25 mM group of G, n = 16 for 0 mM of G, n = 17 for 25 mM group of H, n = 19 for 0 mM group of H, with P values calculated using unpaired two-tailed Student’s *t*-test). I, J. The mutation of K608 does not affect the basal activity of TRPV4, but renders TRPV4 able to be closed in foetal hepatocytes in low glucose conditions. BNL-CL2 stably expressing TRPV4 K608R-GCaMP6s was glucose-starved for 4 h. TRPV4-GCaMP6s were used to detect local calcium concentration, which can reflect whether TRPV4 is closed or not. The intensities of the indicator were then determined. The result showed that K608R-TRPV4, but not wildtype TRPV4, was closed in BNL-CL2 in low glucose conditions. Statistical analysis data are shown as mean ± SEM; 25 mM n = 14 for TRPV4 K608R group of C, n = 19 for TRPV4 K608R GS 4 h group of C; P values by unpaired two-tailed Mann-Whitney test (C). K. The mutation of K608 of TRPV4 does not change its affinity with the FBP-unoccupied aldolase. The BNL-CL2 (foetal hepatocytes) cells stably expressing HA-tagged TRPV4 K608R and Flag-tagged aldolase were glucose-starved for 4 h. Cells were then lysed, and the lysates were incubated with 200 μM FBP for 12 h. The interaction between aldolase and TRPV4 in the lysates was then determined by immunoprecipitation of HA-tag, followed by immunoblotting. Experiments in this figure were performed three times.

**Figure S11.**
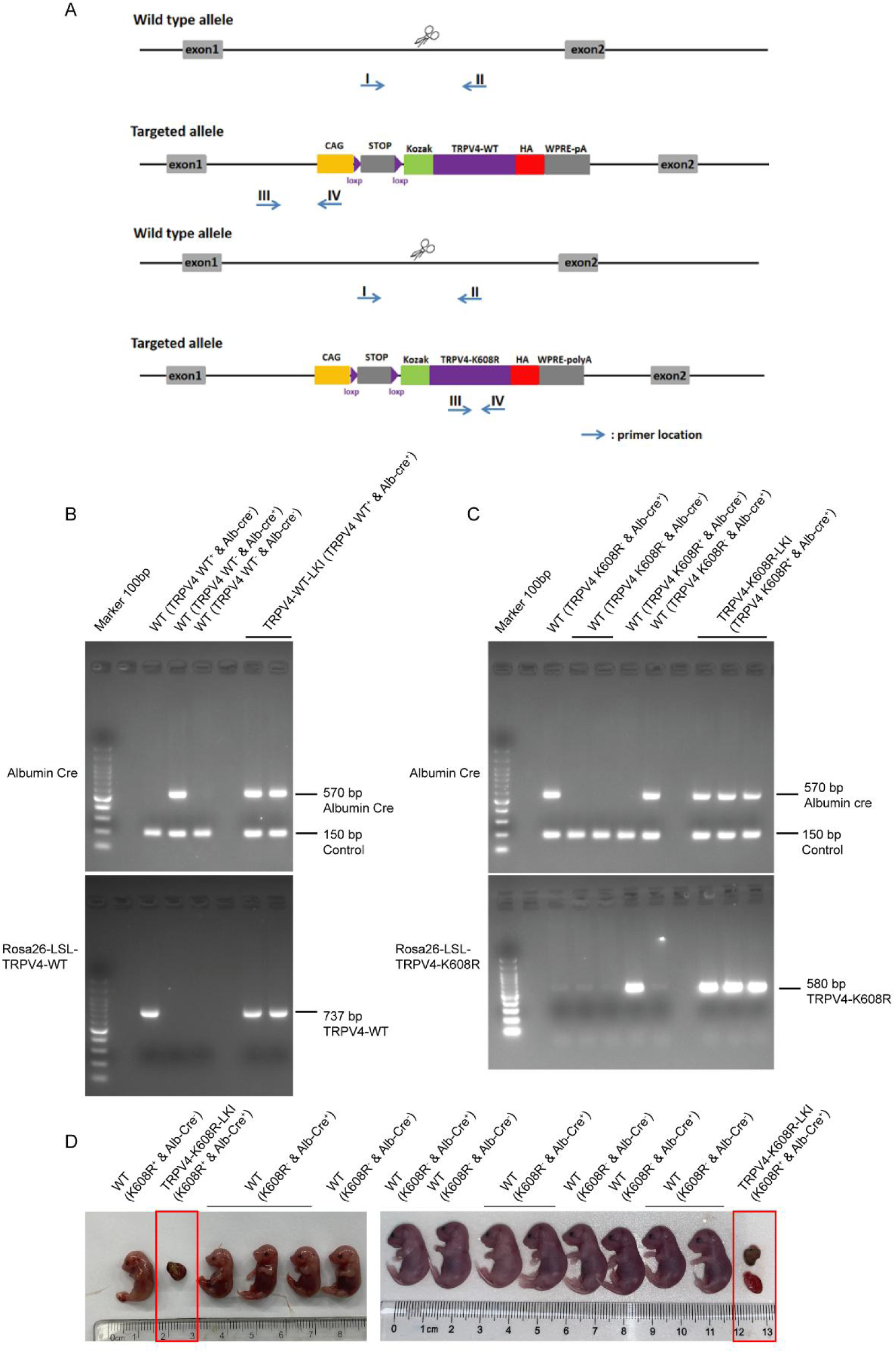
Phenotypes of foetal mice with TRPV4-K608R expression, related to Figure 4. A-C. Genotypes of foetal liver samples. D. Approximately 18% (2/11) of embryos with specific K608R-TRPV4 expression in the liver died during intrauterine development. Experiments in this figure were performed three times

**Figure S12.**
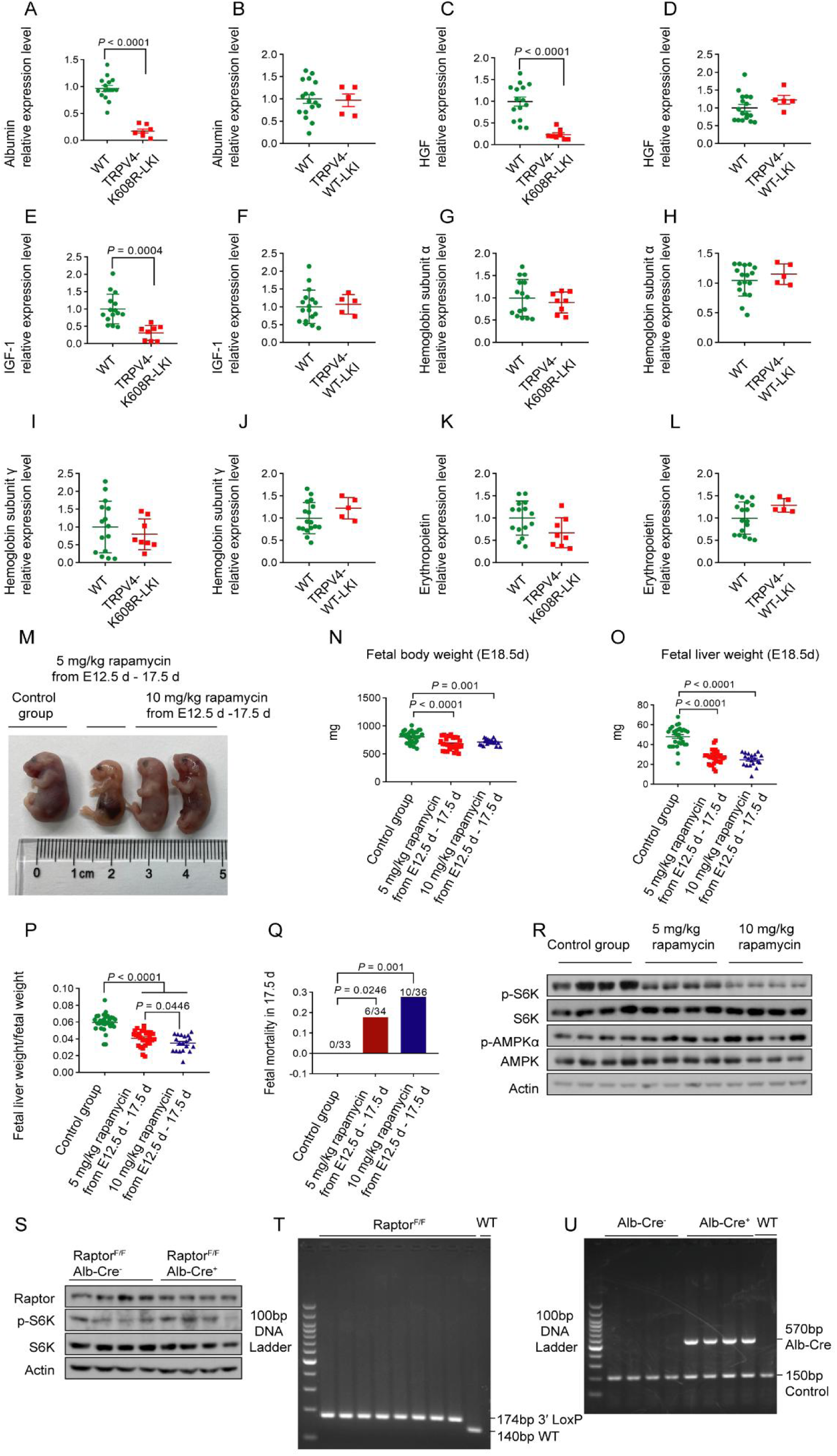
Rapamycin administration in wild-type embryos, which inhibits mTORC1 in foetal livers, recapitulates the phenotypes observed in the K608R-TRPV4-expressing embryos, related to Figure 4. A-L. TRPV4-K608R dominantly inhibits albumin, HGF, and IGF-1 in foetal livers but has little effect on hemoglobin and erythropoietin levels. Liver tissues collected from E18.5d foetal mice with liver-specific TRPV4-K608R (Figure 4F; A, C, E, G, I, K) or wildtype TRPV4 (Figure 4G; B, D, F, H, J, L) knocked-in were lysed, followed by determination of albumin, HGF, IGF-1, hemoglobin α/γ and erythropoietin levels by immunoblotting. Statistical analysis data are shown as mean ± SEM, n = 15 for WT group of A, C, E, G, I, K; n = 8 for TRPV4-K608R-LKI group of A, C, E, G, I, K; n = 17 for WT group of B, D, F, H, J, L; n = 5 for TRPV4-WT-LKI group of B, D, F, H, J, L and P values by unpaired two-tailed Student’s *t*-test (A, B, D-L) and by unpaired two-tailed Mann-Whitney test (c). M-Q. Rapamycin administration in wild-type embryos resembles the phenotypes observed in the K608R-TRPV4-expressing embryos. Rapamycin leads to a decrease in liver weight (O), foetal liver weight/foetal weight (P), and an increase in intrauterine mortality rate (Q) in foetal mice. Statistical analysis data are shown as mean ± SEM, control group n = 30 for control group of N-P, n = 28 for 5 mg/kg rapamycin group N-P, n = 19 for 10 mg/kg rapamycin group of N-P; n = 33 for control group of Q, n = 34 for 5mg/kg rapamycin group of Q, n = 36 for 10 mg/kg rapamycin group of Q. P values by Fisher’s exact test (Q) and unpaired two-tailed Student’s *t*-test, except control group vs 10 mg/kg rapamycin group (N), 5 mg/kg rapamycin group vs 10 mg/kg rapamycin group (N) where unpaired two-tailed Mann-Whitney test is used. R. Rapamycin administration in wild-type embryos inhibits mTORC1 in foetal livers, as assessed by the p-S6K levels via immunobloting. S. Blockage of the mTORC1 signaling by knocking out Raptor in the foetal liver is unsuccessful. The protein levels of Raptor remained mostly unaffected in the foetal liver (Raptor^f/f^ & Alb-cre^−^ vs Raptor^f/f^ & Alb-cre^+^). T, U. Genotyping of Alb-cre^−^, Raptor ^f/f^ and Alb-cre^+^, Raptor ^f/f^ foetal mice. Alb-cre^+^, Raptor^f/-^ male mice were mated with Raptor ^f/f^ female mice to obtain Raptor^f/f^ & Alb-cre^−^, Raptor^f/f^ & Alb-cre^+^ foetal mice. Experiments in this figure were performed three times

**Figure S13.**
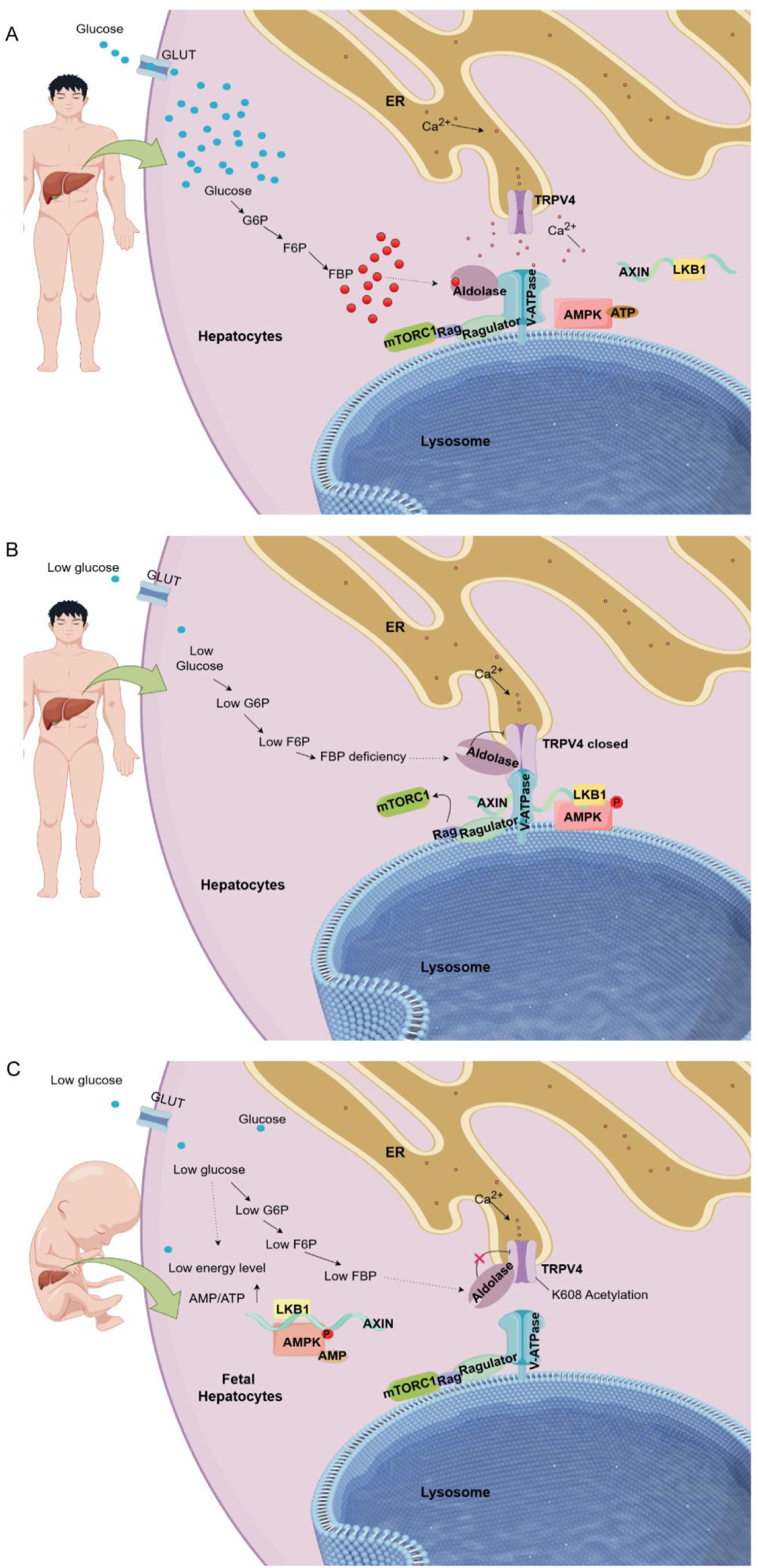
Models of the glucose-sensing pathway in adult and foetal livers. These simplified models illustrate how the absence of glucose and FBP is sensed and relayed from aldolase to TRPVs (represented by TRPV4), then to v-ATPase, ultimately leading to the formation of an AXIN-based complex on the lysosome for inhibition of mTORC1. A. Adult livers with abundant glucose: When glucose supply is abundant, the FBP-bound aldolase is associated with v-ATPase, which is crucial for maintaining its activity. The TRPV4 channel releases Ca²⁺ locally at the ER-lysosome contact site. Here, v-ATPase remains active, and mTORC1 is anchored to the lysosome and maintains activity. B. Adult livers with insufficient glucose: When glucose levels are low, FBP-unbound aldolase interacts with and inhibits the adjacent TRPV4 on the ER. As local Ca²⁺ dissipates, TRPV4 becomes accessible to v-ATPase, leading to a reconfiguration of the complex into an “aldolase:TRPV:v-ATPase” formation. Subsequently, v-ATPase is inhibited, and allows the binding of AXIN and causes mTORC1 to dissociate from the lysosome. In addition, LKB1, brought about by AXIN, activates AMPK on the lysosome. C. Foetal livers with insufficient glucose: In foetal livers with low glucose, ATP levels drop, resulting in an increase in AMP/ATP ratio, which allosterically activates AMPK. The interaction between FBP-unbound aldolase and TRPV4 occurs, but it does not result in inhibition due to acetylation at the K608 site of TRPV4. As a result, v-ATPase remains active, and mTORC1 stays anchored to the lysosome, thereby maintaining its activity.

**Table S1.** The number of surviving offspring from transgenic mice with liver-specific expression of K608R-TRPV4 or TRPV4 wide-type (induced by *Alb-Cre*).

| Genotype | Number of mice |
| --- | --- |
| *TRPV4-K608R<sup>-</sup> & Alb-Cre<sup>-</sup>* | 13 |
| *TRPV4-K608R<sup>+</sup> & Alb-Cre<sup>-</sup>* | 18 |
| *TRPV4-K608R<sup>-</sup> & Alb-Cre<sup>+</sup>* | 15 |
| *TRPV4-K608R<sup>+</sup> & Alb-Cre<sup>+</sup>* | 0 |
| *TRPV4-WT<sup>-</sup> & Alb-Cre<sup>-</sup>* | 13 |
| *TRPV4-WT<sup>+</sup> & Alb-Cre<sup>-</sup>* | 13 |
| *TRPV4-WT<sup>-</sup> & Alb-Cre<sup>+</sup>* | 11 |
| *TRPV4-WT<sup>+</sup> & Alb-Cre<sup>+</sup>* | 12 |

## STAR★METHODS

### Data reporting

The chosen sample sizes were similar to those used in this field: n = 4-21 samples were used to evaluate the levels of blood glucose in serum^12,58,59^; n = 4-21 samples were used to evaluate the levels of metabolites in cells^12,60,61^; n = 7-27 samples were used to evaluate the levels of metabolites in tissues^61–63^; n = 7-11 samples were used to evaluate the levels of adenylates in tissues^64^; n = 3-7 samples were used to determine the mRNA levels of a specific gene^11,65,66^; n = 3-6 samples were used to determine the expression levels and phosphorylation levels of a specific protein^11,60^; n = 16-62 cells were used to determine the Mander’s overlap coefficients of mTORC1 or AXIN with LAMP2, or the intensities of Lysosensor dye^10,11,13^. No statistical methods were used to predetermine the sample size. All experimental findings were repeated as stated in the figure legends, and all additional replication attempts were successful. For animal experiments, mice were housed under the same conditions. For cell experiments, cells of each genotype were cultured in the same CO2 incubator and seeded in parallel. Each experiment was designed and performed with proper controls, and samples for comparison were collected and analysed under the same conditions. Randomisation was applied wherever possible. For example, during mass spectrum (MS) analyses, samples were processed and subjected to MS in random orders. For animal experiments, sex-matched, age-matched adult mice were randomly assigned to the control group, fasting group or rapamycin treatment group. Foetal mice were not differentiated by sex. In cell experiments, cells of each genotype were seeded in parallel and randomly assigned to different treatments. Otherwise, randomisation was not performed. For example, when performing immunoblotting, samples needed to be loaded in a specific order to generate the final figures. Blinding was applied wherever possible. For example, samples and cages during sample collection and processing were labelled with code names that were later revealed by the individual who picked and treated animals or cells, but did not participate in sample collection and processing, until the outcome was assessed. Similarly, during microscopy data collection and statistical analyses, the fields of view were chosen on a random basis, often performed by different operators, preventing potentially biased selection for desired phenotypes.

### Experimental model and subject details

#### Studies in animals

Wild-type C57BL/6J (#000664), *Raptor*^F/F^ (#013191, generated as in ref ^50^), and *Alb*-*Cre* mice (#003574) were obtained from the Jackson Laboratory. *Raptor-LKO* mice were generated by crossing *Raptor*^F/F^ mice with *Alb*-*Cre* mice. Genotyping primers for *Raptor* and *Alb*-*Cre* were supplied in Key Resources Table. For identification *Raptor*^F/F^ mice, the wild-type allele was present, the PCR reaction generated a 140-bp product, while, the presence of the 3’-LoxP element resulted in a 174-bp amplicon. The PCR reaction generated an amplicon of approximately 570 bp, indicating the presence of *Alb*-*Cre*. To ensure the reliability of the PCR system, a fragment of approximately 150 bp was used as a positive control. The TRPV4 wild-type (N7-100246) and TRPV4-K608R (N7-100247) knock-in mice were customised by Shanghai Model Organisms Center, Inc by introducing the the CAG-LSL-*TRPV4*-WT (or K608R)-HA-WPRE-pA cassette into the recipient C57BL/6J mice at the *Rosa26* locus, following the protocol as described previously^66^ (see details in the Supplementary Information, Pages 10 to 26). Briefly, to create these mice, Cas9 mRNA and gRNA were first produced by in vitro transcription. Then, a donor vector containing a 3.3-kb 5’ homologous arm, the knock-in sequence, and a 3.3-kb 3’ homologous arm was constructed by the in-fusion cloning strategy. Next, the Cas9 mRNA, gRNA, and the donor vector were mixed, and micro-injected into fertilised eggs of C57BL/6J mice, and the F0 mice with successfully knock-in, after identifying by PCR and sequencing, were crossed with wild-type C57BL/6J mice to generate the F1 mice. The homozygous *TRPV4*^F/F^ and *TRPV4-K608R^F/F^* mice resulting from this process were viable, fertile, and normal in size with no apparent phenotypic abnormalities, and were crossed with *Alb*-*Cre* mice to generate the *TRPV4*-LKI and *TRPV4*-*K608R*-LKI mice. Primers for genotyping *TRPV4-K608R* and *TRPV4-WT* were supplied in Key Resources Table. The PCR reaction generated an amplicon of 967 bp when the wild-type allele site was present, and of 580 bp when the *TRPV4-K608R* was present. The PCR reaction generated an amplicon of 967 bp when the wild-type allele site was present and of 737 bp when the *TRPV4-WT* was present. *Alb*-*Cre* was genotyped as in the *Raptor*-LKO mice.

The genotyping process was carried out as follows: Initially, a 1.5-mm length of the mouse’s tail was cut using a surgical blade (to determine the genotype of foetal mice, a limb or entire tail of the mice was cut), and placed in a DNA-free, 1.5-ml EP tube. Some 100 μl of 25 mM NaOH solution was then added to the 1.5-ml EP tube, followed by centrifuge at 3000*g* for 1 min to ensure that the tail tissues were fully immersed in the NaOH solution, and then heated in a metal bath at 100 °C for 1 h. After heating, the tube was centrifuged (3000*g*, 1 min) to ensure that all liquids or tissues were pelleted at the bottom of the tube, and the mixture was then neutralised by addition of 100 μl of 40 mM Tris-HCl (pH 8.0) solution. The PCR reaction system consists of 10 μl of 2× Taq PCR Master Mix, 1 μl of mouse tail DNA sample, 0.5 μl (10 µM) of forward primer, 0.5 μl (10 µM) of reverse primer, and 8 μl of ddH_2_O. The programs were set as follows:

a. For *Alb*-*Cre*:

- Initial denaturation at 95 °C for 3 min
- 35 cycles of denaturation at 95 °C for 30 s, annealing at 58 °C for 40 s, extension at 72 °C for 30 s
- Final extension at 72°C for 5 min
- PCR products were analysed using a 2% DNA agarose gel.
b. For *Raptor*^F/F^:

- Initial denaturation at 95 °C for 5 min
- 10 cycles of denaturation at 95 °C for 30 s, annealing starting at 65 °C with a decrease of 0.5 °C per cycle, extension at 72 °C for 30 s
- Followed by 28 cycles of denaturation at 95 °C for 30 s, annealing at 60 °C for 30 s, extension at 72 °C for 30 s
- Final extension at 72 °C for 5 min
- PCR products were identified using a 3% DNA agarose gel.
c. For *TRPV4*^F/F^ and *TRPV4-K608R*^F/F^:

- Initial denaturation at 95 °C for 5 min
- 35 cycles of denaturation at 95 °C for 30 s, annealing at 60 °C for 30 s, extension at 72 °C for 30 s
- Final extension at 72 °C for 5 min
- PCR products were identified using a 2% DNA agarose gel.

### Mouse fasting treatments

Protocols for all mouse experiments were approved by the Institutional Animal Care and the Use Committee of Xiamen University (XMULAC20210014). Unless stated otherwise, mice were housed with free access to water and a standard diet (65% carbohydrates, 11% fat, 24% protein) under specific pathogen-free conditions. The lighting was on from 8:00 to 20:00, with the temperature maintained at 21-24 °C and humidity at 40-70%. Unless stated otherwise, male and female mice were randomly selected in this study at an approximately 1:1 ratio. Mouse used for collection of the liver tissues or isolation of primary hepatocytes were also selected randomly, without distinction of sex. Mice were individually caged for one week before each treatment. For fasting experiments, the mice were placed in a new cage at 15:00 p.m., and the experiment was conducted the next day at 15:00 p.m.

### Rapamycin treatment of mice

The E11.5 C57BL/6 dams were randomly divided into three groups. The dams received intraperitoneal injections of either corn oil, 5 mg/kg rapamycin (MedChemExpress, cat. HY-10219; dissolved in anhydrous ethanol with a stock solution of 5 mg/ml, and freshly diluted to a working concentration of 0.5 mg/ml by diluting the stock solution with 9 volumes of corn oil), or 10 mg/kg rapamycin once a day from E12.5 to E17.5. Note that rapamycin should be injected slowly due to the high viscosity of corn oil to avoid generating bubbles. In addition, extra caution should be taken when administering drugs as the gestational age increases to avoid puncturing the amniotic sac with the needle when the abdominal bulge of pregnant mice becomes more pronounced. At 18.5 days of pregnancy, the pregnant mice were anesthetized with isoflurane, and the foetal livers, hearts, lungs, and kidneys were excised and weighed, or quickly lysed for the determination of the levels of p-AMPKα, AMPKα, p-S6K, S6K, Raptor, and β-actin by immunoblotting.

### Primary hepatocytes

Mouse adult primary hepatocytes were isolated with a modified two-step perfusion method, as described previously^66,67^. Briefly, mice were anaesthetised before the isolation, followed by the insertion of a 0.72 × 19 mm intravenous catheter into the post-cava. After cutting off the portal vein, mice were perfused with 50 ml of liver perfusion medium (A buffer; prepared by mixing 49 ml of Kreb buffer with 1 ml of 50 mM EGTA buffer) at a rate of 5 ml/min, followed by 50 ml of liver digest buffer (B buffer; by dissolving 25 mg Type IV collagenase (Sigma, cat. C5138) in a solution made by mixing 49 ml of Kreb buffer with 1 ml of 250 mM CaCl_2_ (Sango Biotech, cat. A610050-0500)) at a rate of 2.5 ml/min. The Kreb buffer (pH 7.4) was prepared by dissolving 7 g of NaCl (Sango Biotech, cat. A610476), 2 g of NaHCO_3_ (Sango Biotech, cat. A500873-0500), 1.1915 g of HEPES (Gibco, cat. 15630130), and 1 g Glucose (Sigma, cat. G7021-1Kg), in a solution containing 10 ml of Solution C (180 mM KCl (Sango Biotech, cat. A100395), 120 mM MgSO_4_·7H_2_O (Sigma, cat. M1880-500G), 120 mM KH_2_PO_4_ (Sango Biotech, cat. A100781-0100)) and 990 ml deionized water, followed by filtration with 0.22 μm pore size filters (Sartorius, cat. 16541K). The digested liver was then briefly rinsed with PBS and dissected by gently tearing apart the Glisson’s capsule with two sterilised, needle-pointed tweezers on a 6-cm dish containing 3 ml of PBS. The dispersed cells were mixed with 10 ml of cold William’s medium E plus 10% foetal bovine serum (FBS) and filtered by passing through a 100-μm cell strainer (BD Falcon). Cells were then centrifuged at 50*g* at 4 °C for 2 min, followed by washing twice with 10 ml of ice-cold William’s medium E plus 10% FBS (Gibco, cat. 30044333). Cells were then immediately plated (at 60-70% confluence) in collagen-coated 6-well plates in William’s medium E plus 10% FBS, 100 IU penicillin and 100 mg/ml streptomycin. After 4 h of attachment, the medium was replaced with fresh Dulbecco’s modified Eagle’s medium with 10% BSA for another 48 h before further use.

Mouse foetal primary hepatocytes were isolated following the procedures described previously^68^, with minor modifications:

a. Preparation of HEPES buffer:

- HEPES buffer was prepared by adding 8 g NaCl, 0.2 g KCl, 0.1 g Na_2_HPO_4_·12H_2_O, and 10 ml of 1 M HEPES solution to 750 ml of autoclaved distilled water, followed by adjusting the pH to 7.65 and the volume to 1 l. The solution was filtered with a 0.22-μm filter and were stored at 4 °C.
b. Preparation of basal culture media and complete culture media:

- Basal culture media: by addion of 50 ml of FBS, 5 ml of L-glutamine (200 mM stock solution), and 5 ml of the antibiotic (streptomycin 10000μg/ml, penicillin 10000IU/ml) solution to William’s E medium to a final volume of 500 ml.
- Complete culture media: by addition of HGF (final concentration 25 ng/ml), EGF (25 ng/ml), Insulin (5 μg/ml), Hydrocortisone (0.5 μM), and Dexamethasone (0.1 μM) to the basal culture media.
c. Preparation of Liver Dissociation Solution:

- Dissolved the lyophilised liberase (5 mg of collagenase I) in 2 ml of sterile water (to get a 13 U/ml enzyme solution).
d. Isolation of foetal livers from anesthetised pregnant mice:

- Pregnant mice were anesthetised and the foetal liver tissues were excised using sterilised tweezers and scissors. Some 300 mg of liver tissues were then placed in cold HEPES buffer (4 °C) in a biosafety cabinet.
e. Preparation of foetal liver tissue:

- The connective tissue and the gastrointestinal tract attached to the foetal liver tissue were removed using sterile forceps. The liver was then transferred to the clean, cool HEPES buffer and was rinsed twice by the cool HEPES buffer, followed by sliced into small pieces by sterilised blades, and then incubated in 10 ml of liberase-HEPES-calcium buffer for 5 min at 37 °C. The homogenates were then aspirated using a 10-ml glass pipette once per minute. Note that any pipetting (e.g., using the 1-ml pipette) should be avoided, as it will severely decrease cell viability. After 5 min of incubation, tissues were allowed for another 5 min of incubation at 37 °C, followed by mixing with 25 ml of basal media to terminate the digestion.
f. Cell isolation and culture:

- The homogenates were then filtrated using a 70-µm filter (Corning. cat. 352350). After transferring the filtrate to a sterile, 50-ml centrifuge tube, cells in the filtrate were collected by centrifuged at 1,000*g* for 2 min at 4 °C. Cells were then resuspended the with 25 ml of basal medium, followed by wash with the complete culture medium, and then resuspended in complete culture medium for culturing.

Note that in this study, the isolated foetal liver cells were not further purified by magnetic bead sorting^68^, as we did not observe any significant difference in the context of this study. The primary foetal hepatocytes were cultured for 2-3 days, before subjected to further immunofluorescence, immunoblotting, and metabolite analysis.

### Clinical specimens

Foetal liver tissues (n = 12) were collected from women who voluntarily terminated the pregnancy at the Obstetrics and Gynecology Hospital of Fudan University. Adult liver tissues (n = 10) were the adjacent tissue of benign lesions from surgery at Fudan University Shanghai Cancer Center. The procedures involving adult participants (adult liver tissues) approved by the Ethics Committee of Fudan University Shanghai Cancer Center (050432-4-2108*). The procedures involving pregnant participants (foetal liver tissues) were reviewed and approved by the Ethics Committee of Obstetrics and Gynecology Hospital of Fudan University (2023-66). Written informed consents were obtained from all participants. All tissues were collected in sterile tubes and quickly stored at −80 °C at the surgical site.

Tissue lysates were prepared as followed: Appropriate amounts of tissue were cut (30-50mg), and 10 μl/mg of protein lysis solution, pre-mixed with phosphatase inhibitors, was added to 1.5 ml EP tube containing the tissues. A handheld homogenizer was used to break the tissue, and an ultrasound was applied to further lyse the cells. Centrifuged at 14,000*g* for 10 min, and the supernatant was collected and mixed with an equal volume of 2 × SDS solution. The mixture was well mixed and the proteins were denatured at 100 ° C for 15 min. The levels of p-AMPKα, AMPKα, p-S6K, S6K and β-actin proteins were detected in the clinical specimens by western blot. Based on the levels of p-AMPKα, the samples were categorized into two groups: AMPK-relatively inactive and AMPK-relatively active. The ratios of p-S6K/S6K and p-AMPKα/AMPKα in each group and their correlations were analysed.

### Cell lines

MEFs cells were established as described previously^64^. BNL-CL2, HEK293T and MEF were maintained in Dulbecco’s modified Eagle’s medium (Gibco, cat. 11960044) supplemented with 10% FBS (Gibco, cat. 10099158), 100 IU penicillin, 100 mg/ml streptomycin at 37 °C in a humidified incubator containing 5% CO2. The AML12 cultured in DMEM/F12 (Gibco, cat. 11320033) supplemented with 10% FBS, 100 IU penicillin, 100 mg/ml streptomycin, 4 ng/ml dexamethasone and ITS Liquid media supplement (Sigma, cat. I3146). All cell lines were verified to be free of mycoplasma contamination. Cells were plated in 6-well plates one day before the experiments. On the next day, cells were rinsed once with PBS, and incubated with amino-acid-free EBSS and/or glucose-free DMEM supplemented with 10% FBS and relevant concentration drugs (Rapamycin, BCTC, Concanamycin A etc.) for the indicated time. 1L Amino acid-free EBSS buffer (PH 7.4): 0.2 g CaCl_2_ (Sango Biotech, cat. A610050-0500), 0.2 g MgSO_4_.7H_2_O (Sigma, cat. M1880-500G), 0.4 g KCl (Sango Biotech, cat. A100395), 2.2 g NaHCO_3_ (Sango Biotech, cat. A500873-0500), 6.8 g NaCl (Sango Biotech, cat. A610476), 0.14 g NaH_2_PO_4_.H_2_O (Sango Biotech, cat. A100823) in 1000 ml deionized water. The dissolved EBSS liquid was filtered and sterilized using a filter with a pore size of 0.22 µm. Taking a 6-well plate as an example, to study the effect of amino acids on mTORC1 and AMPK in foetal hepatocytes, first washed the cells once with PBS, added 2 ml of EBSS solution to each well, and then added the corresponding reagents in sequence, +G+A (25 mM glucose (Sigma, cat. G7021-1Kg) + 1 × MEM Amino Acids (Gibco, cat.11130-051) + 1 × MEM Non-essential Amino Acid solution (Sigma, cat. M7145), −G-A (No glucose, no amino acid), +G-A (25mM glucose, no amino acid), −G+A (No glucose, + 1 × MEM Amino Acid (Gibco, cat. 11130-051) + 1 × MEM Non-essential Amino Acid solution. Taking the 6-well plate as an example, if only the effect of glucose starvation on AMPK and mTORC1 in foetal liver was studied, the cells were washed once with PBS, and then 2ml of glucose-free DMEM medium containing 10% FBS was added to each well, followed by the corresponding reagents (Different concentrations of glucose). The glucose storage solution was 2.5M (Sigma, cat. G7021-1Kg), with working concentrations of 0 mM, 1 mM, 3 mM, 5 mM, 10 mM and 25 mM. Taking a 6-well plate as an example, to study the effects of growth factors on AMPK and mTORC1 in foetal hepatocytes, the cells were first treated overnight with high-glucose DMEM without serum, and then 2ml of glucose-free DMEM medium without serum was added to each well. Finally, corresponding reagents were added. +G+I (5 µg/ml insulin (Procell, cat. PB180432) + 25 mM Glucose), +G-I (no insulin + 25 mM glucose), −G+I (no glucose + 5 µg/ml insulin), −G-I (no insulin, no glucose). Then, the cells were lysed and relevant proteins were detected by western blot.

### Plasmids

The restriction enzyme cleavage sites of pBOBI are BamHI and XhoI. The WT human *TRPV4* sequence was based on CCDS9134.1. Point mutations of *TRPV4* were performed by PCR-based site-directed mutagenesis using PrimeSTAR HS polymerase (Takara, cat. R010A). The construction method for the deacetylation point mutation involves mutating the lysine at the corresponding site to arginine. The full sequence of TRPV4 WT, TRPV4 K608R, GCaMP6s were showed in the Supplementary Information files. Purified PCR products and pBOBI plasmid (digested with BamHI and XhoI) were incubated with Exonuclease III and the relevant buffer (NEW ENGLAND Biolabs, cat. M0206L) on ice for 1 hour. Then, added 1 μL of 0.5 M EDTA to terminate the enzyme digestion reaction, and heated at 65°C for 10 min to inactivate the enzyme. After cooling, the reaction products were added to competent cells of the STABLE 3 *Escherichia coli* on ice for 30 min. Performed a 42°C thermal activation for 90s. Cultivated STABLE 3 *E. coli* in LB solid culture medium with ampicillin resistance. After incubating at 37 °C for 16 h, removed the solid culture medium. Selected monoclonal colonies and placed them in 25 ml of ampicillin-resistant liquid LB medium. Incubated the liquid LB culture medium with colonies on a 37 °C shaker for 20 h. Finally, lysed the bacteria and extracted the plasmid. All mutation proteins were stably expressed in BNL CL.2 with pBOBI for lentivirus packaging. The methods for constructing stable gene expression cell lines were as follows: (1) Mixed 300 μl OPTI-MEM (Gibco, cat. 31985070), 1.5 μg plasmid, and 1.5 μg Lenti-vector (pMOL: VSV-G: REV=5:3:2) thoroughly. (2) Added 7 μl of Lip2000 Transfection Reagent (Biosharp, cat. BL623B) and mix thoroughly. (3) Kept the mixture at room temperature for 30 min. (4) Digested a 10 cm dish of 293T cells (cell density 100%) and resuspended with 900 μl of DMEM medium. (5) Plated the plasmid mixture and cells in proportion on a 6-well plate. Each well: 700 μl nutrition rich DMEM culture medium (Dulbecco’s modified Eagle’s medium (Gibco, cat. 11960044) + 10% FBS (Gibco, cat. 10099158) + 1 × MEN Non-essential Amino Acid Solution (Sigma, cat. M7145-100mL) + 1 × GlutaMAX^TM^ (Gibco, cat. 35050061) + 1 mM Sodium pyruvate solution (Sigma, cat. S8636-100ML) + 100 IU penicillin, 100 mg ml^−1^ streptomycin) + 160 μl 293T suspension + 300 μl plasmid mixture. (6) Placed the cells in a 37°C incubator for cultivation, and changed the medium after 10 h. Added 2 ml of nutrient-rich DMEM culture medium to each well. (7) Collected the culture medium 48 h later, centrifuged and took the supernatant, which was the Lenti-virus medium used to infect cells. (8) Added 10 mg/ml polybrene (Hexadimethrine Bromide, Sigma, cat. H9268) to the Lenti-virus solution in a 1:1000 ratio. Mixed thoroughly and centrifuged. (9) Digested BNL-CL2 and resuspended them, inoculated 1.67 x 10^5^ cells per well in a 6-well plate. Cultivated BNL-CL2 in the incubator for 30 min, then changed the culture medium with 500 μl DMEM medium (Gibco, cat. 11960044) supplemented with 10% FBS (Gibco, cat. 10099158), 100 IU penicillin, 100 mg/ml streptomycin + 2 ml Lenti-virus solution. (10) Placed the 6-well plate in a horizontal centrifuge and centrifuged at 30°C and 2500 rpm for 30 min. (11) After centrifugation, placed the 6-well plate in a 37°C incubator for 48 h. (12) Detected plasmid expression through immunoblotting.

### Protein expression in cell lines

This study involved the over-expression of target genes in cells, including over-expression of TRPV4-WT, TRPV4-K608R, aldolase A, aldolase B, aldolase C, TRPV4 WT-GCaMP6s, and TRPV4 K608R-GCaMP6s in BNL-CL2, and over-expression of TRPV4 WT-GCaMP6s in AML12 and 293T cells. In Figure 3a, BNL-CL2 over-expressed TRPV4 WT-HA and aldolase A/B/C-Flag, the specific experimental steps were as follows: Mixed pBOBI homo sapiens TRPV4-WT-HA (N-terminal tag) 1.5 μg, pBOBI homo sapiens aldolase A/B/C-Flag (N-terminal tag) 1.5 μg, lentiviral plasmid (pMDL:VSVG:REV=5:3:2) 1.5 μg, and 300 μl OPTI-MEM well in 1.5 ml sterile EP tube by vortexing and used a centrifuge (3000*g* for 1 min) to pellet the liquid at the bottom of the tube. Then added 7 μl Lipo2000 to the 1.5ml EP tube, mixed well by vortexing, and centrifuged. Placed the 1.5 ml EP tube on a clean bench and let it stand at room temperature for at least 30 min. Prepared 293T grown to 80-90% confluence in a 10cm dish, gently removed the cell culture medium, rinsed the culture dish with 10ml PBS buffer, added 1ml of commercial trypsin and placed it in a 37 °C incubator until the cells were completely digested (approximately 1 min), then added 1ml of complete culture medium (10% FBS, DMEM high glucose medium, penicillin streptomycin) to stop the digestion process. Transferred the cell suspension to a 2ml sterile cryovial and centrifuged (1000*g*, 3 min). Removed the supernatant and resuspended the 293T cells using 1ml of nutrient-rich DMEM culture medium (Dulbecco’s modified Eagle’s medium (Gibco, cat.11960044) + 10% FBS (Gibco, cat.10099158) + 1 × MEN Non-essential Amino Acid Solution (Sigma, cat. M7145-100mL) + 1 × GlutaMAX^TM^ (Gibco, cat. 35050061) + 1 mM Sodium pyruvate solution (Sigma, cat. S8636-100ML) + 100 IU penicillin, 100 mg/ml streptomycin). Plated the plasmid mixture and cells in proportion on a 6-well plate. Each well: 700 μl nutrition rich DMEM culture medium + 160 μl 293T suspension + 300 μl plasmid mixture. Placed the 6-well plate in a 37 °C incubator for cultivation, changed the medium after 10 h, and added 2 ml fresh nutrition-rich DMEM culture medium to each well. Collected the culture medium 48 h later, centrifuge and take the supernatant, which was the Lenti-virus medium used to infect BNL-CL2 cells. Added 10 mg/ml polybrene (Hexadimethrine Bromide, Sigma, cat.H9268) to the Lenti-virus solution in a 1:1000 ratio. Mixed thoroughly and centrifuged. Digest BNL-CL2 cells and resuspend them, inoculated 1.67 × 10^5^ cells per well in a 6-well plate. Cultivated BNL-CL2 cells in the incubator for 30 min (at this point, the cells should be just adhering to the wall, still maintaining a circular shape, and not yet fully spread), then changed the culture medium by 500 μl DMEM medium (Gibco, cat.11960044) supplemented with 10% FBS (Gibco, cat.10099158), 100 IU penicillin, 100 mg/ml streptomycin + 2 ml Lenti-virus solution. Placed the 6-well plate in a horizontal centrifuge and centrifuged at 30°C and 2500 rpm for 30 min. After centrifugation, placed the 6-well plate in a 37 °C incubator for 48 h until the cells reach 70-80% confluence. Subsequently, transferred the BNL-CL2 cells infected with the lenti-virus to a 6-cm dish and continued culturing until the cells reach 70-80% confluence. Selected half of the cells in the 6-cm dish to lyse and extracted protein, and used mouse anti-Flag (cat. F1804-5MG, 1:1000 for IB) and mouse anti-HA (cat.sc-7392, 1:1000 for IB) to detect the presence of target protein bands. If both TRPV4 WT-HA and aldolase A/B/C-Flag bands were detected in the cell lysate, it proved the successful over-expression of TRPV4 WT and aldolase A/B/C in BNL-CL2 cells. The remaining BNL-CL2 cells infected with lenti-virus in the previous 6-cm dish can be used to complete the Figure 3a experiment. Other over-expressed target genes adopt a similar approach, with differences in the type of infected cells (BNL-CL2, 293T, AML12) and the plasmids of the target genes.

### Cell lysates and immunoprecipitation

Cells were rinsed once with PBS and lysed in ice-cold lysis buffer (40 mM HEPES [pH 7.4], 2 mM EDTA or 5mM MgCl2, 10 mM pyrophosphate, 10 mM glycerophosphate, 0.3% CHAPS, or 1% Trition X-100) containing one tablet of EDTA-free protease inhibitors (Roche) per 25 ml. The soluble fractions of cell lysates were isolated by centrifugation at 13,000 rpm for 10 min by centrifugation.

For immunoprecipitations, primary antibodies and protein A/G beads were added to the lysates and incubated with rotation for overnight at 4°C. Immunoprecipitates were washed three times with fresh lysis buffer. Immunoprecipitated proteins were denatured with a moderate dose of 2× SDS buffer and boiling for 15 min, separated by 8%-15% SDS-PAGE, and analyzed by immunoblotting. For Flag purifications, Flag M2 affinity gel was washed with lysis buffer three times. Then, 10 μl of beads in 50% slurry was added to pre-cleared cell lysates and incubated with rotation for 24 h at 4°C. Finally, the beads were washed three times with lysis buffer. Immunoprecipitated proteins were denatured by the addition of 2× SDS buffer and boiling for 15 min.

### Qualitative mass spectrometry analyses

Fifty 10-cm dishes of BNL-CL2 (grown to 80% confluence) stably expressing HA-tagged TRPV4 were lysed, and anti-HA immunoprecipitations were performed in lysis buffer (20 mM Tris-HCl, pH 7.5, 150 mM NaCl, 1 mM EDTA, 1 mM EGTA, 1% (v/w) Triton X-100, 2.5 mM sodium pyrophosphate, 1 mM β-glycerophosphate, with protease inhibitor cocktail). The specific method was as follows: Ten plates of cells were taken out from the incubator each time, and one experimenter used a vacuum pump to remove the culture medium from the culture dish (do not completely remove, leave 0.5ml of culture medium). Another experimenter used a clean cell scraper to scrape off the adherent cells as quickly as possible, and used a 1 ml pipette to transfer the scraped cell suspension to a sterile 50 ml centrifuge tube until all the cells (from 50 × 10 cm culture dishes) were transferred to the 50 ml centrifuge tube. The cell collection process was carried out at room temperature. Subsequently, the 50 ml centrifuge tube containing cells was centrifuged using a horizontal centrifuge until the cells settled at the bottom of the tube (1000*g*, 2 min, 25°C). The supernatant was removed, about 40ml lysis buffer was added, and a handheld ultrasonic disruptor was used to thoroughly break down the cell membrane and organelle membrane, thereby fully releasing the protein. The cell lysis process needed to be carried out on ice. After working for 15 s, the ultrasonic disruptor needed to pause for a while to prevent the lysate from overheating. Subsequently, the 50ml centrifuge tube containing cell lysate was centrifuged using a high-speed centrifuge (10000*g*, 10 min). The supernatant was taken and placed in a new 50 ml centrifuge tube. Added 200μl of anti-HA antibody and 200 μl A/G beads and placed it on a 4°C rotor to invert overnight. The next day, the 50ml centrifuge tube was taken out, and at this point, the beads in the centrifuge tube could be seen to be distributed in a flocculent manner. The 50ml centrifuge tube was centrifuged (3000*g*, 1 min, 4 °C) and the supernatant was aspirated off. The beads were then washed with 30 ml of fresh Triton lysis buffer for three times at 4°C. Then, the beads were mixed with an equal volume of 2x SDS sample buffer (bromophenol blue free), followed by subjected to SDS-PAGE. After staining with 5% (m/v) Coomassie Brilliant Blue (Sigma, cat. C.I.42660B0149) dissolved in 45% (v/v) methanol and 5% (v/v) acetic acid in water) for 30 min, the gels were decolorized (in 45% (v/v) methanol and 5% (v/v) acetic acid in water) overnight (during this period, it was necessary to replace the decolorizing solution with fresh one) and the excised gel segments were subjected to in-gel chymotrypsin digestion, and then dried.

Selected the gel segment between 85KDa and 125KDa which TRPV4 located. Samples were analyzed on a nanoElute (Bruker) coupled to a timsTOF Pro (Bruker) equipped with a CaptiveSpray source. Peptides were dissolved in 10 μl of 0.1% formic acid (v/v) and were loaded onto a homemade C18 column (35 cm × 75 μm, ID of 1.9 μm, 100 Å). Samples were then eluted with linear gradients of 3-35% acetonitrile (v/v, in 0.1% formic acid) at a flow rate of 0.3 μl/min, for 60 min. MS data were acquired with a timsTOF Pro MS (Bruker) operated in PASEF mode, and were analyzed using Peaks Studio Xpro software (PEAKS Studio 10.6 build 20201221, Bioinformatics Solutions). The human UniProt Reference Proteome database was used for data analysis, during which the parameters were set as: a) precursor and fragment mass tolerances: 20 ppm and 0.05 Da; b) semi-specific digest mode: allowed; c) maximal missed cleavages per peptide.

### Quantitative mass spectrometry

Stable over-expression TRPV4 cell lines were constructed in BNL-CL2 and AML12, and TRPV4 protein was enriched from lysed cells. The specific steps for protein enrichment and preparation of gel samples are elaborated in the **Mass spectrometry analyses** section. Samples were subjected to in-gel digestion, dried, and then analyzed using an Orbitrap Fusion Lumos mass spectrometer (Thermo SCIENTIFIC) equipped with an EASY-IC ion source. Peptides were eluted for 120 min with linear gradients of 3-35% acetonitrile in 0.1% formic acid at a flow rate of 300 nl/min. The data-dependent acquisition raw files were analyzed by Proteome Discoverer 2.5 software against Uniprot database (Mus musculus) to determine the m/z, z, and start/stop time of the TRPV4 peptides^69^. It should be noted that the over-expressed TRPV4 protein in cell lines was of homo sapiens origin. Due to the highly conserved amino acid sequence of TRPV4 in both homo sapiens and mus musculus, and the complete consistency of the amino acid sequence near the K608 site between humans and mice, using the Uniport database of mus musculus TRPV4 to match the mass spectrometry data of human TRPV4 protein was feasible. The two TRPV4 peptide sequences selected in this experiment were completely identical in both human and mouse TRPV4 proteins. Peptide 1, TGTYSIMIQK(+42.01): m/z was 592.299, z was 2, start time was 45.01 min, stop time was 65.01 min, RT was 55.01 min. Peptide 2, GVPNPIDLLESTLYESSVVPGPK: m/z was 1206.13902, z was 2, start time was 118.1866 min, stop time was 138.1866 min, RT was 128.1866 min. The above two peptides of TRPV4 protein were constructed and deemed as a reference for TRPV4 acetylation peptides in mouse liver tissues. The next step was to prepare tissue samples for Parallel Reaction Monitoring (PRM) mass spectrometry. The 8-week aged mice, D1 newborn, E18.5 foetal liver tissues were collected, homogenized in lysis buffer (2 μl lysis buffer / 3 mg tissues), and followed by sonication and centrifugation at 4 °C for 15 min. Then, extracted the supernatant and mixed with an equal volume of 2x SDS sample buffer (bromophenol blue free), followed by subjected to SDS-PAGE. After staining with 5% (m/v) Coomassie Brilliant Blue (Sigma, cat. C.I.42660B0149) dissolved in 45% (v/v) methanol and 5% (v/v) acetic acid in water) for 30 min, the gels were decolorized (in 45% (v/v) methanol and 5% (v/v) acetic acid in water) overnight (during this period, it was necessary to replace the decolorizing solution with fresh one) and the excised gel segments were subjected to in-gel chymotrypsin digestion, and then dried. Selected the gel segment between 85KDa and 125KDa which TRPV4 located. Further PRM parameters included an automatic gain control (AGC) of 1 × 10^5^, a maximum injection time of 1000 ms, and a precursor isolation window width of m/z 1. Skyline-daily 21.2.1.424 was used to analyze the PRM data^70^. And acetylated amino acid sites of TRPV4 identification in liver tissues were compared and matched with cell line TRPV4 acetylation peptides.

### Measurement of ATP, ADP and AMP

ATP, ADP and AMP from cells or liver tissues were analysed by CE-MS. Each measurement required cells collected from a 10-cm dish or 30-100 mg of liver tissues dissected by freeze clamp. The following steps were carried out as described in previous studies^12,69,71,72^:

a. After anaesthetising, the mouse was placed in a supine position, followed by wet the abdominal hair with 75% alcohol using a spray bottle;
b. An inverted T-shaped incision was made below the sternum xiphoid of the mouse with sharp scissors. Note that the transverse incision should be kept at approximately 1 cm long, and the longitudinal incision approximately 0.5 cm long;
c. The upper limbs of the mouse were pressed down with the thumbs of both hands, and the back to the ventral side was then pushed by the index fingers until the liver was prolapsed from the incision, while the mouse’s body was supported by the remaining fingers. During this process, any compressed or congested on the liver should be avoided;
d. Once one lobe of the liver is fully exposed, a clamp that has been pre-cooled in liquid nitrogen is quickly removed from the liquid nitrogen bucket by another assistant. The mouse liver is then clamped to rapidly freeze it into thin sheets of tissue until it becomes brittle;
e. Approximately 50-100 mg of liver tissues were rapidly collected using liquid nitrogen-cooled forceps. The tissues were then placed into a 1.5-ml tube and stored in liquid nitrogen for future use.
f. The tissue was grounded in 1 ml of methanol with IS1 after pre-cooled by liquid nitrogen. The lysate was then mixed with 1 ml of chloroform and 400 μl of water by 20 s of vortexing. After centrifugation at 15,000*g* for 15 min at 4 °C, 450 μl of aqueous phase was collected and was then filtrated through a 5-kDa cutoff filter (cat. OD003C34, PALL) by centrifuging at 12,000*g* for 3 h at 4 °C.
g. For measuring the adenylates of cells, the cells cultured in 10-cm dish were rinsed with 20 ml of 5% (m/v) mannitol solution (dissolved in water) and instantly frozen in liquid nitrogen. Cells were then lysed with 1 ml of methanol containing IS1 (50 µM L-methionine sulfone, 50 µM D-campher-10-sulfonic acid, dissolved in water; 1:500 (v/v) added to the methanol and used to standardise the metabolite intensity and to adjust the migration time), and were scraped from the dish.
h. In parallel, quality control samples were prepared by combining 10 μl of the aqueous phase from each sample and then filtered alongside the samples.
i. The filtered aqueous phase was then freeze-dried in a vacuum concentrator at 4 °C, and then dissolved in 100 μl of water containing IS2 (50 µM 3-aminopyrrolidine dihydrochloride, 50 µM N-diethyl-2-phenylacetamide, 50 µM trimesic acid, 50 µM 2-naphtol-3,6-disulfonic acid disodium salt, dissolved in methanol; used to adjust the migration time).
j. A total of 20 μl of re-dissolved solution was then loaded into an injection vial (cat. 9301-0978, Agilent; equipped with a snap cap (cat. 5042-6491, Agilent)). Before CE-MS analysis, the fused-silica capillary (cat. TSP050375, i.d. 50 µm × 80 cm; Polymicro Technologies) was installed in a CE/MS cassette (cat. G1603A, Agilent) on the CE system (Agilent Technologies 7100).
k. The capillary was then pre-conditioned with Conditioning Buffer (25 mM ammonium acetate, 75 mM diammonium hydrogen phosphate, pH 8.5) for 30 min, followed by balanced with Running Buffer (50 mM ammonium acetate, pH 8.5; freshly prepared) for another 1 h.
l. CE-MS analysis was run at anion mode, during which the capillary was washed by Conditioning Buffer, followed by injection of the samples at a pressure of 50 mbar for 25 s, and then separation with a constant voltage at −30 kV for another 40 min. Sheath Liquid (0.1 μM hexakis(1H, 1H, 3H-tetrafluoropropoxy)phosphazine, 10 μM ammonium trifluoroacetate, dissolved in methanol/water (50% v/v); freshly prepared) was flowed at 1 ml/min through a 1:100 flow splitter (Agilent Technologies 1260 Infinity II; actual flow rate to the MS: 10 μl/min) throughout each run.
m. The parameters of MS (Agilent Technologies 6545) were set as: a) ion source: Dual AJS ESI; b) polarity: negative; c) nozzle voltage: 2,000 V; d) fragmentor voltage: 110 V; e) skimmer voltage: 50 V; f) OCT RFV: 500 V; g) drying gas (N_2_) flow rate: 7 L/min; h) drying gas (N_2_) temperature: 300 °C; i) nebulizer gas pressure: 8 psig; j) sheath gas temperature: 125 °C; k) sheath gas (N_2_) flow rate: 4 L/min; l) capillary voltage (applied onto the sprayer): 3,500 V; m) reference (lock) masses: m/z 1,033.988109 for hexakis(1H, 1H, 3H-tetrafluoropropoxy)phosphazine, and m/z 112.985587 for trifluoroacetic acid; n) scanning range: 50-1,100 m/z; and o) scanning rate: 1.5 spectra/s. Data were collected using MassHunter LC/MS acquisition 10.1.48 (Agilent Technologies), and were processed using Qualitative Analysis B.06.00 (Agilent Technologies).
n. Levels of AMP, ADP and ATP were measured using full scan mode with m/z values of 346.0558, 426.0221 and 505.9885. Note that a portion of ADP and ATP could lose one phosphate group during in-source fragmentation, thus leaving the same m/z ratios as AMP and ADP, and should be corrected according to their different retention times in the capillary. Therefore, the total amount of ADP is the sum of the latter peak of the m/z 346.0558 spectrogramme and the former peak of the m/z 426.0221 spectrogramme, and the same is applied for ATP. Note that the retention time of each metabolite may vary between each run, and can be adjusted by isotope-labelled standards (dissolved in individual cell or tissue lysates) run between each sample, so do IS1 and IS2.

### Measurement of intracellular amino acids

The normal and being fasted 24 h pregnant E18.5 C57BL/6 female mice were anesthetized with isoflurane, and the 50 mg foetal livers were used to detect the intracellular amino acids by HPLC-MS (SCIEX, QTRAP 5500) descripted^61,66^ as in the following:

a. The foetal livers were grounding and lysed in 1 mL of ice-cold methanol. The lysates were then mixed with 1 mL of chloroform and 400 µL of water (containing 4 µg/mL [U-13C]-glutamine as an IS), followed by 20 s of vortexing.
b. After centrifugation at 15,000× g for another 15 min at 4 °C, 600 µL of the aqueous phase was collected, lyophilized in a vacuum concentrator at 4 °C for 8 h, and then dissolved in 30 µL of 50% (v/v, in water) acetonitrile, followed by loading 20 µL of solution into an injection vial (Cat# 5182-0714, Agilent; with an insert (Cat# HM-1270, Zhejiang Hamag Technology)) equipped with a snap cap (Cat# HM-2076, Zhejiang Hamag Technology).
c. A total of 2 µl of each sample was loaded onto a HILIC column (ZIC-pHILIC, 5 μm, 2.1 × 100 mm, PN: 1.50462.0001, Millipore). The mobile phase consisted of 15 mM ammonium acetate containing 3 ml l–1 ammonium hydroxide (>28%, v/v) in the LC–MS-grade water (mobile phase A) and LC–MS-grade 90% (v/v) acetonitrile in LC–MS-grade water (mobile phase B) run at a flow rate of 0.2 ml min^−1^.
d. The amino acids were separated with the following HPLC gradient elution programme: 95% B held for 2 min, then to 45% B in 13 min, held for 3 min, and then back to 95% B for 4 min. The mass spectrometer was run on a Turbo V ion source in negative mode with a spray voltage of −4,500 V, source temperature at 550 °C, gas no.1 at 50 psi, gas no.2 at 55 psi, and curtain gas at 40 psi.
e. Metabolites were measured using the multiple reactions monitoring mode, and declustering potentials and collision energies were optimized using analytical standards. The Data were collected using Analyst 1.7.1 software (SCIEX), and the relative amounts of metabolites were analysed using MultiQuant 3.0.3 software (SCIEX). The protein wet weight of each sample was determined by Bradford assay after dissolving the naturally dried protein sediment with 0.2 M KOH at room temperature.

### Measurement of glycolytic intermediates

Changes of glycolytic intermediates in the hepatic cell lines, primary hepatocytes and foetal livers after glucose starvation.

a. Foetal liver tissues were prepared in the following: the normal and being fasted 24 h pregnant E18.5 C57BL/6 female mice were anesthetized with isoflurane, and the 50 mg foetal livers frozen in liquid nitrogen. Sample preparation method had been described in the **Measurement of intracellular amino acids** section.
b. The adult and foetal primary hepatocytes were isolated and cultured as in **the Primary hepatocytes** section. And the primary hepatocytes, cell line BNL-CL2 and AML12 were under glucose starvation for 2 h, 4h, 8 h and 24 h, then cells in 10 cm dish were rinsed with 20 ml PBS and instantly frozen in liquid nitrogen. Then removed the 10 cm cell dish from liquid nitrogen and placed it on ice for about 1 minute, added 1ml of ice-cold methanol and collect the cells using a cell scraper. The lysates were then mixed with 1 mL of chloroform and 400 µL of water (containing 4 µg/mL [U-13C]-glutamine as an IS), followed by 20 s of vortexing. The subsequent sample preparation steps were consistent with the tissue sample preparation.
c. Glycolytic intermediates were detected by HPLC-MS in the **Measurement of intracellular amino acids** section.

### RNA-seq and qRT-PCR

The mRNAs of BNL-CL2 cells cultured in high-glucose DMEM medium supplemented with 10% FBS, 100 IU penicillin, 100 mg/ml streptomycin were isolated using 1 ml of TRIzol™ LS Reagent (Thermo Fisher Scientific, Catalog. 10296010), and the samples were delivered on dry ice to the Shanghai Ouyi Biomedical Technology Co., Ltd for RNA-seq library construction. Fastp^73^ (v0.23.1) (https://github.com/OpenGene/fastp) was used to trim the Illumina next-generation sequencing adapters, and low-quality and short-length reads were filtered from the RNA-seq data. The RNA-seq reads were processed using the ENCODE RNA-seq-pipeline (https://github.com/ENCODE-DCC/rna-seq-pipeline). Briefly, cleaned fastq reads were aligned to the mm9 reference genome, and unique genome-wide signal coverage tracks were generated using samtools (1.9.0)^74^, bedGraphToBigWig, and bedSort UCSC tools (https://hgdownload.soe.ucsc.edu). Then, the Fragments Per Kilobase of exon model per Million mapped fragments (FPKM) value was calculated using the TopHat-Cufflinks pipeline based on the count number. The DESeq2 R package (v1.36.0)^75^ was used to identify differentially expressed genes with *p*-value < 0.05 and log_2_Fold-change > 1 as cut-offs.

Total RNA of 50 mg of E18.5d foetal liver was extracted with 900 µl of TRIzol™ LS Reagent and subjected to three rounds of freeze/thaw cycles before being homogenised. The homogenate was mixed with 200 µl of chloroform and then left at room temperature for 5 min, followed by centrifuge at 4 °C, 12,000*g* for another 15 min. The upper aqueous layer (approximately 450 µl) was carefully transferred to a new RNase-free tube. To precipitate the RNA, 450 µl of isopropanol was added, and then it was centrifuged at 12,000*g* for 30 min at 4 °C. The resulting pellet was washed twice with 75% ethanol and once with 100% ethanol before being dissolved in 20 µl of DEPC-treated water. The RNA concentration was measured using a NanoDrop 2000 spectrophotometer (Thermo). Approximately 1 µg of RNA was diluted with DEPC-treated water to a final volume of 10 µl, heated to 65 °C for 5 min, and then immediately placed on ice. Reverse transcription was performed using the RT Reagent Kit and gDNA Eraser (Takara, Japan), followed by incubation at 37 °C for 15 min, and then at 98 °C for 5 min in a thermocycler. The primers for *TRPV*1-6 descripted as the reference^10^ and the primers for internal reference *GAPDH* descripted as^76^. The reverse-transcribed cDNA was quantified using the Maxima SYBR Green/ROX qPCR Master Mix on a LightCycler 480 II System (Roche) with the following program: pre-denaturation at 95 °C for 10 min; denaturation at 95 °C for 10 s, followed by annealing and extension at 65 °C for 30 s for a total of 45 cycles. The parameters for annealing and extending were optimised based on amplification curves, melting curves, and bands from agarose gel electrophoresis, adjusting the annealing temperature for each primer pair accordingly.

### Immunofluorescence assays

Immunofluorescence-based imaging of LAMP2 and mTOR were performed as follows: (1) Prepared cell cultures: Placed sterile cover glass of an appropriate size in a 6-well plate, then added 2ml of culture medium in each well and inoculated cells. (2) Cell growth: Allowed the cells to grow to 50-80% confluence in a well of a 6-well dish before conducting subsequent glucose starvation or drug treatment experiments. (3) Post-treatment: After completing glucose starvation or drug treatment, aspirated the culture medium. (4) Fixation: Fixed the cultured cells in each well of 6-well plate with 2 ml of 4% paraformaldehyde for 15 min at room temperature. (5) Permeabilization: After washing with 2ml PBS three times, incubated cells with 2 ml of 0.05% Triton X-100 (Sigma, cat. P9416-100ML) for 15 min. (6) Blocking: Incubated the cells with PBS contained 5% bovine serum albumin. (7) Primary antibody incubation: Incubated the cells with primary antibody (LAMP2 (Abcam, cat. ab13524), mTOR (CST, cat.2983t) overnight at 4°C (protect from light). The antibody diluent is PBS containing 5% foetal bovine serum. The dilution ratios were LAMP2 (1:120, 120 µl diluent +1 µl antibody), mTOR (1:80, 80 µl diluent +1 µl antibody). (8) Secondary antibody incubation: After washing with PBS, incubated the cells with Alexa Fluor 488- or 594-conjugated secondary antibodies (Invitrogen, Carlsbad, CA, USA) for overnight at 4°C, protected from light. The antibody diluent was PBS containing 5% foetal bovine serum. The dilution ratio was 1:100 (100 µl diluent +1 µl antibody). (9) Imaging: Obtain tiled images using an inverted epifluorescence microscope (Zeiss LSM 780 or LSM 980) with a 63× 1.4 NA oil objective at regular intervals. Ensure the exposure time for each channel was kept constant for all slides on a given day.

Immunofluorescence-based imaging of LAMP2 and AXIN were performed similarly, with the following adjustments: (1) Cell growth: Allowed the cells to grow to 70-80% confluence before conducting subsequent glucose starvation or drug treatment experiments. (2) Primary antibody incubation: Used primary antibodies (LAMP2 (Abcam, cat. ab13524), AXIN (Santa Cruz, cat. sc-8567)) with dilution ratios of LAMP2 (1:120, 120 μl diluent + 1 μl antibody), AXIN1 (1:20-50, 20-50 μl diluent + 1 μl antibody). Pixel location was quantified using ZEN 3.4 (blue) as described in **Image analysis**.

### Indirectly reflecting v-ATPase activity by detecting lysosomal pH

Firstly, added 2ml of culture medium to the 35-mm glass bottom dish (Cellvis, cat: D35-20-0-N) or the 4-chamber glass bottom dish (Cellvis, cat: D35C4-20-0-N) and inoculated cells. When the cells have grown to 70-80% confluence in a well of a 35-mm glass bottom dish, subsequent glucose starvation or drug treatment experiments can be conducted. Next, prepared the fluorescent dye: (1) Calculated the required amount of culture medium based on the need for 2 ml of culture medium containing fluorescent dye required for each 35-mm glass bottom culture dish. (2) Added the corresponding amount of LysosensorTM Green DND-189 (Thermo L7535, 1mM) to the culture medium (working concentration is 1 μM).

Carefully removed the original culture medium from the 35-mm glass bottom dish, and then added 2ml of culture medium containing LysosensorTM Green DND-189 to each 35-mm glass bottom culture dish. Incubated the cells with 1 μM LysosensorTM Green DND-189 for 30 min.

Subsequently, detected the fluorescence intensity of the cells using an inverted epifluorescence microscope (Zeiss LSM 780 or LSM 980). The excitation wavelength was 488nM. Capturing images used a Zeiss LSM 780 or LSM980 with a 63× 1.4 NA oil objective at regular intervals. During imaging, maintained live cells at 37°C, 5% CO2 in a humidified incubation chamber (ZEISS, Incubator PM S1). Kept the laser intensity and voltage settings constant for slides from the same batch. Due to the variation in fluorescence intensity with prolonged incubation time, it was necessary to capture samples from the same batch as quickly as possible. The activity of v-ATPase, assessed by the fluorescent intensity of Lysosensor dye, positively correlated with the lysosomal acidity^11^. Signal intensity was quantified using ZEN 3.4 (blue) as described in **Image analysis**.

### TRPV4 activity measurement

BNL-CL2 expressing TRPV4-GCaMP6s-fused protein were directly imaged by an inverted epifluorescence microscope (Zeiss LSM 780 or LSM 980)^10,77^. The excitation wavelength was set to 488nM. During imaging, live cells were maintained at 37°C, 5% CO2 in a humidified incubation chamber (ZEISS, Incubator PM S1). Images were captured using a Zeiss LSM 780 with a 63× 1.4 NA oil objective at regular intervals. To minimize cells damage and fluorescence quenching caused by excessive laser, the laser intensity and voltage should be as low as possible, and the shooting should be performed quickly. Signal intensity was quantified using ZEN 3.4 (blue) as described in **Image analysis** section.

### Image analysis

Quantification of fluorescence intensity, pixel location, and hepatocyte size was performed using the imaging software ZEN 3.4 (blue) and Image J. The analysis was conducted on greyscale 1,048 × 1,048 pixel images acquired at 63× magnification of cells immunostained with the anti-mTOR antibody. These images were utilized for measuring cellular mTOR, AXIN localization.

### Statistical analysis

Statistical analyses were performed using Prism 9 (GraphPad software). The normality of each group of data was assessed using the following tests where applicable: Anderson-Darling test, Kolmogorov-Smirnov test, Anderson-Darling test, or Shapiro-Wilk test. The homogeneity of variance test used the F-test. When the data conforms to both homogeneities of variance and normal distribution simultaneously, an unpaired two-tailed Student’s t-test was used to determine the significance between the two groups. If the variances between the groups were unequal, Welch’s correction was applied. For data that did not follow a normal distribution or homogeneity of variance, an unpaired two-tailed Mann-Whitney test was used to determine the significance between the groups. The adjusted means and standard error of the mean (SEM) were calculated and recorded when the data met the above standards. Differences were considered statistically significant when the p-value was less than 0.05.

## Notes

### Competing Interest Statement

The authors have declared no competing interest.

