## Supplementary information for "Prenatal liver is resistant to low glucose-induced inhibition of mTORC1"

Chen-Jie Zhang (张辰颉）[1,2,3,4#], Chuan-Jin Yu （余传金）[3,5#], Yi Cheng （程旖）[6], Jie-Xue Pan （潘洁雪）[3,5], Long-Yun Ye （叶龙云）[7], Yun-Hui Tang （唐蕴慧）[3], Xue-Yun Qin （秦薛云）[3], Zhong-Liang Lin （林忠亮）[1], Chen-Song Zhang （张宸崧）[8****], Guo-Lian Ding （丁国莲）[3,5***], Sheng-Cai Lin （林圣彩）[8**], He-Feng Huang （黄荷凤）[1,2,3,4,5*]

[1]Department of Obstetrics and Gynaecology, Center for Reproductive Medicine, the Fourth Affiliated Hospital of School of Medicine, and International School of Medicine, International Institutes of Medicine, Zhejiang University, Yiwu, China.

[2]Key Laboratory of Reproductive Genetics (Ministry of Education), Department of Reproductive Endocrinology, Women's Hospital, Zhejiang University School of Medicine, Hangzhou, China.

[3]Obstetrics and Gynaecology Hospital, Institute of Reproduction and Development, Fudan University, Shanghai, China.

[4] The International Peace Maternity and Child Health Hospital, School of Medicine, Shanghai Jiao Tong University, Shanghai, China.

[5] Shanghai Key Laboratory of Reproduction and Development, Shanghai, China.

[6]The First Affiliated Hospital, Zhejiang University School of Medicine, Hangzhou, China.

[7]Department of Pancreatic Surgery, Fudan University Shanghai Cancer Center, Shanghai, China.

[8]Metabolic Research Center, State-Province Joint Engineering Research Center of Targeted Drugs from Natural Products, School of Life Sciences, Xiamen University, Fujian, China.

[#]These authors contributed equally: Chen-Jie Zhang, Chuan-Jin Yu.

**This file includes:**

Full sequence of TRPV4-WT, TRPV4-K608R, GCaMP6s (Pages 3to 9)

The details of constructing gene knock-in mouse models (Pages 10 to 26)

Rosa26-LSL-TRPV4-WT-HA Genotyping Protocol (Pages 27 to 29)

Rosa26-LSL-TRPV4-K608R-HA Genotyping Protocol (Pages 30 to 32)

Supplementary Fig. 1 (gel source data, Pages 33 to 58)

Full sequence: TRPV4-WT Homo sapiens CCDS9134.1

ATGGCGGATTCCAGCGAAGGCCCCCGCGCGGGGCCCGGGGAGGTGGCTGA

GCTCCCCGGGGATGAGAGTGGCACCCCAGGCGGGGAGGCTTTTCCTCTCT

CCTCCCTGGCCAATCTGTTTGAGGGGGAGGATGGCTCCCTTTCGCCCTCAC

CGGCTGATGCCAGTCGCCCTGCTGGCCCAGGCGATGGGCGACCAAATCTG

CGCATGAAGTTCCAGGGCGCCTTCCGCAAGGGGGTGCCCAACCCCATCGA

TCTGCTGGAGTCCACCCTATATGAGTCCTCGGTGGTGCCTGGGCCCAAGAA

AGCACCCATGGACTCACTGTTTGACTACGGCACCTATCGTCACCACTCCAG

TGACAACAAGAGGTGGAGGAAGAAGATCATAGAGAAGCAGCCACAGAGC

CCCAAAGCCCCTGCCCCTCAGCCGCCCCCCATCCTCAAAGTCTTCAACCGG

CCTATCCTCTTTGACATCGTGTCCCGGGGCTCCACTGCTGACCTGGACGGG

CTGCTCCCATTCTTGCTGACCCACAAGAAACGCCTAACTGATGAGGAGTTT

CGAGAGCCATCTACGGGGAAGACCTGCCTGCCCAAGGCCTTGCTGAACCT

GAGCAATGGCCGCAACGACACCATCCCTGTGCTGCTGGACATCGCGGAGC

GCACCGGCAACATGCGGGAGTTCATTAACTCGCCCTTCCGTGACATCTACTA

TCGAGGTCAGACAGCCCTGCACATCGCCATTGAGCGTCGCTGCAAACACTA

CGTGGAACTTCTCGTGGCCCAGGGAGCTGATGTCCACGCCCAGGCCCGTG

GGCGCTTCTTCCAGCCCAAGGATGAGGGGGGCTACTTCTACTTTGGGGAGC

TGCCCCTGTCGCTGGCTGCCTGCACCAACCAGCCCCACATTGTCAACTACC

TGACGGAGAACCCCCACAAGAAGGCGGACATGCGGCGCCAGGACTCGCG

AGGCAACACAGTGCTGCATGCGCTGGTGGCCATTGCTGACAACACCCGTG

AGAACACCAAGTTTGTTACCAAGATGTACGACCTGCTGCTGCTCAAGTGTG

CCCGCCTCTTCCCCGACAGCAACCTGGAGGCCGTGCTCAACAACGACGGC

CTCTCGCCCCTCATGATGGCTGCCAAGACGGGCAAGATTGGGATCTTTCAG

CACATCATCCGGCGGGAGGTGACGGATGAGGACACACGGCACCTGTCCCG

CAAGTTCAAGGACTGGGCCTATGGGCCAGTGTATTCCTCGCTTTATGACCTC

TCCTCCCTGGACACGTGTGGGGAAGAGGCCTCCGTGCTGGAGATCCTGGT

GTACAACAGCAAGATTGAGAACCGCCACGAGATGCTGGCTGTGGAGCCCA

TCAATGAACTGCTGCGGGACAAGTGGCGCAAGTTCGGGGCCGTCTCCTTCT

ACATCAACGTGGTCTCCTACCTGTGTGCCATGGTCATCTTCACTCTCACCGC

CTACTACCAGCCGCTGGAGGGCACACCGCCGTACCCTTACCGCACCACGGT

GGACTACCTGCGGCTGGCTGGCGAGGTCATTACGCTCTTCACTGGGGTCCT

GTTCTTCTTCACCAACATCAAAGACTTGTTCATGAAGAAATGCCCTGGAGT

GAATTCTCTCTTCATTGATGGCTCCTTCCAGCTGCTCTACTTCATCTACTCTG

TCCTGGTGATCGTCTCAGCAGCCCTCTACCTGGCAGGGATCGAGGCCTACC

TGGCCGTGATGGTCTTTGCCCTGGTCCTGGGCTGGATGAATGCCCTTTACTT

CACCCGTGGGCTGAAGCTGACGGGGACCTATAGCATCATGATCCAGAAGAT

TCTCTTCAAGGACCTTTTCCGATTCCTGCTCGTCTACTTGCTCTTCATGATCG

GCTACGCTTCAGCCCTGGTCTCCCTCCTGAACCCGTGTGCCAACATGAAGG

TGTGCAATGAGGACCAGACCAACTGCACAGTGCCCACTTACCCCTCGTGCC

GTGACAGCGAGACCTTCAGCACCTTCCTCCTGGACCTGTTTAAGCTGACCA

TCGGCATGGGCGACCTGGAGATGCTGAGCAGCACCAAGTACCCCGTGGTC

TTCATCATCCTGCTGGTGACCTACATCATCCTCACCTTTGTGCTGCTCCTCA

ACATGCTCATTGCCCTCATGGGCGAGACAGTGGGCCAGGTCTCCAAGGAG

AGCAAGCACATCTGGAAGCTGCAGTGGGCCACCACCATCCTGGACATTGA

GCGCTCCTTCCCCGTATTCCTGAGGAAGGCCTTCCGCTCTGGGGAGATGGT

CACCGTGGGCAAGAGCTCGGACGGCACTCCTGACCGCAGGTGGTGCTTCA

GGGTGGATGAGGTGAACTGGTCTCACTGGAACCAGAACTTGGGCATCATC

AACGAGGACCCGGGCAAGAATGAGACCTACCAGTATTATGGCTTCTCGCAT

ACCGTGGGCCGCCTCCGCAGGGATCGCTGGTCCTCGGTGGTACCCCGCGTG

GTGGAACTGAACAAGAACTCGAACCCGGACGAGGTGGTGGTGCCTCTGG

ACAGCATGGGGAACCCCCGCTGCGATGGCCACCAGCAGGGTTACCCCCGC

AAGTGGAGGACTGATGACGCCCCGCTCTAG

Full sequence: TRPV4-K608R Homo sapiens

ATGGCGGATTCCAGCGAAGGCCCCCGCGCGGGGCCCGGGGAGGTGGCTGA

GCTCCCCGGGGATGAGAGTGGCACCCCAGGCGGGGAGGCTTTTCCTCTCT

CCTCCCTGGCCAATCTGTTTGAGGGGGAGGATGGCTCCCTTTCGCCCTCAC

CGGCTGATGCCAGTCGCCCTGCTGGCCCAGGCGATGGGCGACCAAATCTG

CGCATGAAGTTCCAGGGCGCCTTCCGCAAGGGGGTGCCCAACCCCATCGA

TCTGCTGGAGTCCACCCTATATGAGTCCTCGGTGGTGCCTGGGCCCAAGAA

AGCACCCATGGACTCACTGTTTGACTACGGCACCTATCGTCACCACTCCAG

TGACAACAAGAGGTGGAGGAAGAAGATCATAGAGAAGCAGCCACAGAGC

CCCAAAGCCCCTGCCCCTCAGCCGCCCCCCATCCTCAAAGTCTTCAACCGG

CCTATCCTCTTTGACATCGTGTCCCGGGGCTCCACTGCTGACCTGGACGGG

CTGCTCCCATTCTTGCTGACCCACAAGAAACGCCTAACTGATGAGGAGTTT

CGAGAGCCATCTACGGGGAAGACCTGCCTGCCCAAGGCCTTGCTGAACCT

GAGCAATGGCCGCAACGACACCATCCCTGTGCTGCTGGACATCGCGGAGC

GCACCGGCAACATGCGGGAGTTCATTAACTCGCCCTTCCGTGACATCTACTA

TCGAGGTCAGACAGCCCTGCACATCGCCATTGAGCGTCGCTGCAAACACTA

CGTGGAACTTCTCGTGGCCCAGGGAGCTGATGTCCACGCCCAGGCCCGTG

GGCGCTTCTTCCAGCCCAAGGATGAGGGGGGCTACTTCTACTTTGGGGAGC

TGCCCCTGTCGCTGGCTGCCTGCACCAACCAGCCCCACATTGTCAACTACC

TGACGGAGAACCCCCACAAGAAGGCGGACATGCGGCGCCAGGACTCGCG

AGGCAACACAGTGCTGCATGCGCTGGTGGCCATTGCTGACAACACCCGTG

AGAACACCAAGTTTGTTACCAAGATGTACGACCTGCTGCTGCTCAAGTGTG

CCCGCCTCTTCCCCGACAGCAACCTGGAGGCCGTGCTCAACAACGACGGC

CTCTCGCCCCTCATGATGGCTGCCAAGACGGGCAAGATTGGGATCTTTCAG

CACATCATCCGGCGGGAGGTGACGGATGAGGACACACGGCACCTGTCCCG

CAAGTTCAAGGACTGGGCCTATGGGCCAGTGTATTCCTCGCTTTATGACCTC

TCCTCCCTGGACACGTGTGGGGAAGAGGCCTCCGTGCTGGAGATCCTGGT

GTACAACAGCAAGATTGAGAACCGCCACGAGATGCTGGCTGTGGAGCCCA

TCAATGAACTGCTGCGGGACAAGTGGCGCAAGTTCGGGGCCGTCTCCTTCT

ACATCAACGTGGTCTCCTACCTGTGTGCCATGGTCATCTTCACTCTCACCGC

CTACTACCAGCCGCTGGAGGGCACACCGCCGTACCCTTACCGCACCACGGT

GGACTACCTGCGGCTGGCTGGCGAGGTCATTACGCTCTTCACTGGGGTCCT

GTTCTTCTTCACCAACATCAAAGACTTGTTCATGAAGAAATGCCCTGGAGT

GAATTCTCTCTTCATTGATGGCTCCTTCCAGCTGCTCTACTTCATCTACTCTG

TCCTGGTGATCGTCTCAGCAGCCCTCTACCTGGCAGGGATCGAGGCCTACC

TGGCCGTGATGGTCTTTGCCCTGGTCCTGGGCTGGATGAATGCCCTTTACTT

CACCCGTGGGCTGAAGCTGACGGGGACCTATAGCATCATGATCCAGAGGAT

TCTCTTCAAGGACCTTTTCCGATTCCTGCTCGTCTACTTGCTCTTCATGATCG

GCTACGCTTCAGCCCTGGTCTCCCTCCTGAACCCGTGTGCCAACATGAAGG

TGTGCAATGAGGACCAGACCAACTGCACAGTGCCCACTTACCCCTCGTGCC

GTGACAGCGAGACCTTCAGCACCTTCCTCCTGGACCTGTTTAAGCTGACCA

TCGGCATGGGCGACCTGGAGATGCTGAGCAGCACCAAGTACCCCGTGGTC

TTCATCATCCTGCTGGTGACCTACATCATCCTCACCTTTGTGCTGCTCCTCA

ACATGCTCATTGCCCTCATGGGCGAGACAGTGGGCCAGGTCTCCAAGGAG

AGCAAGCACATCTGGAAGCTGCAGTGGGCCACCACCATCCTGGACATTGA

GCGCTCCTTCCCCGTATTCCTGAGGAAGGCCTTCCGCTCTGGGGAGATGGT

CACCGTGGGCAAGAGCTCGGACGGCACTCCTGACCGCAGGTGGTGCTTCA

GGGTGGATGAGGTGAACTGGTCTCACTGGAACCAGAACTTGGGCATCATC

AACGAGGACCCGGGCAAGAATGAGACCTACCAGTATTATGGCTTCTCGCAT

ACCGTGGGCCGCCTCCGCAGGGATCGCTGGTCCTCGGTGGTACCCCGCGTG

GTGGAACTGAACAAGAACTCGAACCCGGACGAGGTGGTGGTGCCTCTGG

ACAGCATGGGGAACCCCCGCTGCGATGGCCACCAGCAGGGTTACCCCCGC

AAGTGGAGGACTGATGACGCCCCGCTC

Full sequence: GCaMP6s

ATGGGTTCTCATCATCATCATCATCATGGTATGGCTAGCATGACTGGTGGAC

AGCAAATGGGTCGGGATCTGTACGACGATGACGATAAGGATCTCGCCACCA

TGGTCGACTCATCACGTCGTAAGTGGAATAAGACAGGTCACGCAGTCAGA

GCTATAGGTCGGCTGAGCTCACTCGAGAACGTCTATATCAAGGCCGACAAG

CAGAAGAACGGCATCAAGGCGAACTTCCACATCCGCCACAACATCGAGGA

CGGCGGCGTGCAGCTCGCCTACCACTACCAGCAGAACACCCCCATCGGCG

ACGGCCCCGTGCTGCTGCCCGACAACCACTACCTGAGCGTGCAGTCCAAA

CTTTCGAAAGACCCCAACGAGAAGCGCGATCACATGGTCCTGCTGGAGTT

CGTGACCGCCGCCGGGATCACTCTCGGCATGGACGAGCTGTACAAGGGCG

GTACCGGAGGGAGCATGGTGAGCAAGGGCGAGGAGCTGTTCACCGGGGT

GGTGCCCATCCTGGTCGAGCTGGACGGCGACGTAAACGGCCACAAGTTCA

GCGTGTCCGGCGAGGGTGAGGGCGATGCCACCTACGGCAAGCTGACCCTG

AAGTTCATCTGCACCACCGGCAAGCTGCCCGTGCCCTGGCCCACCCTCGTG

ACCACCCTGACCTACGGCGTGCAGTGCTTCAGCCGCTACCCCGACCACATG

AAGCAGCACGACTTCTTCAAGTCCGCCATGCCCGAAGGCTACATCCAGGA

GCGCACCATCTTCTTCAAGGACGACGGCAACTACAAGACCCGCGCCGAGG

TGAAGTTCGAGGGCGACACCCTGGTGAACCGCATCGAGCTGAAGGGCATC

GACTTCAAGGAGGACGGCAACATCCTGGGGCACAAGCTGGAGTACAACCT

GCCGGACCAACTGACTGAAGAGCAGATCGCAGAATTTAAAGAGGCTTTCT

CCCTATTTGACAAGGACGGGGATGGGACAATAACAACCAAGGAGCTGGGG

ACGGTGATGCGGTCTCTGGGGCAGAACCCCACAGAAGCAGAGCTGCAGG

ACATGATCAATGAAGTAGATGCCGACGGTGACGGCACAATCGACTTCCCTG

AGTTCCTGACAATGATGGCAAGAAAAATGAAATACAGGGACACGGAAGAA

GAAATTAGAGAAGCGTTCGGTGTGTTTGATAAGGATGGCAATGGCTACATC

AGTGCAGCAGAGCTTCGCCACGTGATGACAAACCTTGGAGAGAAGTTAAC

AGATGAAGAGGTTGATGAAATGATCAGGGAAGCAGACATCGATGGGGATG

GTCAGGTAAACTACGAAGAGTTTGTACAAATGATGACAGCGAAGTGA

**SMOC-Report for Rosa26 knock-in mouse model (TRPV4-WT)**

Contents


1. Background and Objective

  1.1 Objective

  1.2 Background

2. Abstract

3. Recombinant strategy

  3.1 Strategy figure：

  3.2 Sequence of sgRNAs

4. Acquisition and genotypic identification of mice

  4.1 F0 mice genotyping

  4.2 F1 mice genotyping

    4.2.1 Genotyping strategy of F1 mice

    4.2.2 PCR genotyping of homologous recombinant F1 mice

    4.2.3 Method for identifying 5'homologous recombinant F1 mice

    4.2.4 Method for identifying 3'homologous recombinant F1 mice

    4.2.5 Information of homologous recombinant F1 mice

### 1. Background and Objective

#### 1.1 Objective

Generation of the CAG-LSL-TRPV4-WT-HA-WPRE-pA cassette knock-in mouse model at Rosa26 gene locus via CRISPR/Cas9 technology.

#### 1.2 Background

Gene Name (MGI Number): Gt(ROSA)26Sor (104735)

Gene URL Link (MGI) : http://www.informatics.jax.org/marker/MGI:104735

Gene URL Link (Ensembl) :

http://asia.ensembl.org/Mus_musculus/Gene/Summary?g=ENSMUSG00000086429;r=6:113067428-113077333

Knockin Cassette: CAG-LSL-TRPV4-WT-HA-WPRE-pA

### 2. Abstract

The project process was as follows: 1) Cas9 mRNA and gRNA were produced by in vitro transcription; 2) donor vectors were constructed by in-fusion. The plasmid structure contains 5'homologous arm (3.3 kb), Knock-in, 3'homologous arm (3.3 kb) ; 3) the mixture of Cas9 mRNA,gRNA and donor vector was micro-injected into fertilized eggs (C57BL/6J), then 1 F0 mouse that identified by PCR and sequencing was generated; 4) F0 positive mouse was crossed with wild-type C57BL/6J mice to generate 6 F1 mice.

### 3. Recombinant strategy

#### 3.1 Strategy figure:

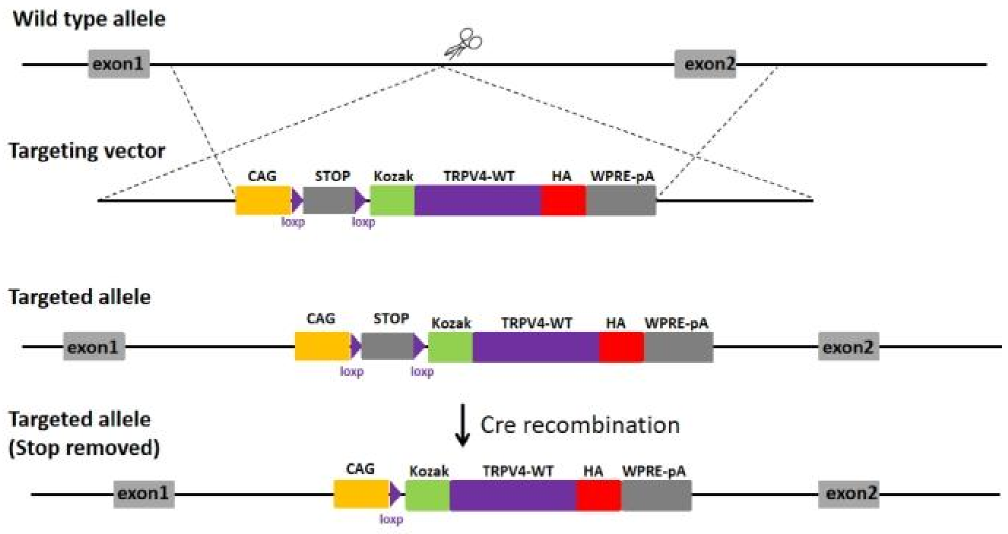


Strategy figure

#### 3.2 Sequence of sgRNAs

| gRNAs | Sequence (5'-3') |
| --- | --- |
| gRNA1 | GGGGACACACTAAGGGAGCT TGG |

### 4. Acquisition and genotypic identification of mice

#### 4.1 F0 mice genotyping

The injected fertilized eggs were transplanted into pseudo-pregnant mice, and the mice born about 20 days were F0 generation mice. The genotype were identified by PCR amplification and sequencing.Because the early cleavage rate of fertilized eggs is very fast, the F0 generation mice obtained are chimera and do not necessarily have the ability of stable inheritance. It is necessary to pass the positive F0 generation mice to obtain stable heritable F1 generation mice.

#### 4.2 F1 mice genotyping

F0 generation positive mice were mated with wild type C57BL/6J mice to obtain F1 generation mice. The genotype of F1 generation mice were identified by PCR method and sequencing.

##### 4.2.1 Genotyping strategy of F1 mice

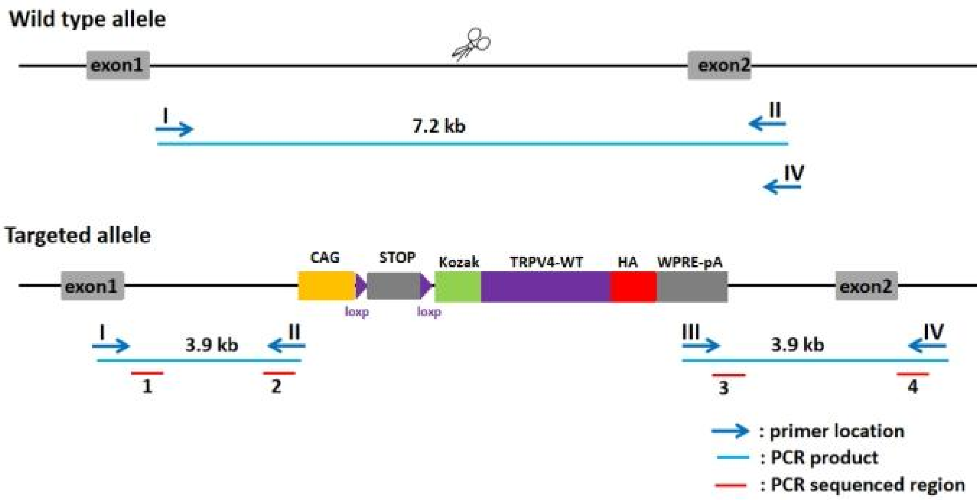


Strategy of F1 mice genotyping

Genotyping method:

5'homologous arm: 3.9 kb fragment should be amplified in the homologous recombinant DNA, and 7.2 kb fragment can be amplified in the negative DNA.

3'homologous arm: 3.9 kb fragment should be amplified in the homologous recombinant DNA, and none of fragments can be amplified in the negative DNA.

**4.2.2 PCR genotyping of homologous recombinant F1 mice**

The number of homologous recombinant F1 mice were 6,7,8,12,13,16. The agarose gel electrophoresis of PCR results were shown in Figure 3. All positive PCR products were confirmed by sequencing.

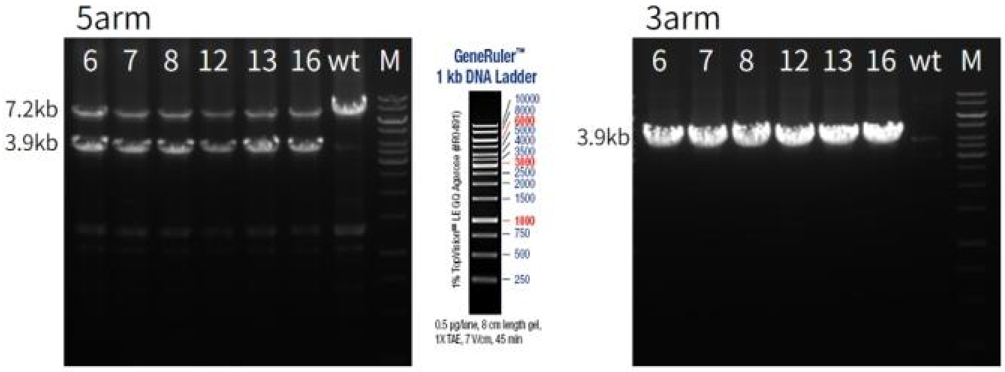


Agarose gel electrophoresis of PCR results.

（number：the number of F1 mice；M：1kb DNA ladder）

**4.2.3 Method for identifying 5’homologous recombinant F1 mice**

Primers：

| Primer | Sequence 5' --> 3' | Primer Type |
|---|---|---|
| I | TTCTGTGAGACAGCCGGGTA | Forward |
| II | TTTTTGGGGGTGATGGTGGTC | Reverse |

Reaction system:

| Reaction Component | Volume (μl) |
|---|---|
| ddH2O | 8.05 |

| | |
|---|---|
| 2xPCR Buffer | 10 |
| Primer I (20pmol/μl) | 0.3 |
| Primer II (20pmol/μl) | 0.3 |
| KOD-Multi&Epi-* | 0.35 |
| Genomic DNA | 1 |
| Total | 20 |

* KOD-Multi&Epi-（TOYOBO，Code No：KME-101）

PCR program：

| Step # | Temp (°C) | Time | Note |
|---|---|---|---|
| 1 | 94 | 3 min | - |
| 2 | 98 | 20 sec | - |
| 3 | 63 | 20 sec | - |
| 4 | 68 | 4 min | repeat steps 2-4 for 35cycles |
| 5 | 68 | 5 min | - |
| 6 | 12 | - | hold |

**4.2.4 Method for identifying 3'homologous recombinant F1 mice**

Primers：

| Primer | Sequence 5' --> 3' | Primer Type |
|---|---|---|
| III | TTGCCAGCCATCTGTTGTT | Forward |
| IV | TGCCACCTTTCACTTAGTTTGT | Reverse |

Reaction system:

| Reaction Component | Volume (μl) |
|---|---|
| ddH2O | 8.05 |
| 2xPCR Buffer | 10 |
| Primer III (20pmol/μl) | 0.3 |
| Primer IV (20pmol/μl) | 0.3 |
| KOD-Multi&Epi-* | 0.35 |
| Genomic DNA | 1 |
| Total | 20 |

* KOD-Multi&Epi-（TOYOBO，Code No：KME-101）

PCR program：

| Step # | Temp (°C) | Time | Note |
|---|---|---|---|
| 1 | 94 | 3 min | - |
| 2 | 98 | 20 sec | - |
| 3 | 63 | 20 sec | - |
| 4 | 68 | 4 min | repeat steps 2-4 for 35cycles |
| 5 | 68 | 5 min | - |
| 6 | 12 | - | hold |

#### 4.2.5 Information of homologous recombinant F1 mice

Table： The information of positive F1 mice

| Mice ID | DOB | Generations | Sex | Type | Genotype | Father | Mother |
|---|---|---|---|---|---|---|---|
| 6 | 2023/9/11 | F1 | ♂ | KI | He | 9F0 | 6J |
| 7 | 2023/9/11 | F1 | ♂ | KI | He | 9F0 | 6J |
| 8 | 2023/9/11 | F1 | ♂ | KI | He | 9F0 | 6J |
| 12 | 2023/9/11 | F1 | ♂ | KI | He | 9F0 | 6J |
| 13 | 2023/9/11 | F1 | ♀ | KI | He | 9F0 | 6J |
| 16 | 2023/9/11 | F1 | ♀ | KI | He | 9F0 | 6J |

**SMOC-Report for Rosa26 knock-in mouse model (TRPV4-K608R)**

Contents

5. Background and Objective

5.1 Objective

5.2 Background

6. Abstract

7. Recombinant strategy

7.1 Strategy figure：

7.2 Sequence of sgRNAs

8. Acquisition and genotypic identification of mice

8.1 F0 mice genotyping

8.2 F1 mice genotyping

8.2.1 Genotyping strategy of F1 mice

8.2.2 PCR genotyping of homologous recombinant F1 mice

8.2.3 Method for identifying 5’homologous recombinant F1 mice

8.2.4 Method for identifying 3’homologous recombinant F1 mice

8.2.5 Information of homologous recombinant F1 mice

### 5. Background and Objective

#### 5.1 Objective

Generation of the CAG-LSL-TRPV4-K608R-HA-WPRE-pA cassette knock-in mouse model at Rosa26 gene locus via CRISPR/Cas9 technology.

#### 5.2 Background

Gene Name (MGI Number): Gt(ROSA)26Sor (104735)

Gene URL Link (MGI) : http://www.informatics.jax.org/marker/MGI:104735

Gene URL Link (Ensembl) :

http://asia.ensembl.org/Mus_musculus/Gene/Summary?g=ENSMUSG00000086429;r=6:113067428-113077333

Knockin Cassette: CAG-LSL-TRPV4-K608R-HA-WPRE-pA

### 6. Abstract

The project process was as follows: 1) Cas9 mRNA and gRNA were produced by in vitro transcription; 2) donor vectors were constructed by in-fusion. The plasmid structure contains 5'homologous arm (3.3 kb), Knock-in, 3'homologous arm (3.3 kb) ; 3) the mixture of Cas9 mRNA,gRNA and donor vector was micro-injected into fertilized eggs (C57BL/6J), then 1 F0 mouse that identified by PCR and sequencing was generated; 4) F0 positive mouse was crossed with wild-type C57BL/6J mice to generate 9 F1 mice.

### 7. Recombinant strategy

#### 7.1 Strategy figure：

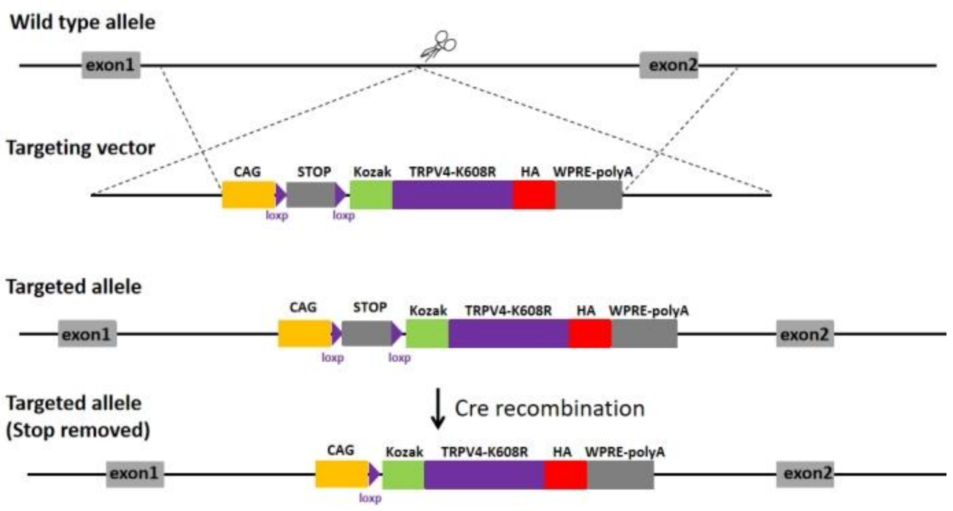


Strategy figure

#### 7.2 Sequence of sgRNAs

| gRNAs | Sequence (5’-3’) |
|---|---|
| gRNA1 | GGGGACACACTAAGGGAGCT TGG |

### 8. Acquisition and genotypic identification of mice

#### 8.1 F0 mice genotyping

The injected fertilized eggs were transplanted into pseudo-pregnant mice, and the mice born about 20 days were F0 generation mice. The genotype were identified by

PCR amplification and sequencing.Because the early cleavage rate of fertilized eggs is very fast, the F0 generation mice obtained are chimera and do not necessarily have the ability of stable inheritance. It is necessary to pass the positive F0 generation mice to obtain stable heritable F1 generation mice.

#### 8.2 F1 mice genotyping

F0 generation positive mice were mated with wild type C57BL/6J mice to obtain F1 generation mice. The genotype of F1 generation mice were identified by PCR method and sequencing.

##### 8.2.1 Genotyping strategy of F1 mice

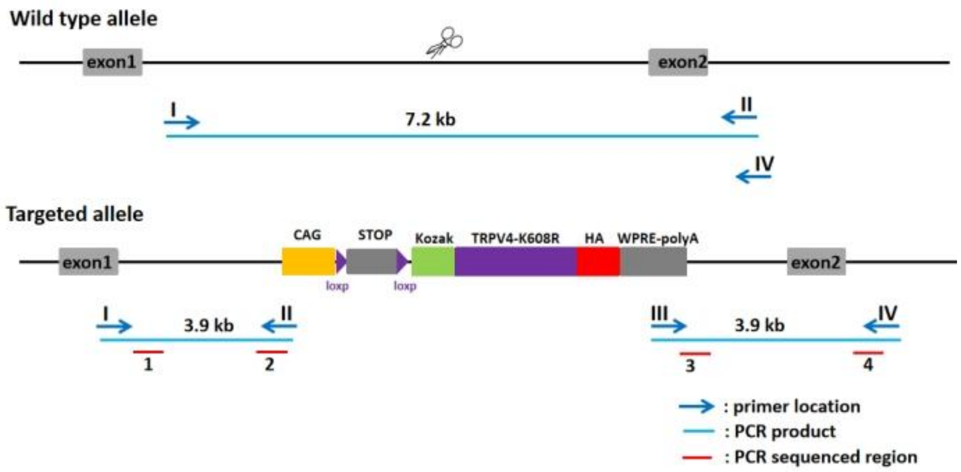


Strategy of F1 mice genotyping

Genotyping method:

5'homologous arm: 3.9 kb fragment should be amplified in the homologous recombinant DNA, and 7.2 kb fragment can be amplified in the negative DNA.

3’homologous arm: 3.9 kb fragment should be amplified in the homologous recombinant DNA, none of fragments can be amplified in the negative DNA.

#### 8.2.2 PCR genotyping of homologous recombinant F1 mice

The number of homologous recombinant F1 mice were 2,4,5,7,13,15,17,19,20. The agarose gel electrophoresis of PCR results were shown in Figure 3. All positive PCR products were confirmed by sequencing.

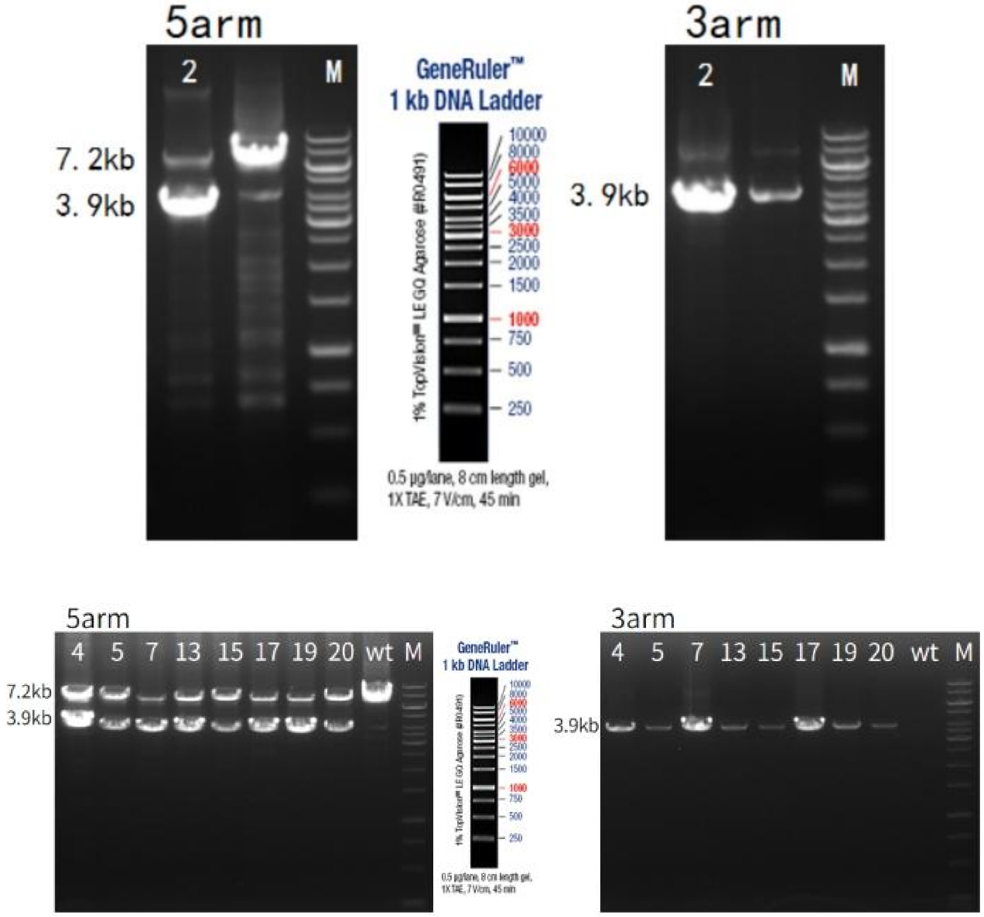


Agarose gel electrophoresis of PCR results.

（number：the number of F1 mice；M：1kb DNA ladder）

#### 8.2.3 Method for identifying 5’homologous recombinant F1 mice

Primers：

| Primer | Sequence 5' --> 3' | Primer Type |
|---|---|---|
| I | TTCTGTGAGACAGCCGGGTA | Forward |
| II | TTTTTGGGGGTGATGGTGGTC | Reverse |

Reaction system:

| Reaction Component | Volume (μl) |
|---|---|
| ddH2O | 8.05 |
| 2xPCR Buffer | 10 |
| Primer I (20pmol/μl) | 0.3 |
| Primer II (20pmol/μl) | 0.3 |
| KOD-Multi&Epi-* | 0.35 |
| Genomic DNA | 1 |
| Total | 20 |

* KOD-Multi&Epi-（TOYOBO，Code No：KME-101）

PCR program：

| Step # | Temp (℃) | Time | Note |
|---|---|---|---|
| 1 | 94 | 3 min | - |
| 2 | 98 | 20 sec | - |

| | | | |
|---|---|---|---|
| 3 | 63 | 20 sec | - |
| 4 | 68 | 4 min | repeat steps 2-4 for 35cycles |
| 5 | 68 | 5 min | - |
| 6 | 12 | - | hold |

#### 8.2.4 Method for identifying 3'homologous recombinant F1 mice

Primers：

| Primer | Sequence 5' --> 3' | Primer Type |
|---|---|---|
| III | TTGCCAGCCATCTGTTGTT | Forward |
| IV | TGCCACCTTTCACTTAGTTTGT | Reverse |

Reaction system:

| Reaction Component | Volume (µl) |
|---|---|
| ddH2O | 8.05 |
| 2xPCR Buffer | 10 |
| Primer III (20pmol/µl) | 0.3 |
| Primer IV (20pmol/µl) | 0.3 |
| KOD-Multi&Epi-* | 0.35 |
| Genomic DNA | 1 |

| | |
|---|---|
| Total | 20 |

* KOD-Multi&Epi-（TOYOBO，Code No：KME-101）

PCR program：

| Step # | Temp (℃) | Time | Note |
|---|---|---|---|
| 1 | 94 | 3 min | - |
| 2 | 98 | 20 sec | - |
| 3 | 63 | 20 sec | - |
| 4 | 68 | 4 min | repeat steps 2-4 for 35cycles |
| 5 | 68 | 5 min | - |
| 6 | 12 | - | hold |

#### 8.2.5 Information of homologous recombinant F1 mice

Table：The information of positive F1 mice

| Mice ID | DOB | Generations | Sex | Type | Genotype | Father | Mother |
|---|---|---|---|---|---|---|---|
| 2 | 2023/8/7 | F1 | ♀ | KI | He | 3F0 | 6J |
| 4 | 2023/8/14 | F1 | ♂ | KI | He | 3F0 | 6J |
| 5 | 2023/8/14 | F1 | ♂ | KI | He | 3F0 | 6J |
| 7 | 2023/8/14 | F1 | ♂ | KI | He | 3F0 | 6J |

| | | | | | | | |
|---|---|---|---|---|---|---|---|
| 13 | 2023/8/14 | F1 | ♂ | KI | He | 3F0 | 6J |
| 15 | 2023/8/14 | F1 | ♀ | KI | He | 3F0 | 6J |
| 17 | 2023/8/14 | F1 | ♀ | KI | He | 3F0 | 6J |
| 19 | 2023/8/14 | F1 | ♀ | KI | He | 3F0 | 6J |
| 20 | 2023/8/14 | F1 | ♀ | KI | He | 3F0 | 6J |

Rosa26-LSL-TRPV4-WT-HA Genotyping Protocol

| Common Name | Rosa26-LSL-TRPV4-WT-HA | Cat. NO. | |
|---|---|---|---|
| Strain of Origin | C57BL/6J | Version | V1 |

Genotyping strategy

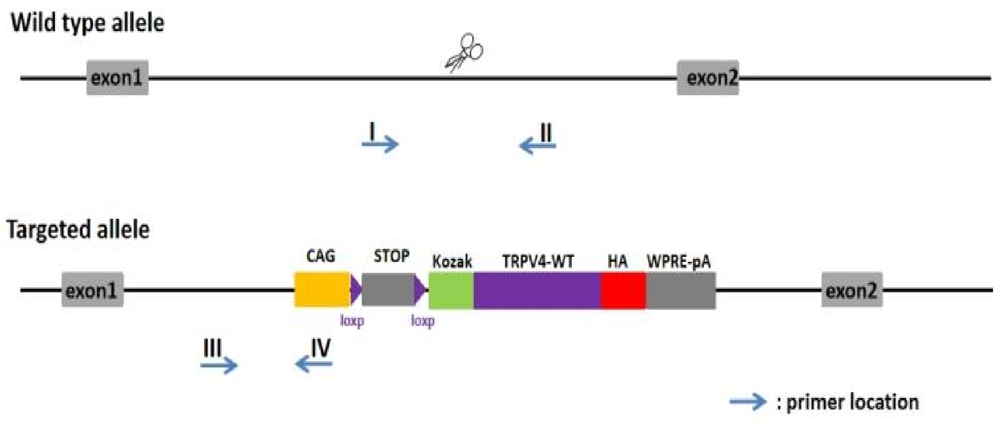


Primers

| Primer | Sequence (5'→3') | Primer type |
|---|---|---|
| P1 | TCAGATTCTTTTATAGGGGACACA | Forward |
| P2 | TAAAGGCCACTCAATGCTCACTAA | Reverse |
| P3 | GGGTGGGGTTGGGAAATCTT | Forward |
| P4 | TAGGGGGCGTACTTGGCATA | Reverse |

Expected results

| | |
|---|---|
| Results | 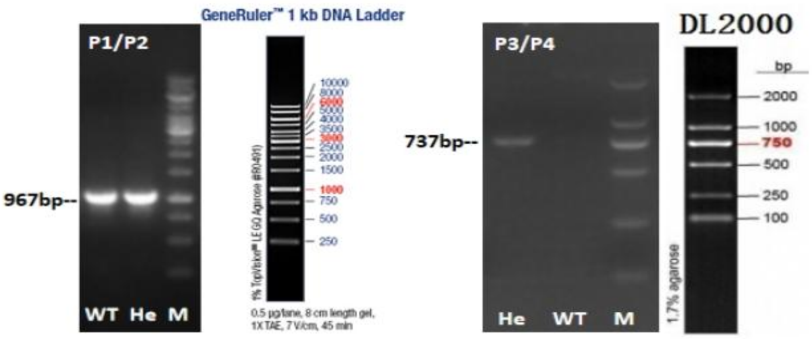 |
| Genotype | Wild type： P1P2 =967 bp；<br>Heterozygote： P1P2 =967 bp； P3P4=737 bp<br>Homozygote： P3P4 =737 bp |

Reaction &Cycling

| | | | | |
|---|---|---|---|---|
| PCR Reaction System | Reaction Component | | | Volume (μl) |
| | ddH2O | | | 8.0 |
| | 2×Taq Plus Master Mix | | | 10.0 |
| | P1(10 pmol/μl) or P3(10 pmol/μl) | | | 0.5 |
| | P2(10 pmol/μl) or P4(10 pmol/μl) | | | 0.5 |
| | Genomic DNA | | | 1.0 |
| | Total | | | 20 |
| | 2×Taq Plus Master Mix from Vazyme(Code Number: P222-1) | | | |
| Cycling | Step | Temp | Time | Note |

| | | | | |
|---|---|---|---|---|
| Reaction | 1 | 95°C | 5 min | |
| | 2 | 95°C | 30 sec | |
| | 3 | 60°C | 30 sec | |
| | 4 | 72°C | 30 sec | 35 repeats to 2 |
| | 5 | 72°C | 5 min | |
| | 6 | 12°C | Hold | |

Rosa26-LSL-TRPV4-K608R-HA Genotyping Protocol

| Common Name | Rosa26-LSL-TRPV4-K608R-HA | Cat. NO. | CM-KI-231843 |
|---|---|---|---|
| Strain of Origin | C57BL/6J | Version | V1 |

Genotyping strategy

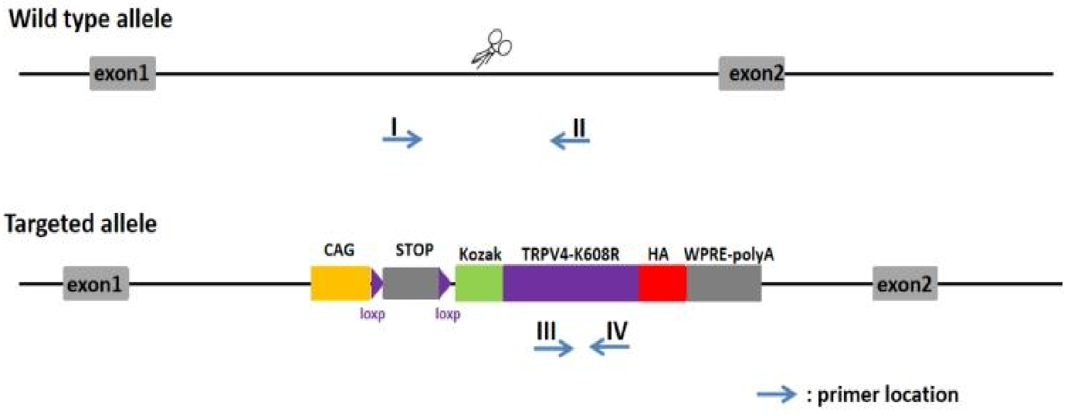


Primers

| Primer | Sequence (5’→3’) | Primer type |
|---|---|---|
| P1 | TCAGATTCTTTTATAGGGGACACA | Forward |
| P2 | TAAAGGCCACTCAATGCTCACTAA | Reverse |
| P3 | CCTGTGTGCCATGGTCATCT | Forward |
| P4 | CAGGAGGAAGGTGCTGAAGG | Reverse |

Expected results

| | |
|---|---|
| Results | 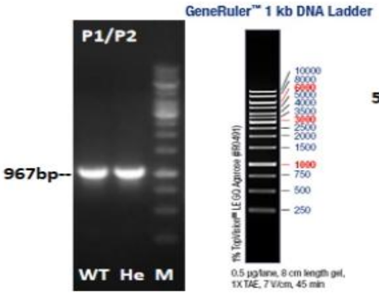  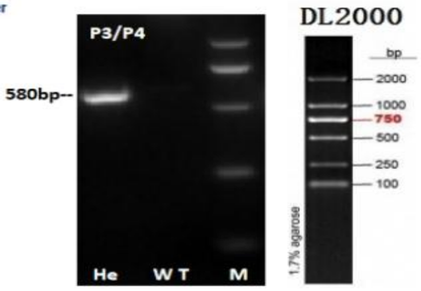  |
| Genotype | Wild type: P1P2 =967 bp;<br>Heterozygote: P1P2 =967 bp; P3P4=580 bp<br>Homozygote: P3P4 =580 bp |

Reaction &Cycling

| | | | | |
|---|---|---|---|---|
| PCR Reaction System | Reaction Component | | | Volume (µl) |
| | ddH2O | | | 8.0 |
| | 2×Taq Plus Master Mix | | | 10.0 |
| | P1(10 pmol/µl) or P3(10 pmol/µl) | | | 0.5 |
| | P2(10 pmol/µl) or P4(10 pmol/µl) | | | 0.5 |
| | Genomic DNA | | | 1.0 |
| | Total | | | 20 |
| | 2×Taq Plus Master Mix from Vazyme(Code Number: P222-1) | | | |
| Cycling Reaction | Step | Temp | Time | Note |
| | 1 | 95°C | 5 min | |

| | | | | |
|---|---|---|---|---|
| | 2 | 95°C | 30 sec | |
| | 3 | 60°C | 30 sec | |
| | 4 | 72°C | 30 sec | 35 repeats to 2 |
| | 5 | 72°C | 5 min | |
| | 6 | 12°C | Hold | |

**Supplementary Fig. 1 | Uncropped gels for Fig. 1 to 4 and Fig. S1, 8 to 10, 12.**

After electrophoretic transfer of proteins, the PVDF membranes were cut into strips containing sets of samples, and were then subjected to immunoblotting. Show here are films that had been exposed and developed to the membrane strips. The Pierce Prestained Protein MW Marker (Cat. 26612, ThermoFisher Scientific) was used as the protein markers.

Fig. 1A

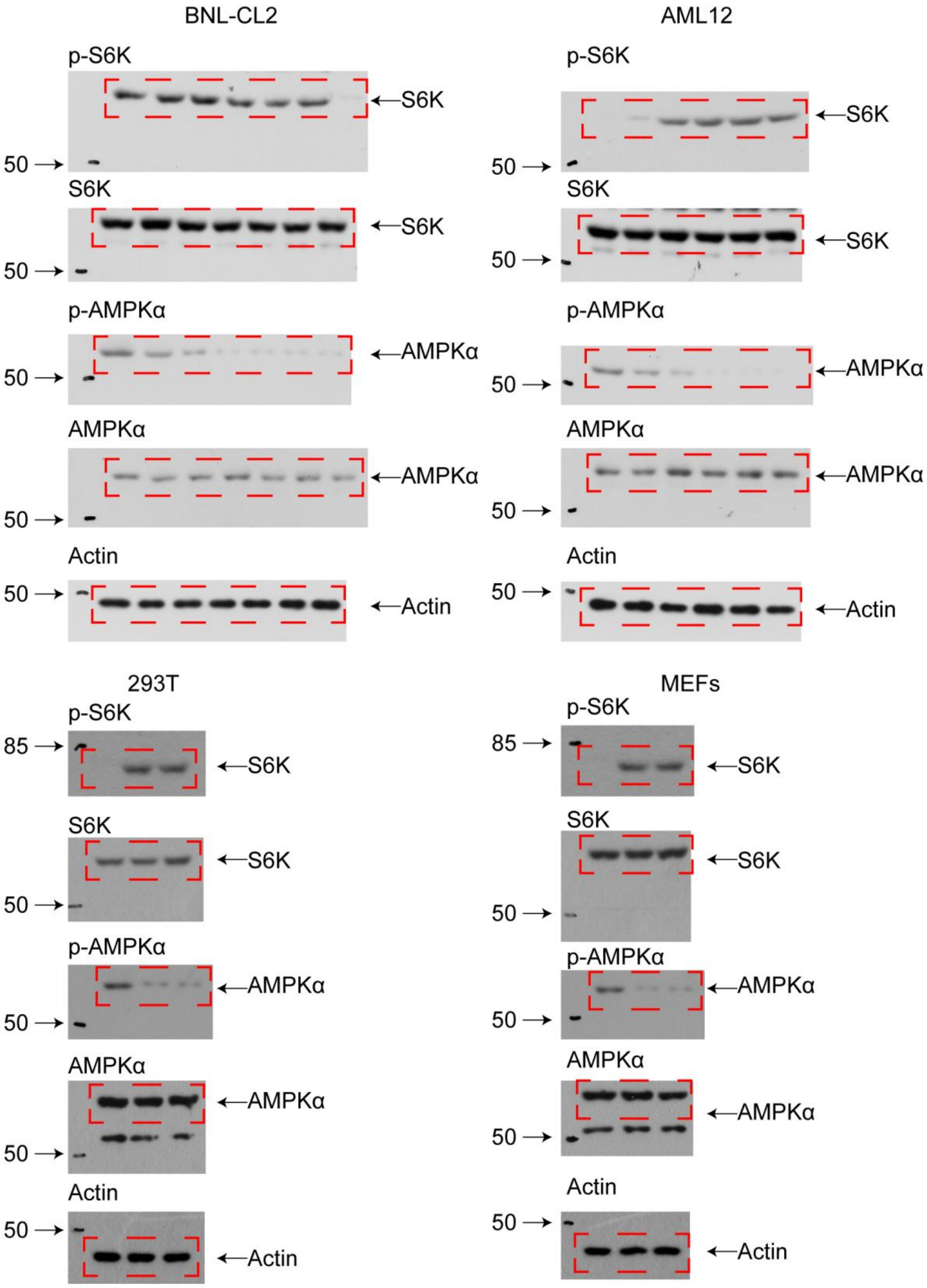

BNL-CL2
AML12
p-S6K
S6K
p-AMPKα
AMPKα
Actin
50
S6K
AMPKα
Actin
293T
MEFs
p-S6K
S6K
p-AMPKα
AMPKα
Actin
85
50
S6K
AMPKα
Actin

Fig. 1B

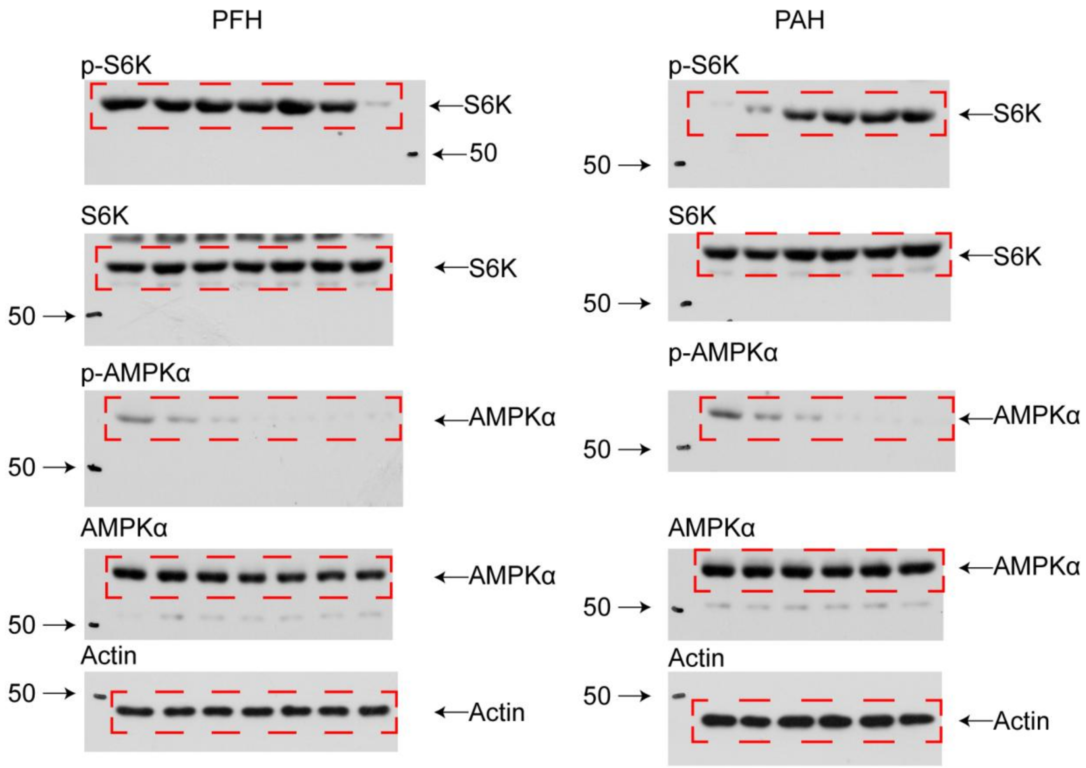

Fig. 1C

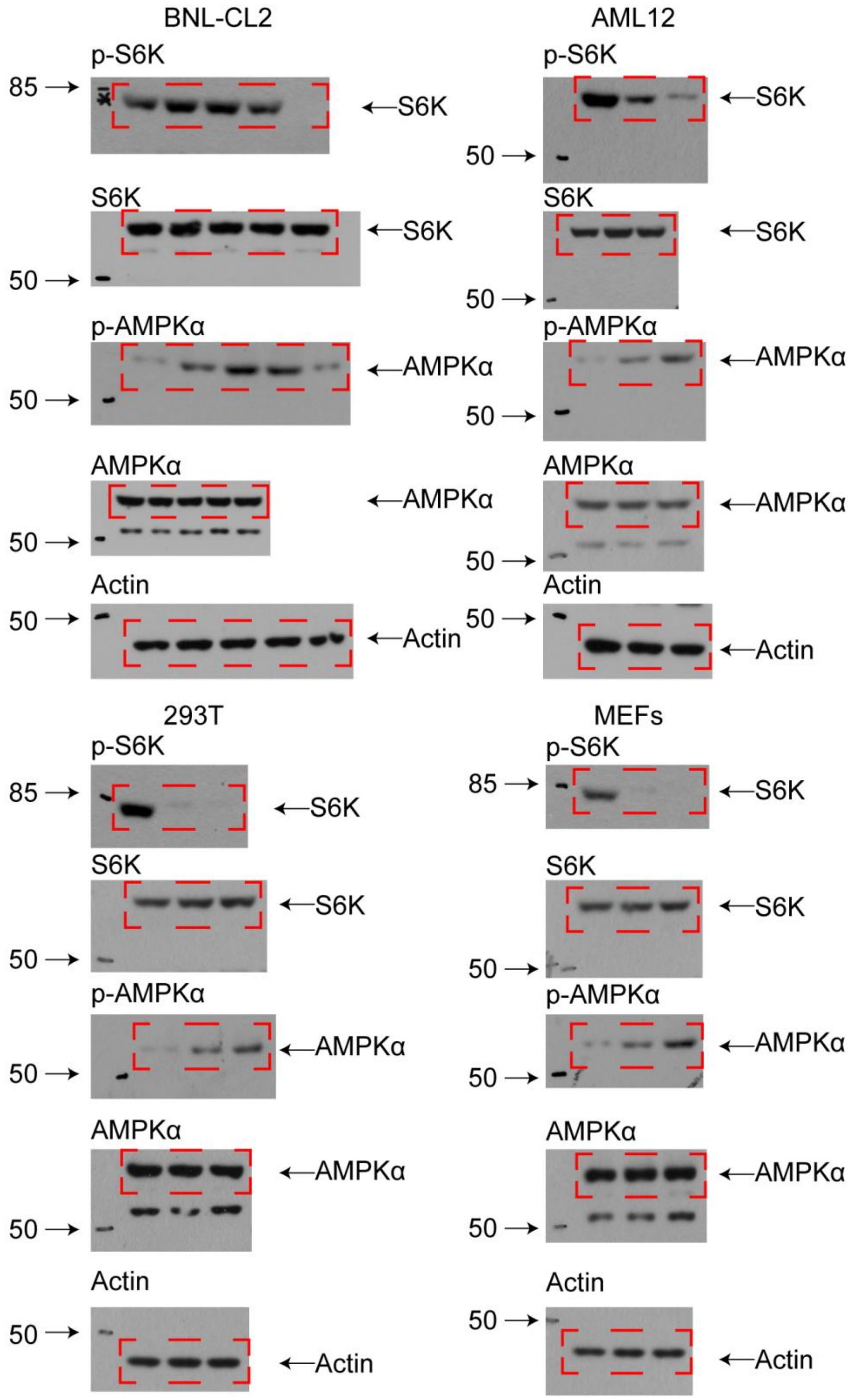

Fig. 1D

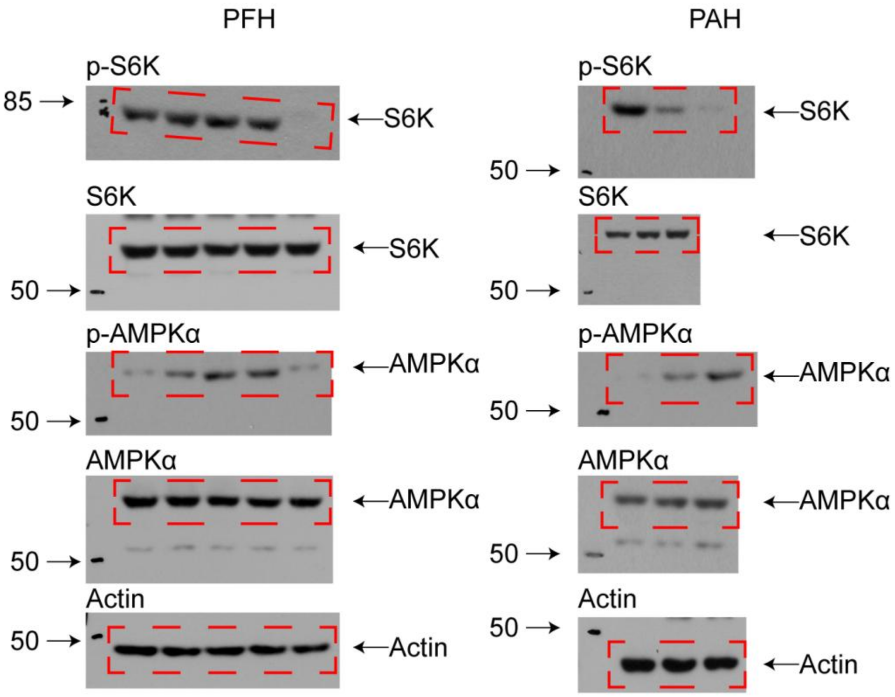
PFH
PAH
p-S6K
85
S6K
S6K
p-AMPKα
AMPKα
50
AMPKα
AMPKα
Actin
Actin
p-S6K
S6K
50
S6K
S6K
p-AMPKα
AMPKα
AMPKα
AMPKα
Actin
Actin

Fig. 1E

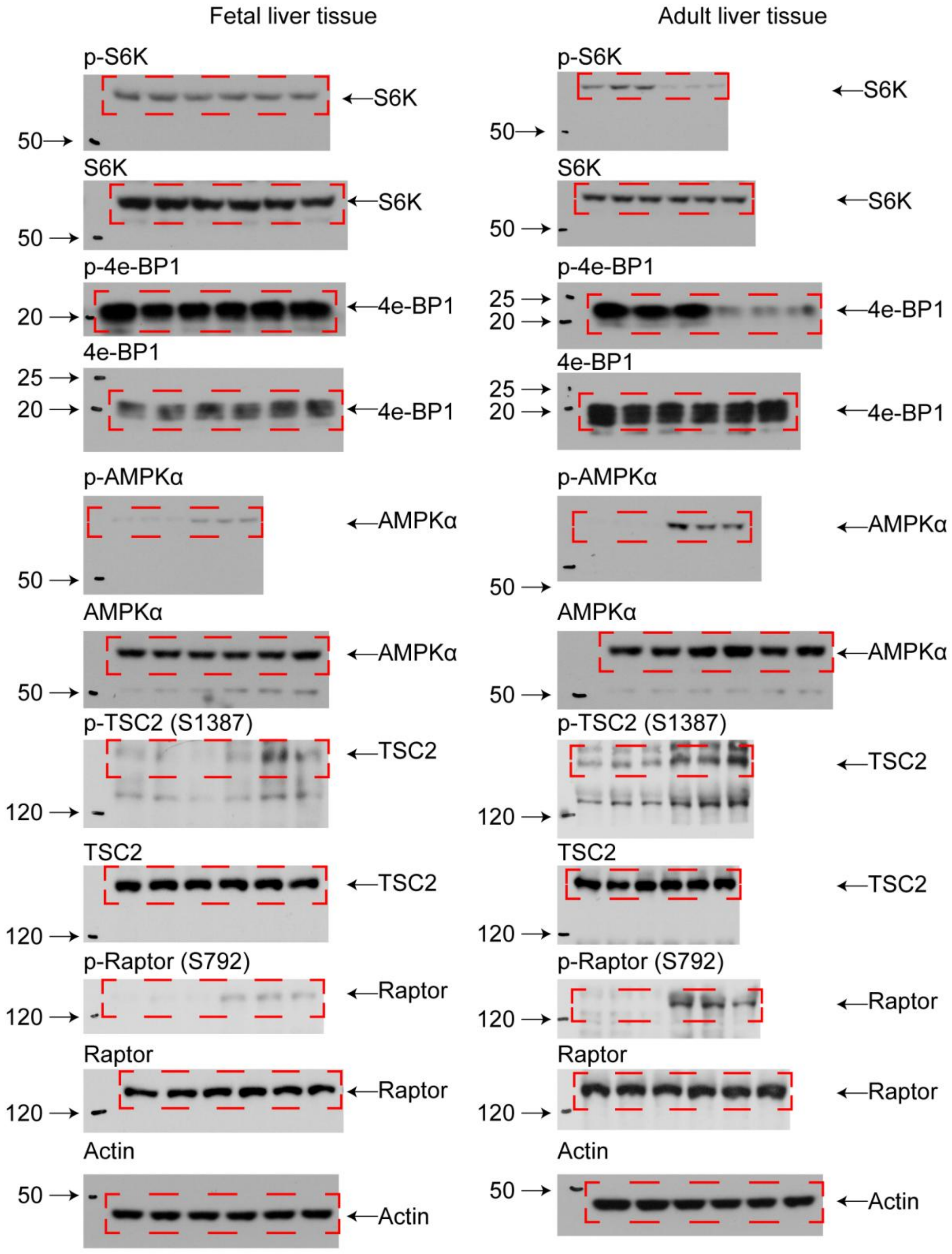
Fetal liver tissue
Adult liver tissue
p-S6K
S6K
50
S6K
p-4e-BP1
4e-BP1
25
20
p-AMPKα
AMPKα
p-TSC2 (S1387)
TSC2
120
p-Raptor (S792)
Raptor
Actin

Fig. 1F

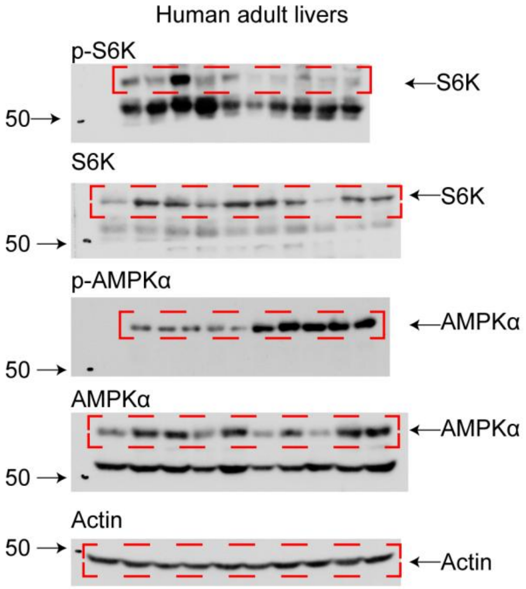
Human adult livers
p-S6K
50
S6K
S6K
50
S6K
p-AMPKα
AMPKα
50
AMPKα
AMPKα
50
Actin
50
Actin

Fig. 1I

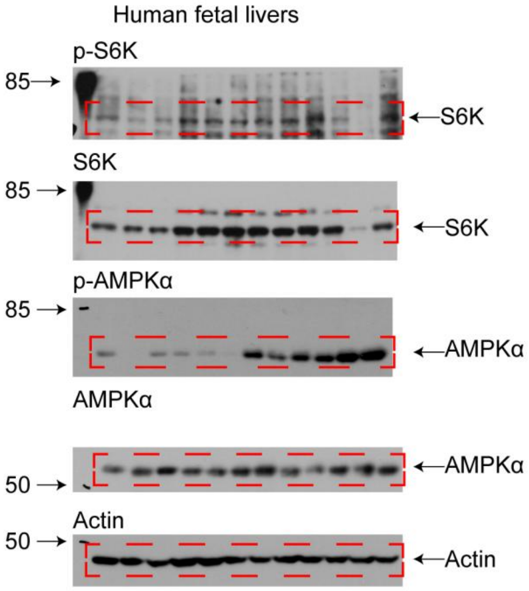
Human fetal livers
p-S6K
85
S6K
S6K
85
S6K
p-AMPKα
85
AMPKα
AMPKα
AMPKα
50
Actin
50
Actin

Fig. 1M

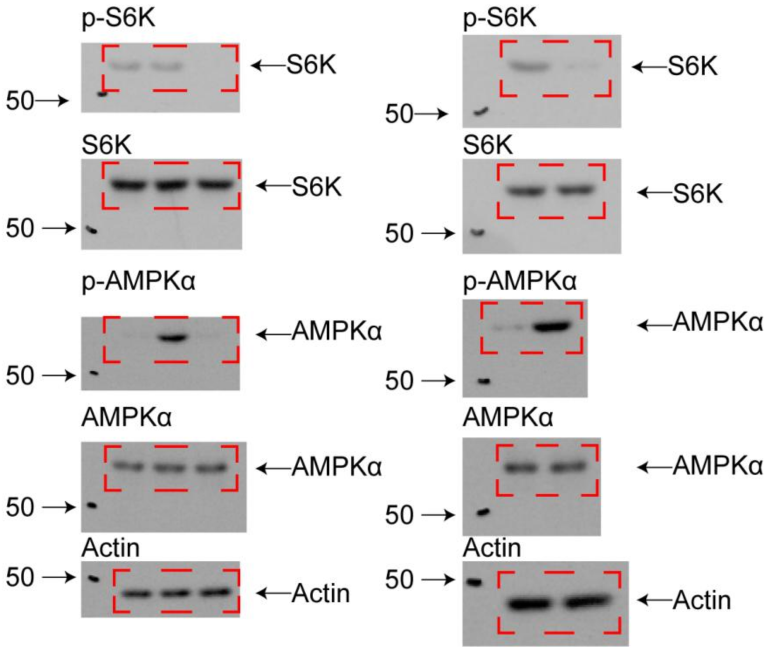

p-S6K
←S6K
50→
S6K
←S6K
50→
p-AMPKα
←AMPKα
50→
AMPKα
←AMPKα
50→
Actin
50→
←Actin
p-S6K
←S6K
50→
S6K
←S6K
50→
p-AMPKα
←AMPKα
50→
AMPKα
←AMPKα
50→
Actin
50→
←Actin

Fig. 2E

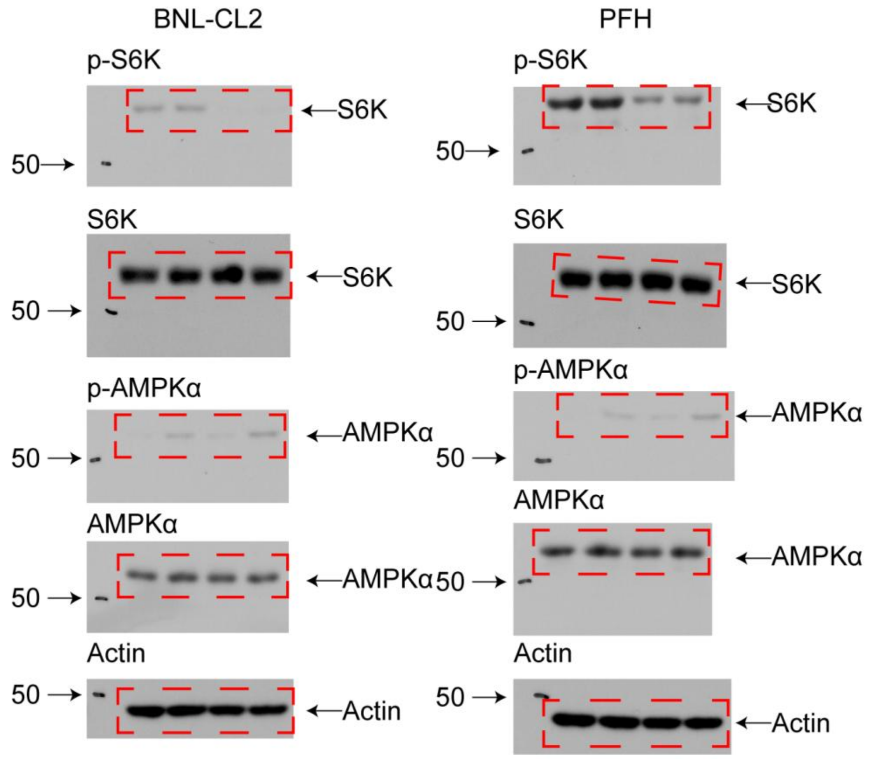

Fig. 2F

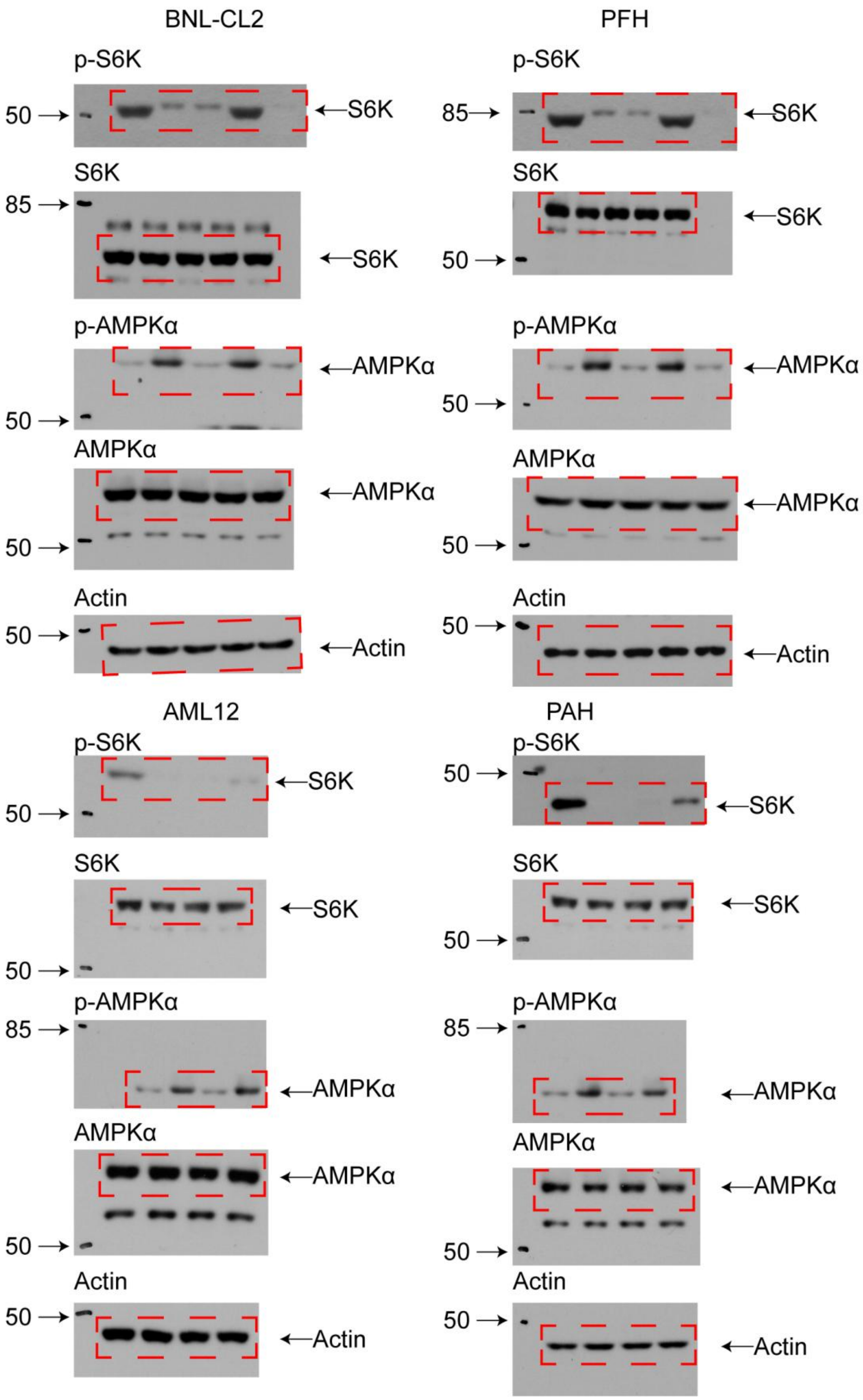
BNL-CL2
PFH
p-S6K
50
S6K
85
S6K
S6K
85
S6K
50
p-AMPKα
AMPKα
50
AMPKα
50
AMPKα
AMPKα
50
Actin
50
Actin
AML12
PAH
p-S6K
50
S6K
50
S6K
S6K
50
p-AMPKα
85
AMPKα
AMPKα
50
Actin
50
Actin

Fig. 2G

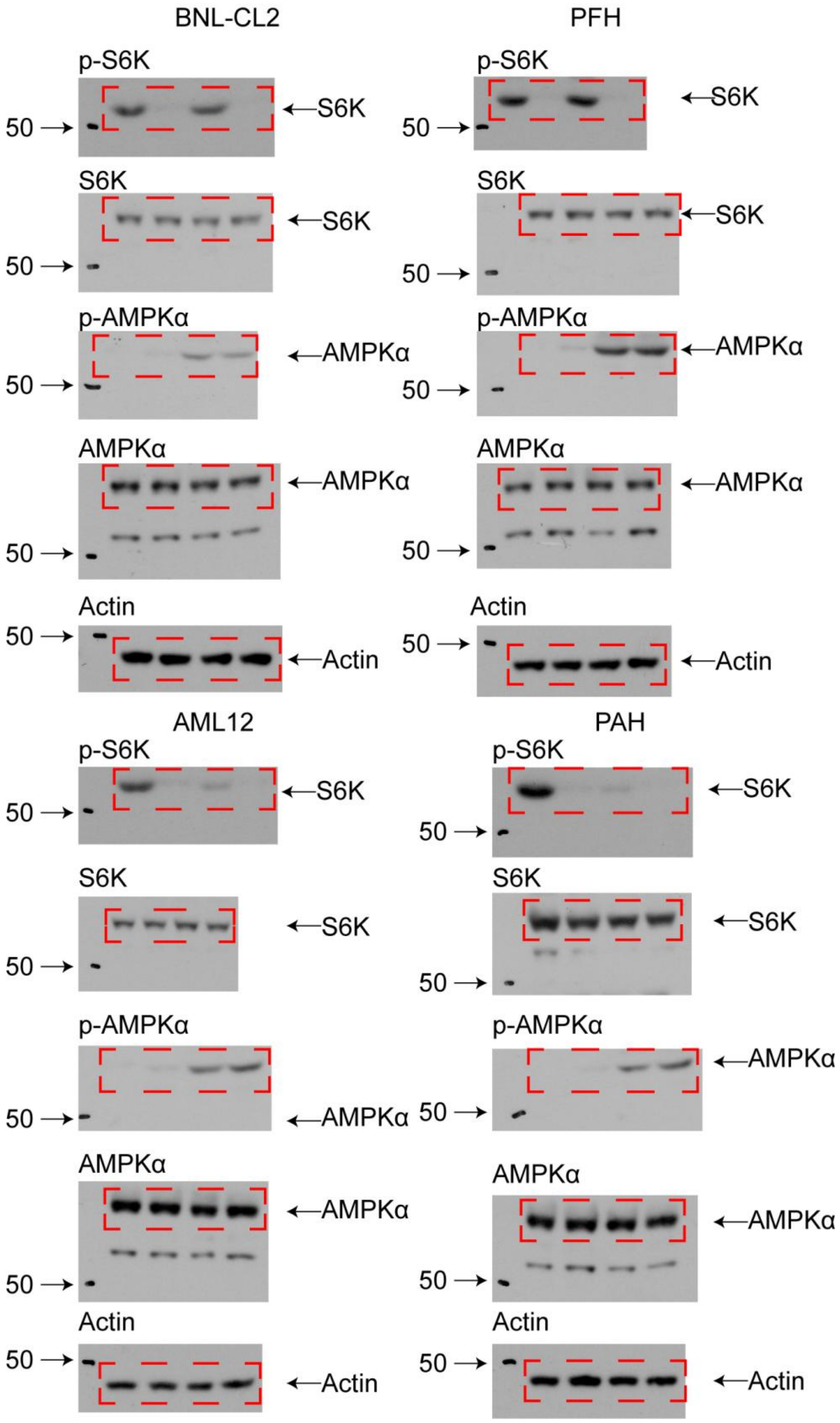

Fig. 2K

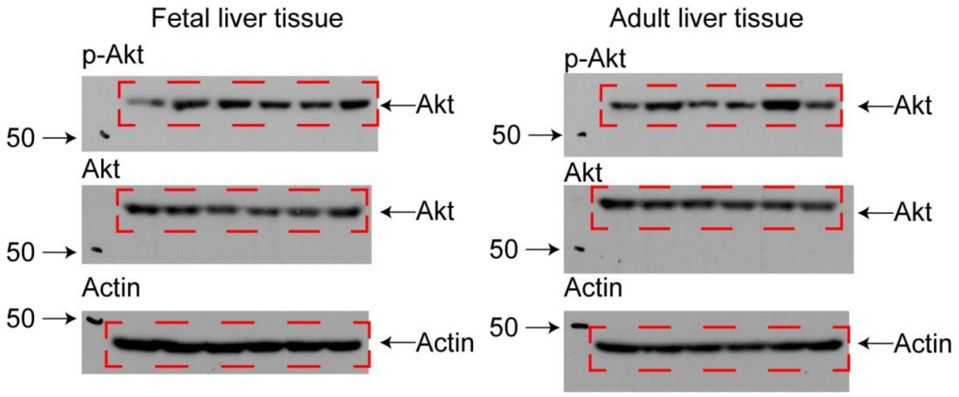

Fetal liver tissue
p-Akt
50
Akt
Akt
50
Akt
Actin
50
Actin
Adult liver tissue
p-Akt
50
Akt
Akt
50
Akt
Actin
50
Actin

Fig. 3A

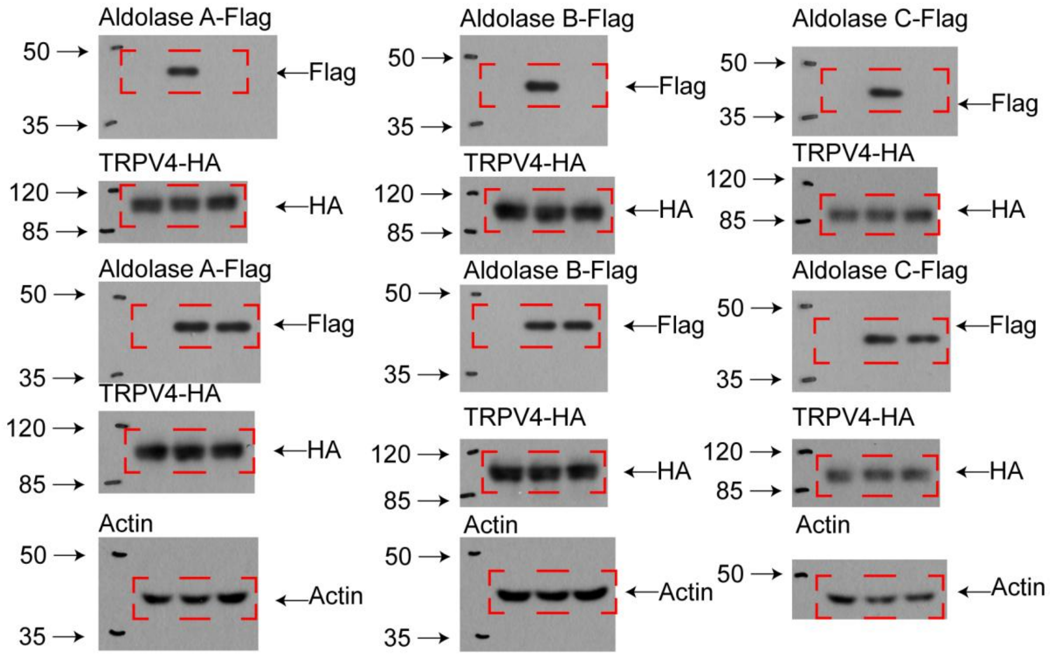
Aldolase A-Flag
50
35
Flag
TRPV4-HA
120
85
HA
Aldolase A-Flag
50
35
Flag
TRPV4-HA
120
85
HA
Actin
50
35
Actin
Aldolase B-Flag
50
35
Flag
TRPV4-HA
120
85
HA
Aldolase B-Flag
50
35
Flag
TRPV4-HA
120
85
HA
Actin
50
35
Actin
Aldolase C-Flag
50
35
Flag
TRPV4-HA
120
85
HA
Aldolase C-Flag
50
35
Flag
TRPV4-HA
120
85
HA
Actin
50
Actin

Fig. 3G

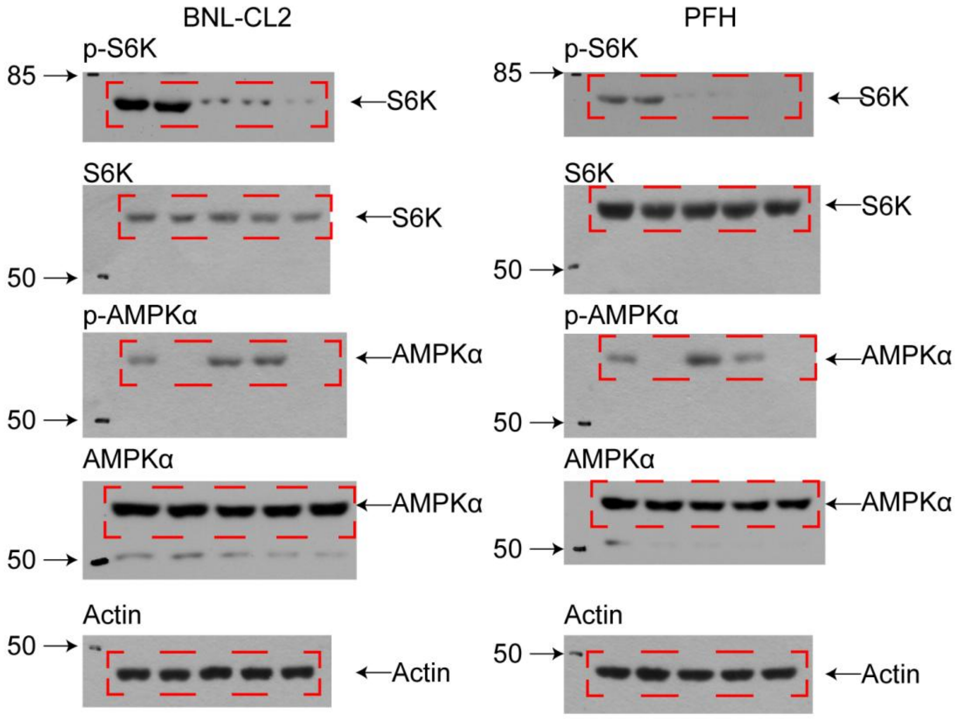
BNL-CL2
PFH
p-S6K
85
S6K
S6K
50
S6K
p-AMPKα
50
AMPKα
AMPKα
50
AMPKα
Actin
50
Actin
p-S6K
85
S6K
S6K
S6K
50
p-AMPKα
AMPKα
50
AMPKα
AMPKα
50
Actin
50
Actin

Fig. 3J

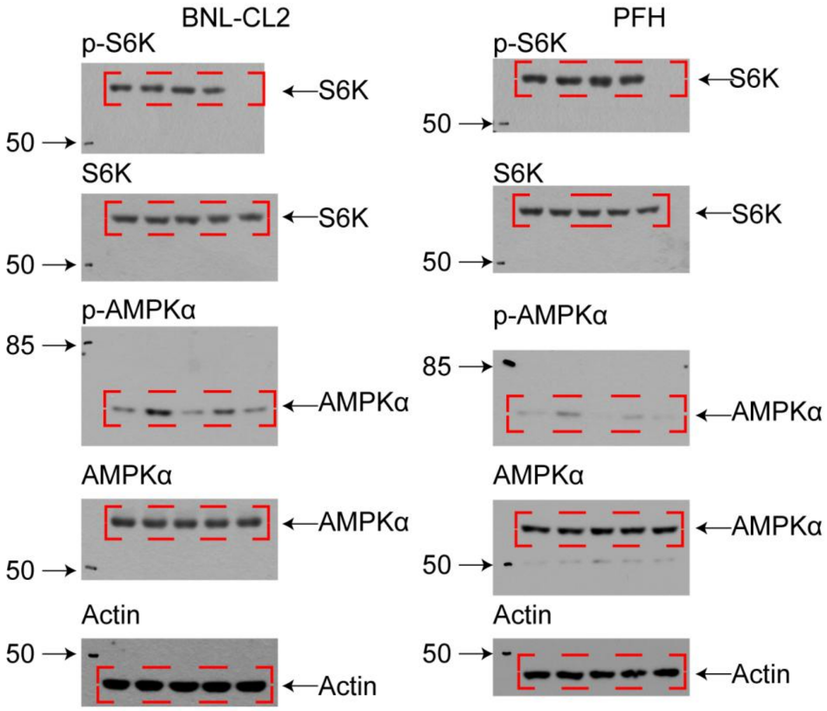

BNL-CL2
p-S6K
S6K
50
S6K
S6K
50
p-AMPKα
85
AMPKα
AMPKα
AMPKα
50
Actin
50
Actin
PFH
p-S6K
S6K
50
S6K
S6K
50
p-AMPKα
85
AMPKα
AMPKα
AMPKα
50
Actin
50
Actin

Fig. 3K

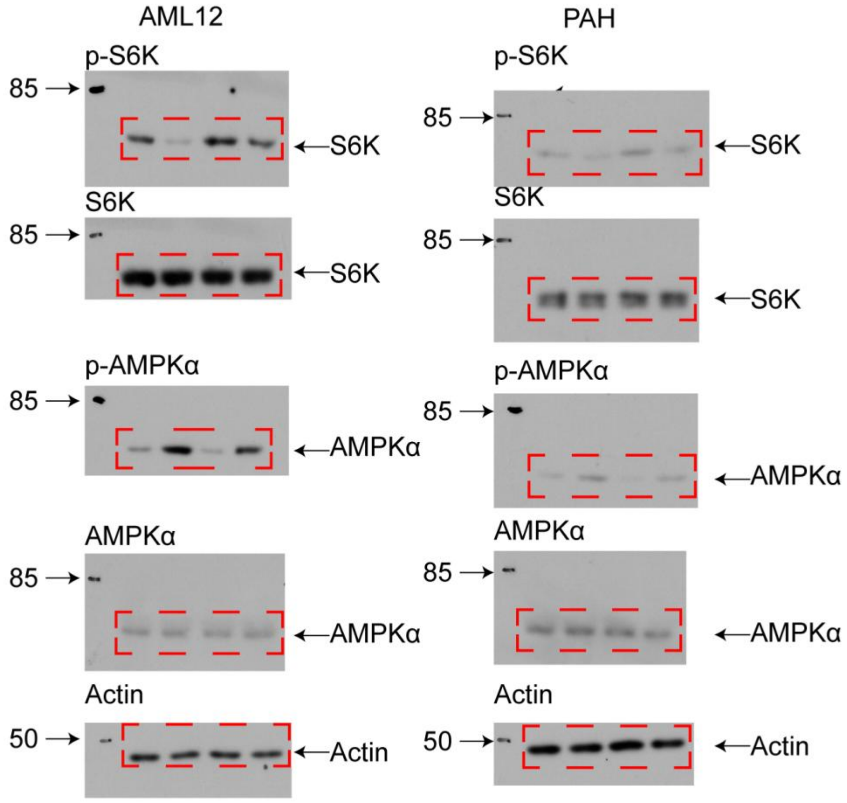
AML12
PAH
p-S6K
85
S6K
85
S6K
p-S6K
85
S6K
S6K
85
S6K
S6K
85
S6K
p-AMPKα
85
AMPKα
p-AMPKα
85
AMPKα
AMPKα
85
AMPKα
AMPKα
85
AMPKα
Actin
50
Actin
Actin
50
Actin

Fig.4A

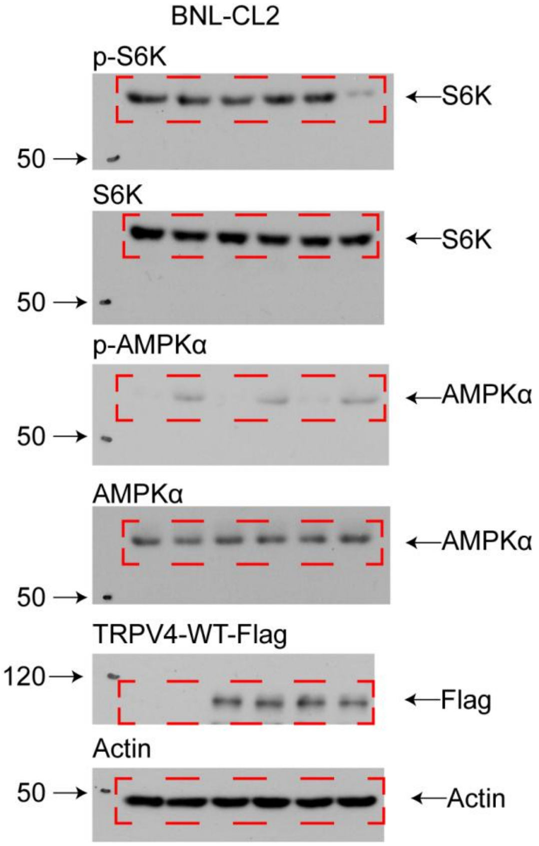
BNL-CL2
p-S6K
S6K
50
S6K
S6K
50
p-AMPKα
AMPKα
50
AMPKα
AMPKα
50
TRPV4-WT-Flag
120
Flag
Actin
50
Actin

Fig. 4F

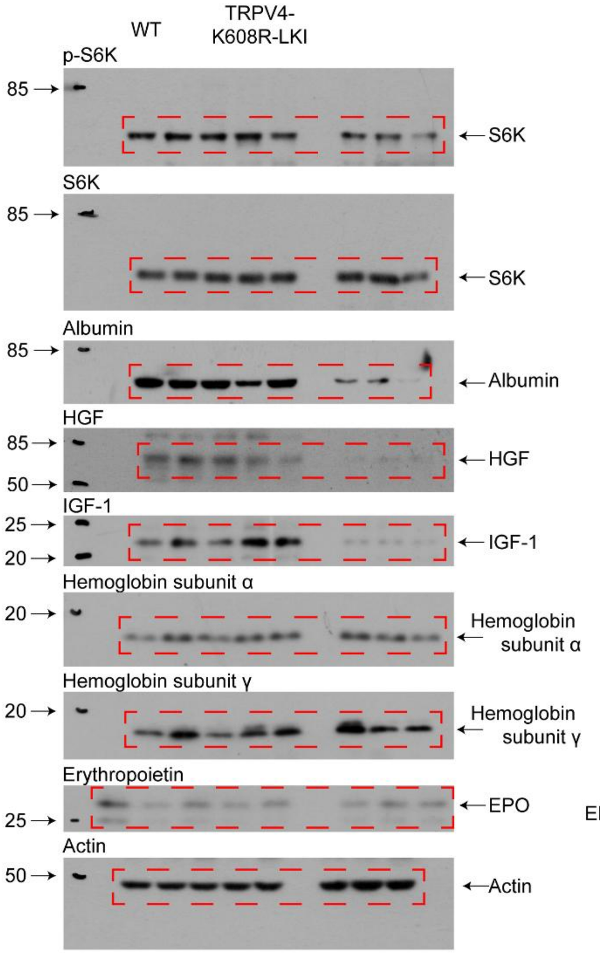
WT
TRPV4-K608R-LKI
p-S6K
85
S6K
S6K
85
S6K
Albumin
85
Albumin
HGF
85
50
HGF
IGF-1
25
20
IGF-1
Hemoglobin subunit α
20
Hemoglobin subunit α
Hemoglobin subunit γ
20
Hemoglobin subunit γ
Erythropoietin
25
EPO
Actin
50
Actin

Fig. 4G

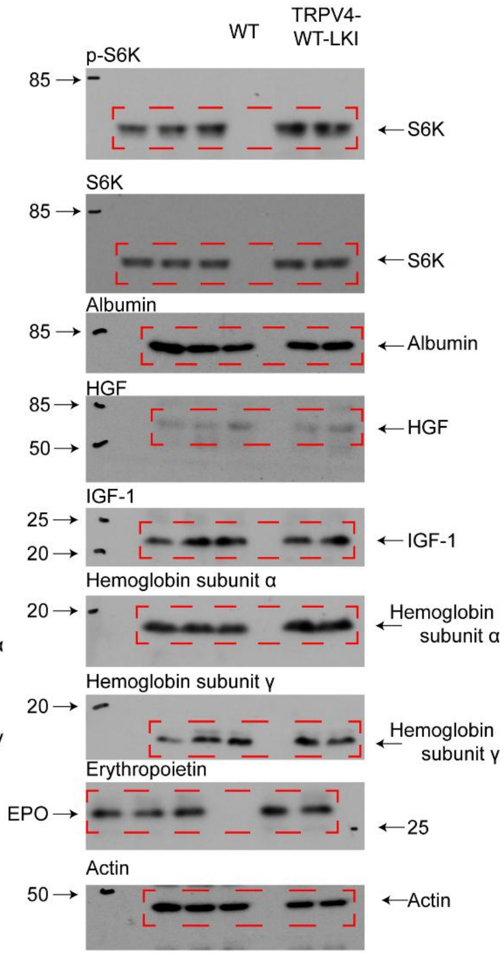
WT
TRPV4-WT-LKI
p-S6K
85
S6K
S6K
85
S6K
Albumin
85
Albumin
HGF
85
50
HGF
IGF-1
25
20
IGF-1
Hemoglobin subunit α
20
Hemoglobin subunit α
Hemoglobin subunit γ
20
Hemoglobin subunit γ
Erythropoietin
EPO
25
Actin
50
Actin

Fig. S1A

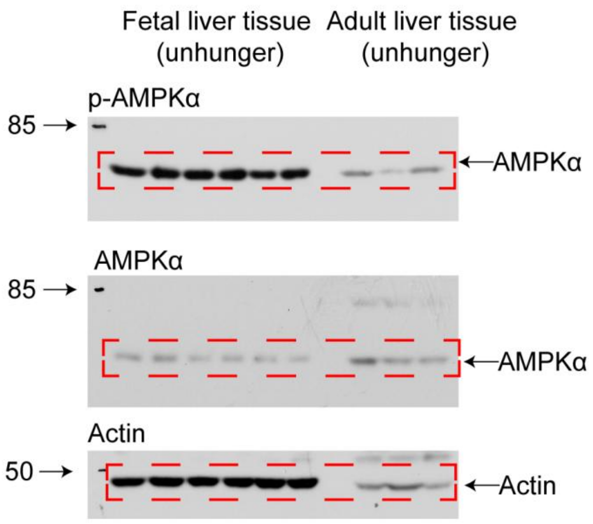


Fig. S1B

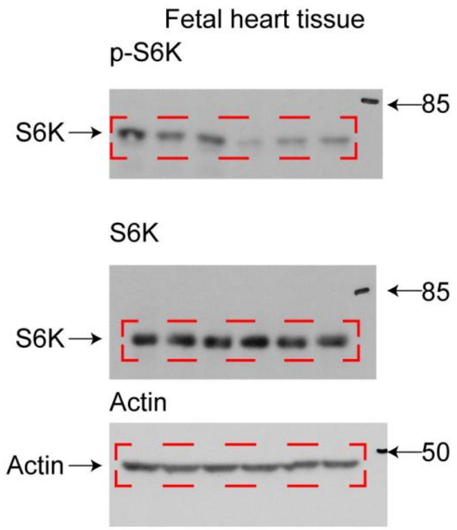

Fig. S1C

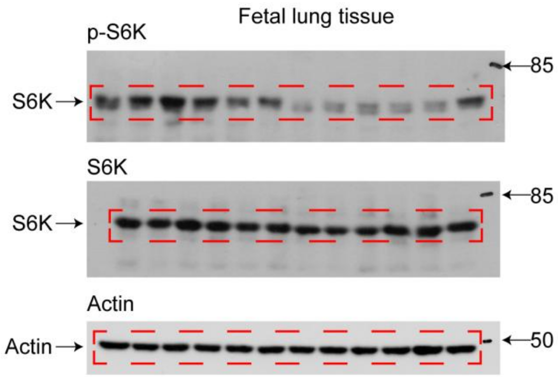


Fig. S1D

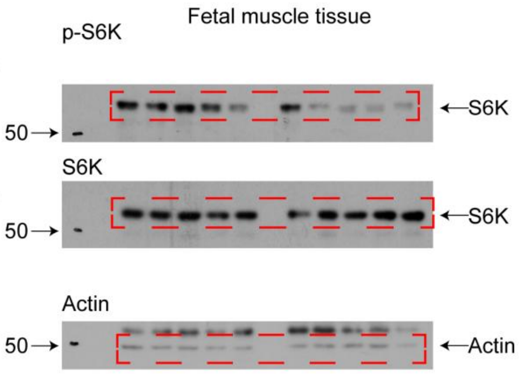

Fig. S1E

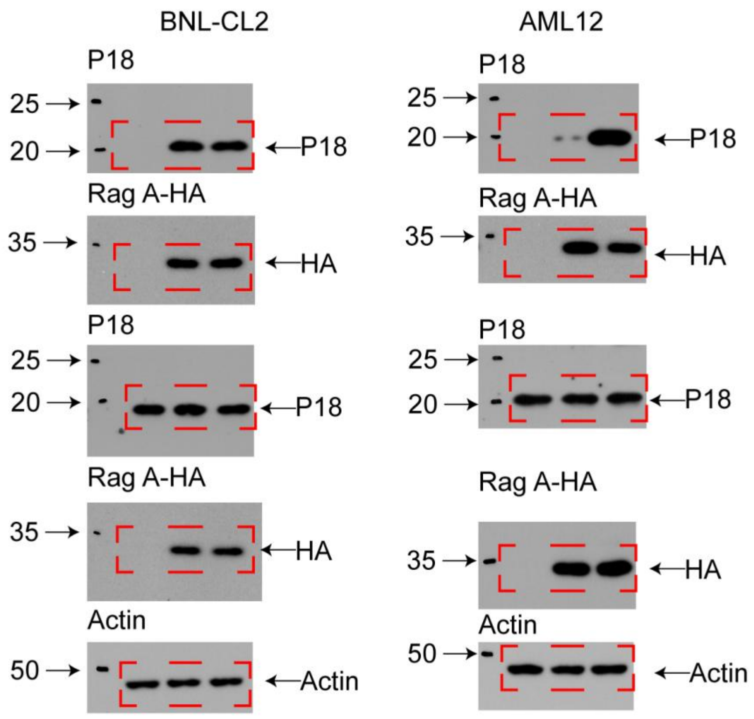
BNL-CL2
AML12
P18
25
20
P18
Rag A-HA
35
HA
P18
25
20
P18
Rag A-HA
35
HA
Actin
50
Actin
P18
25
20
P18
Rag A-HA
35
HA
P18
25
20
P18
Rag A-HA
35
HA
Actin
50
Actin

Fig. S8A

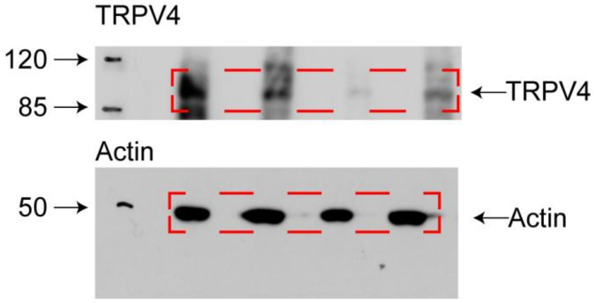

Fig. S9A

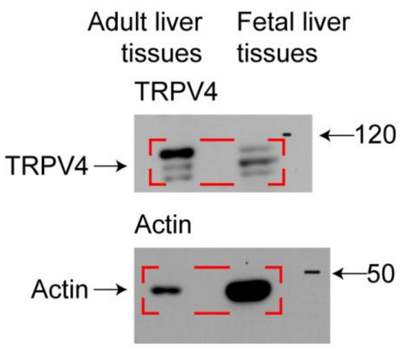


Fig. S9B

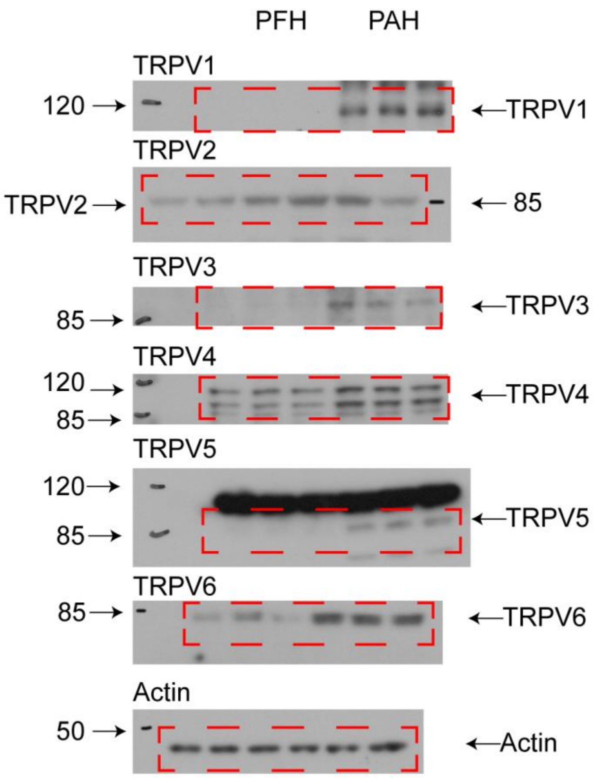

Fig. S10K

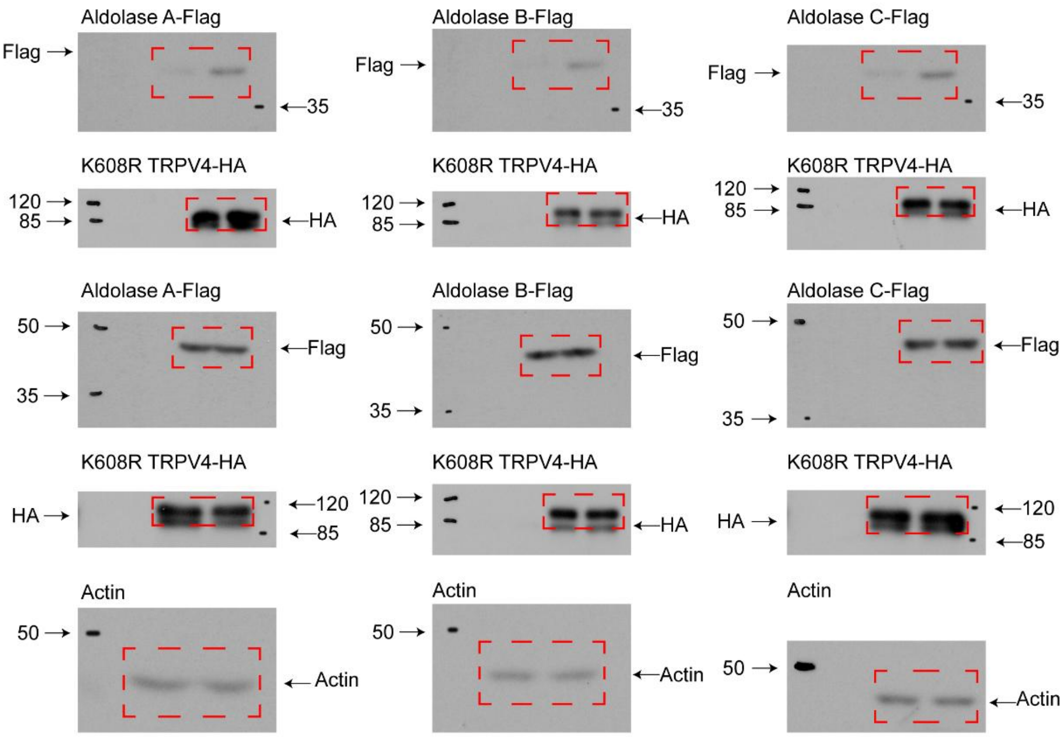
Aldolase A-Flag
Flag
35
Aldolase B-Flag
Flag
35
Aldolase C-Flag
Flag
35
K608R TRPV4-HA
120
85
HA
K608R TRPV4-HA
120
85
HA
K608R TRPV4-HA
120
85
HA
Aldolase A-Flag
50
35
Flag
Aldolase B-Flag
50
35
Flag
Aldolase C-Flag
50
35
Flag
K608R TRPV4-HA
HA
120
85
K608R TRPV4-HA
120
85
HA
K608R TRPV4-HA
HA
120
85
Actin
50
Actin
Actin
50
Actin
Actin
50
Actin

Fig. S12R

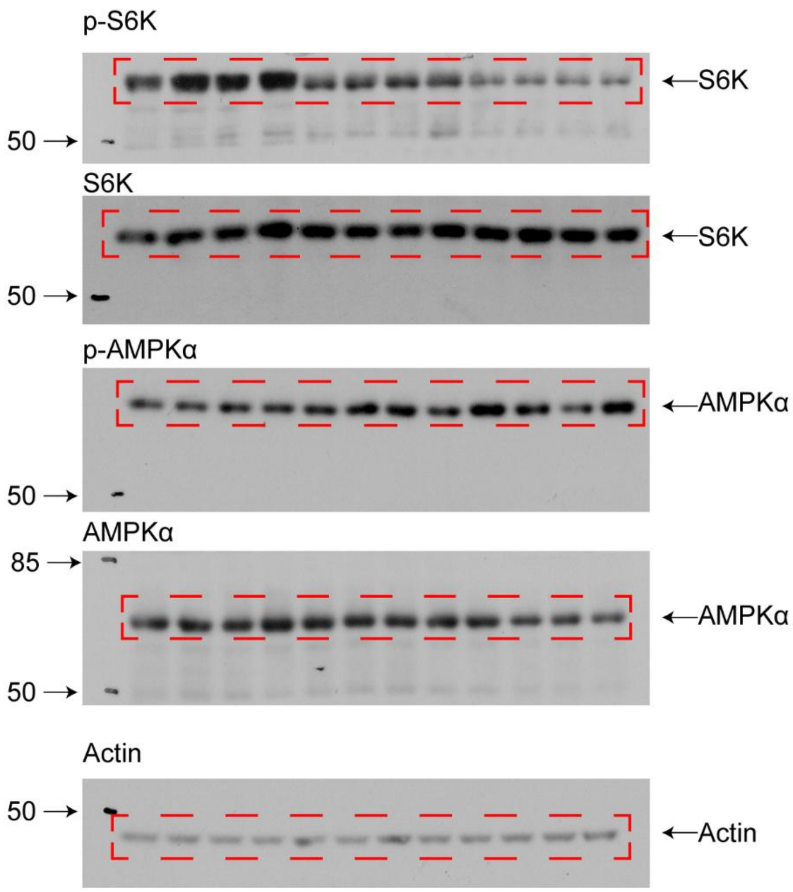
p-S6K
50
S6K
S6K
50
S6K
p-AMPKα
AMPKα
50
AMPKα
85
AMPKα
50
Actin
50
Actin

Fig. S12S

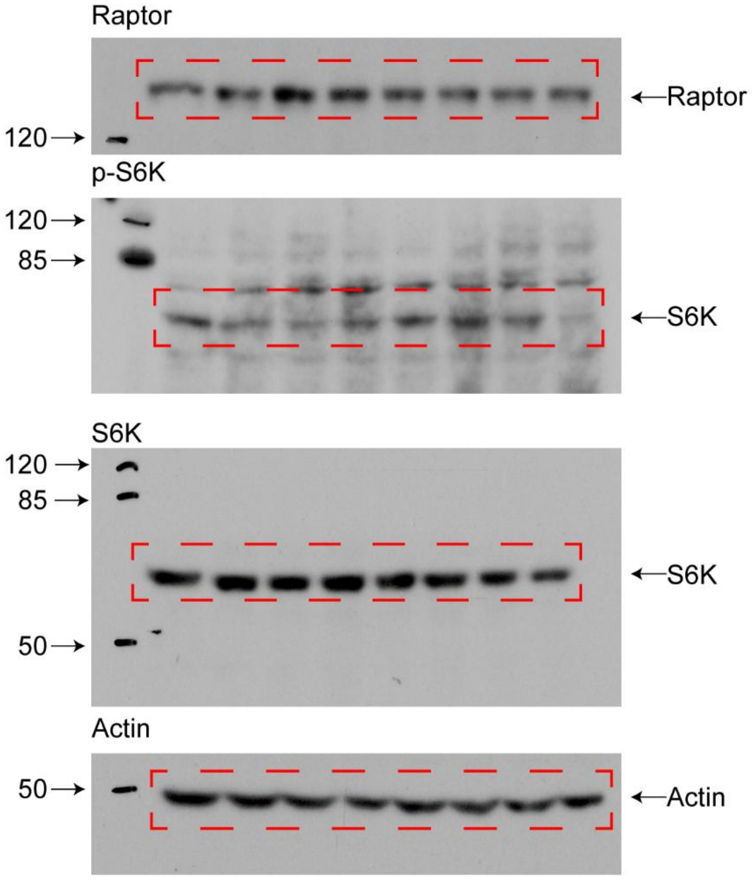

Raptor
←Raptor
120→
p-S6K
120→
85→
←S6K
S6K
120→
85→
←S6K
50→
Actin
50→
←Actin
